## Supplementary material for "Closed-loop phase-locked EEG-tACS enables adaptive control of cortical beta synchrony and motor control": Latex project: 00_Article_Merge.pdf

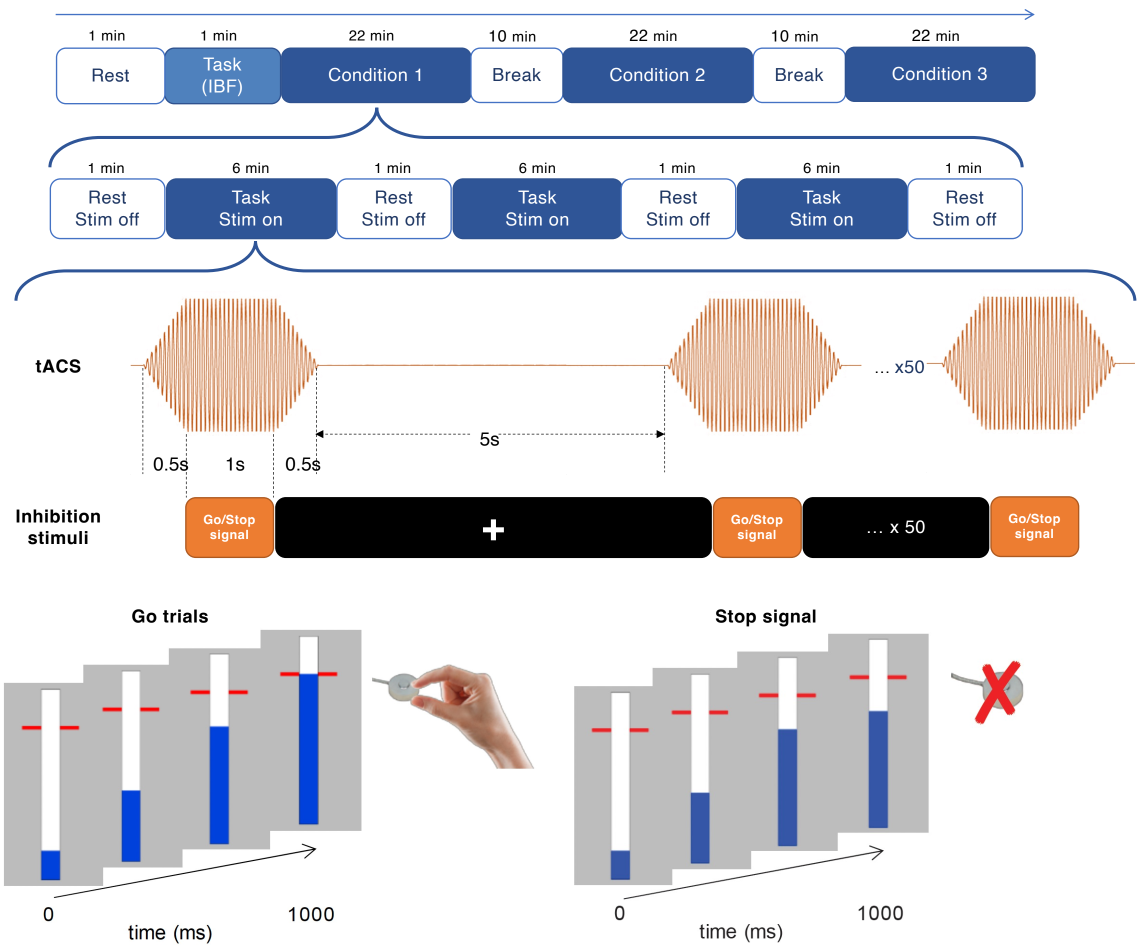


**Figure 1. Experimental Procedure and Stop-Signal Task Design.** Top Panel: The experimental timeline consists of three conditions (in-phase, anti-phase, and sham). Each condition includes three alternating blocks of resting EEG (1 minute) and task EEG (6 minutes) with concurrent tACS stimulation. A 10-minute break separates the conditions.

Middle Panel: Timing of tACS stimulation and stop-signal task within a trial. Stimulation lasts for 2 seconds, with the stop-signal task occurring during the middle 1 second.

Bottom Panel: Stop-signal task. In the majority of trials, participants pinch a force transducer to stop a moving visual indicator as close as possible to the target line. In stop-signal trials, the blue indicator stopped automatically before reaching the target line, serving as a cue for participants to attempt to withhold their response.

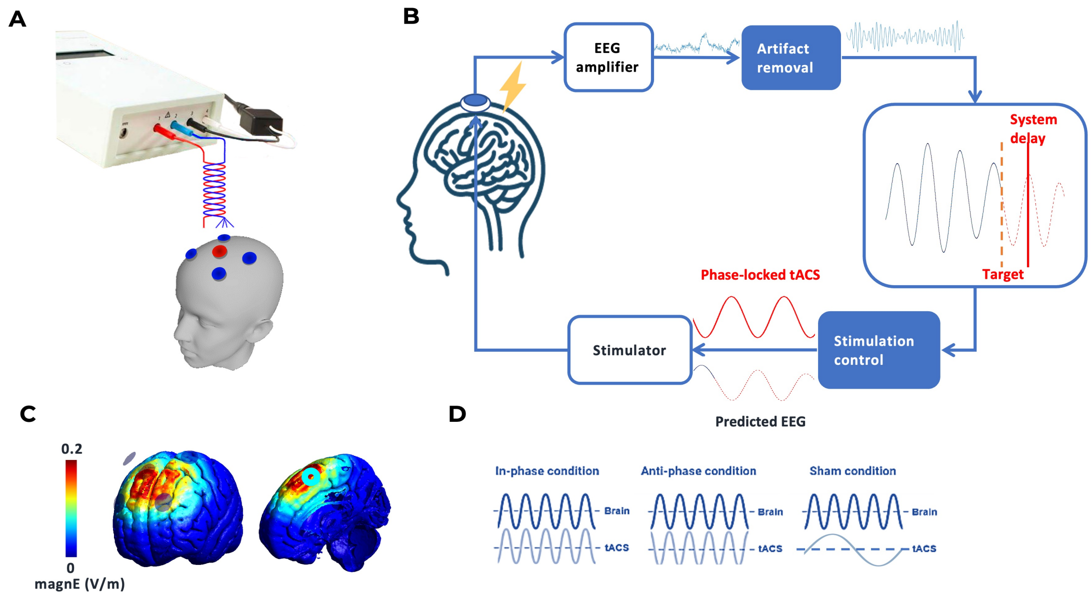


**Figure 2. Closed-Loop tACS-EEG System Setup.** (**A**) 4×1 tACS setup with electrodes placed over the preSMA (centered at FCz) and surrounding regions (F1, F2, C1, C2). (**B**) Real-time closed-loop tACS-EEG system workflow. EEG data from preSMA is processed to predict the phase of beta oscillations, enabling phase-locked tACS delivery synchronized with the stop-signal task. (**C**) Simulated electric field distribution. (**D**) Phase locked tACS for in-phase, anti-phase, and sham conditions.

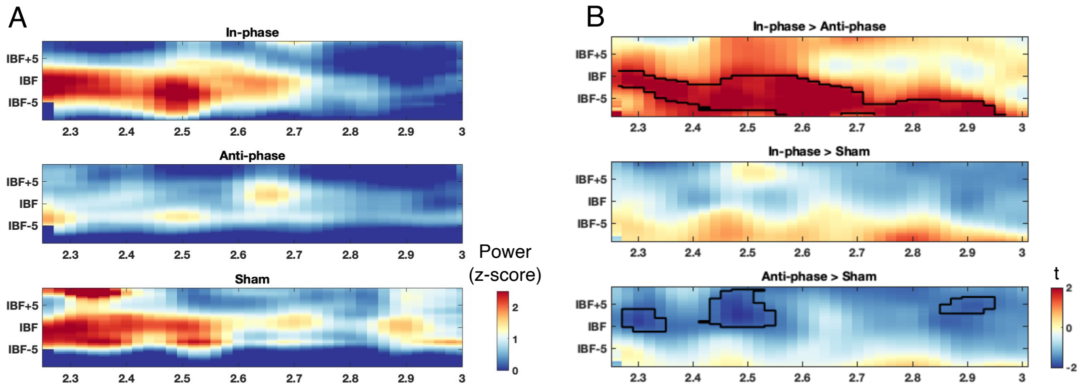


**Figure 3.** Time-frequency representations (x: time [s], y: frequency [Hz]) of tACS aftereffects at FC1/FC2 during the inter-stimulation interval (0.25-1 s post stimulation), aligned to each participant's individual beta frequency (IBF). (**A**) Baseline-corrected power (z-score; baseline 4.5-4.7 s) shown separately for in-phase, anti-phase, and sham. (**B**) Cluster-based paired t-contrasts ($p < 0.05$, TFCE corrected) between conditions: in-phase > anti-phase, in-phase > sham, and anti-phase > sham. Significant clusters are outlined in black.

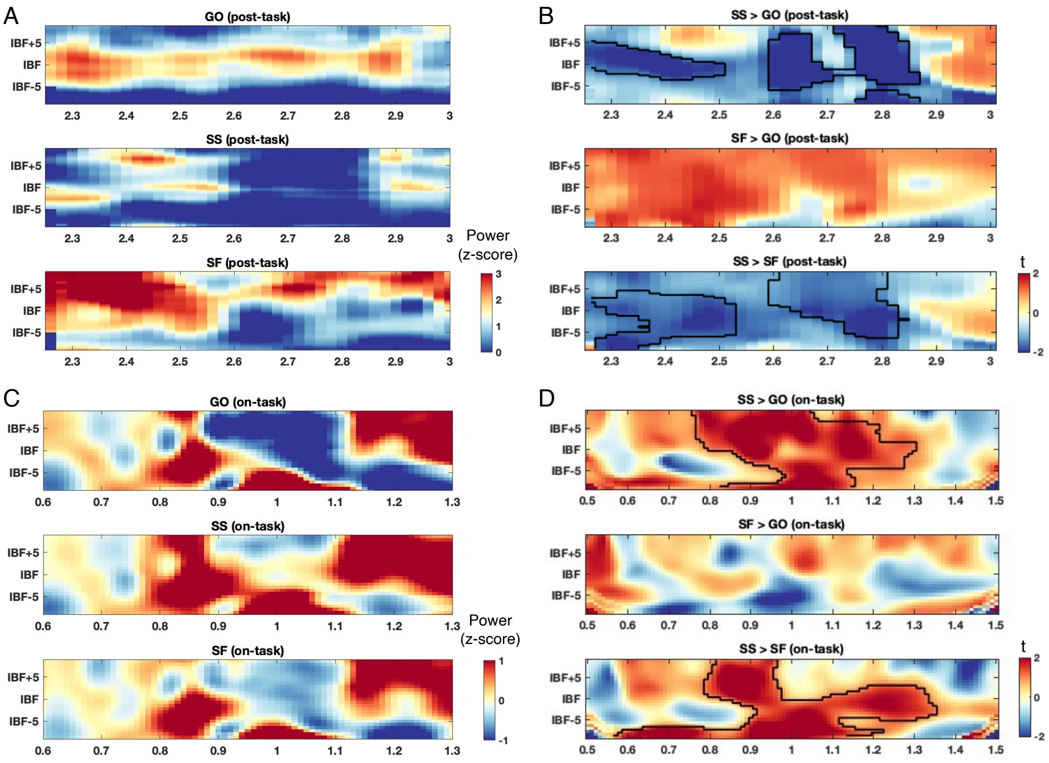


**Figure 4.** Time-frequency representations (x: time [s], y: frequency [Hz]) at FC1/FC2, aligned to each participant's individual beta frequency (IBF). (**A**) Post-task baseline-corrected power (z-score; baseline 4.5-4.7 s) shown separately for go, stop-successful (SS) and stop-failed (SF) trials during the inter-stimulation interval (2.1-3.0 s). (**B**) Cluster-based paired t-contrasts ($p < 0.05$, cluster corrected) of post-task EEG power between trial types (GO > SS, GO > SF, SF > SS). Significant clusters are outlined in black. (**C**) On-task baseline-corrected power (z-score; baseline 0.7-1.1 s, average stop signal presentation time = 1.1) shown separately for go, stop-successful (SS) and stop-failed (SF) trials. (**D**) Cluster-based paired t-contrasts ($p < 0.05$, cluster corrected) of on-task EEG power between trial types (GO > SS, GO > SF, SF > SS). Significant clusters are outlined in black.

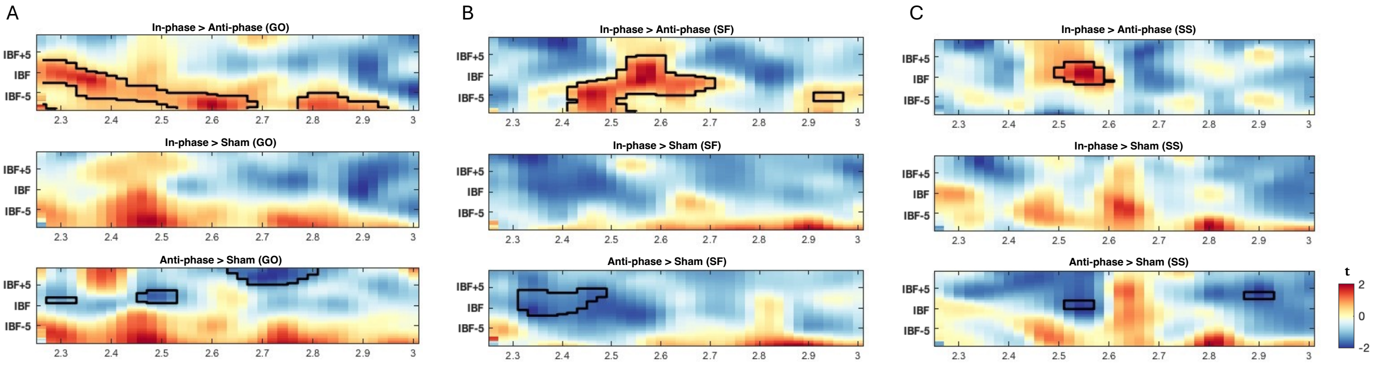


**Figure 5.** Time-frequency representations (x: time [s], y: frequency [Hz]) for cluster-based paired t-statistic ($p < 0.05$, TFCE corrected) between conditions within each trial type. Spectra are at FC1/FC2 during the inter-stimulation interval (2.25-3.0 s), aligned to each participant's individual beta frequency (IBF). Panels depict (**A**) GO, (**B**) SF, and (**C**) SS trials. Rows show condition contrasts: in-phase vs anti-phase (top), in-phase vs sham (middle), and anti-phase vs sham (bottom). Significant clusters are outlined in black ($p < 0.05$).

1 summarizes the posterior means and 95% credible intervals (CIs) for each model parameter across conditions, along with the difference between posterior distributions. Statistical significance is represented by Bayesian p-values, defined as the proportion of the posterior distribution of the latter condition that exceeds $(p > 0.95)$ or falls below $(p < 0.05)$ the former condition. Figure 7 presents the posterior distributions for each population-level parameter, aligning with the statistics in Table 1.

For the 'stop' process, we observed an increase in $\mu_{\text{stop}}$ in the anti-phase (196 ms) compared to the in-phase (191

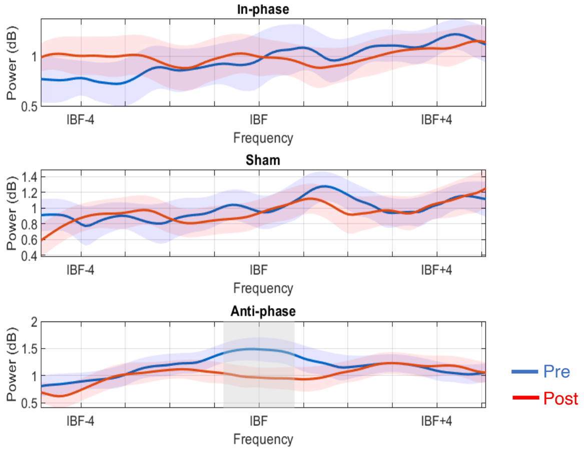


**Figure 6.** Pre- and post-stim EEG power spectra (IBF$\pm$5 Hz) across three conditions. Pre-spectra are in blue, and post-spectra are in red. Shaded regions represent variability ($\pm$SEM), and gray areas indicate frequency ranges with significant pre-post differences from the paired t-test. Anti-phase stimulation resulted in a significant decrease in beta power around IBF$\pm$1 Hz.

ms) condition, suggesting that anti-phase stimulation might slow the stop process. Moreover, $\tau_{\text{stop}}$ was higher in the in-phase (21 ms) condition relative to the anti-phase (12 ms). The overall SSRT remained comparable across conditions, likely due to a trade-off between $\mu_{\text{stop}}$ and $\tau_{\text{stop}}$ components. For the 'go' process, the posterior distribution of $\sigma_{\text{go}}$ is significantly lower in the in-phase condition (33 ms) compared to sham (35 ms). $\tau_{\text{go}}$ was increased in both the in-phase (18 ms) and anti-phase (18 ms) conditions relative to sham (9 ms), while no significant difference was found in $\mu_{\text{go}}$, which was translated into a significantly slower mean GORT $(\mu_{\text{go}} + \tau_{\text{go}})$ in the anti-phase conditions (823 ms) than in sham (815 ms). This pattern suggests that anti-phase tACS did not change the central tendency of the go process, but selectively increased the tail of the finishing time distribution, reflecting a higher frequency of slow responses. In contrast, in-phase tACS appeared to reduce variability of the go finishing time, yet at the same time induced more frequent slow responses.

Finally, we observed significant conditional impacts on the filled-interval illusion, as reflected in the parameters $M$ (shift) and $S$ (spread) of the go finishing distribution on stop-signal trials influenced by the illusion. The $M$ parameter was lower in the in-phase condition (3.6) relative to sham (5.7), suggesting a reduced elongation of the perceived interval following stop signals. In addition, $S$ was significantly lower in the anti-phase condition (1.07) relative to sham (1.14), reflecting reduced variability in interval estimation under anti-phase stimulation.

We did not observe significant differences in PTF and PGF across stimulation conditions. This stable likelihood for both go and stop trigger failures indicates that the observed changes in stop and go distributions are unlikely to stem from attentional fluctuations, but rather reflect changes in inhibitory efficiency caused by the stimulation[51].

**Table 1.** Posterior means and 95% credible intervals (CIs) of the population-level mean parameters across stimulation conditions.

| | Mean (95% CI) | | | In.- Anti. | | In. - Sham | | Anti. - Sham | |
|---|---|---|---|---|---|---|---|---|---|
| Parameter | In-phase | Anti-phase | Sham | Main diff. (95% CI) | p | Main diff. | p | Main diff. | p |
| $P_{TF}$ | 0.009 (-0.014-0.085) | 0.019 (-0.020-0.114) | 0.019 (-0.019-0.118) | 0.011 (-0.078-0.106) | .587 | 0.012 (-0.079-0.111) | .591 | 0.001 (-0.100-0.106) | .505 |
| $P_{GF}$ | 0.028 (0.004-0.069) | 0.026 (0.008-0.054) | 0.025 (0.006-0.057) | -0.003 (-0.049-0.035) | .462 | -0.003 (-0.049-0.037) | .449 | -0.000 (-0.035-0.036) | .492 |
| $\mu_{stop}$ | 191.265 (187.335-195.320) | 196.216 (192.792-199.551) | 192.475 (188.603-196.375) | 4.951 (-0.267-10.061) | **.968*** | 1.210 (-4.304-6.623) | .670 | -3.740 (-8.936-1.489) | .078 |
| $\sigma_{stop}$ | 10.195 (3.446-17.502) | 8.843 (3.036-15.118) | 7.487 (2.510-13.240) | -1.352 (-11.800-8.669) | .402 | -2.708 (-12.659-6.732) | .293 | -1.356 (-10.390-7.455) | .385 |
| $\tau_{stop}$ | 21.437 (13.855-26.133) | 12.542 (4.552-19.677) | 18.898 (7.780-25.881) | -8.894 (-19.079-1.312) | **.043*** | -2.539 (-12.666-9.273) | .292 | 6.355 (-5.295-19.312) | .853 |
| SSRT | 212.701 (204.361-219.072) | 208.500 (199.843-216.505) | 211.349 (199.842-219.484) | -4.202 (-15.843-7.185) | .233 | -1.353 (-12.938-11.521) | .389 | 2.849 (-9.953-16.662) | .664 |
| $\mu_{go}$ | 801.298 (797.328-805.281) | 804.326 (800.563-808.074) | 805.808 (801.384-810.306) | 3.028 (-2.516-8.465) | .861 | 4.510 (-1.460-10.669) | .933 | 1.482 (-4.211-7.341) | .696 |
| $\sigma_{go}$ | 32.929 (31.347-34.512) | 34.470 (32.974-35.967) | 35.397 (33.885-36.881) | 1.541 (-0.663-3.702) | .922 | 2.468 (0.254-4.681) | **.984*** | 0.927 (-1.197-3.059) | .812 |
| $\tau_{go}$ | 18.039 (15.846-20.121) | 18.737 (15.675-21.460) | 9.128 (3.424-13.828) | 0.698 (-2.730-4.534) | .648 | -8.911 (-14.085–2.457) | **.004**** | -9.609 (-15.406–2.900) | **.002**** |
| GORT | 819.337 (814.760-823.786) | 823.064 (818.282-827.737) | 814.809 (807.497-821.470) | 3.727 (-2.845-10.374) | .871 | -4.527 (-12.603-4.325) | .145 | -8.254 (-16.579-0.609) | **.034*** |
| M | 3.565 (1.697-5.446) | 5.735 (3.682-7.800) | 5.868 (3.747-7.869) | 2.170 (-0.647-4.980) | .936 | 2.303 (-0.429-5.144) | **.952*** | 0.133 (-2.831-3.083) | .538 |
| S | 1.119 (1.065-1.174) | 1.066 (1.019-1.115) | 1.139 (1.094-1.184) | -0.053 (-0.127-0.019) | .076 | 0.020 (-0.052-0.089) | .715 | 0.073 (0.006-0.139) | **.984*** |

Values are reported for in-phase, anti-phase, and sham conditions. Pairwise contrasts are reported alongside the corresponding ‘Main diff’ (i.e., the value computed by subtracting the posterior samples of the latter condition from the former condition, such that positive values indicate higher estimates in the first condition) and Bayesian p-values (i.e., posterior probabilities that the parameter value in the second condition exceeds that in the first) indicated with $p < .05$ (*) and $p < .01$ (**).

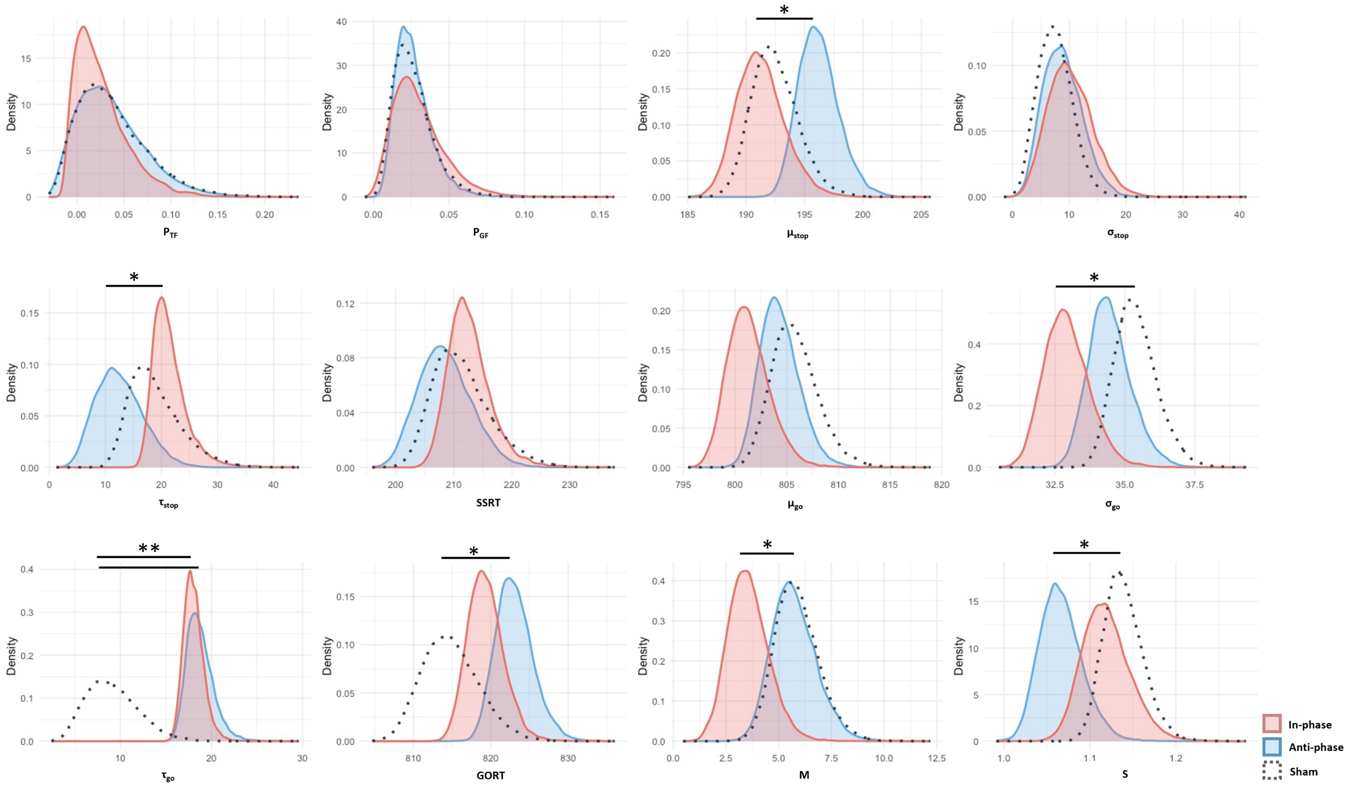


**Figure 7.** Posterior distributions of population-level parameters estimated by the BEESTS-CV-DMC model across in-phase (red), anti-phase (blue), and sham (dot) conditions, aligning with the statistics in Table 1. Asterisks indicate that the Bayesian $p$-values $> 0.95$ or $< 0.05$.

### Force results

A linear mixed-effects model was conducted to examine the effects of stimulation condition (in-phase, anti-phase, and sham) on force outcomes. The analysis revealed a statistically significant main effect of stimulation on peak force rate during GO trials ($F(2, 10034) = 4.68, p < 0.001$) and on response onset time during stop-signal (SS) trials ($F(2, 2296) = 3.18, p = 0.04$; Table 2). Post-hoc pairwise comparisons (Tukey-HSD) revealed that in-phase elicited significantly lower peak force rate during GO trials compared with both anti-phase ($z = -2.45, p = 0.04$) and sham ($z = -2.8, p = 0.01$; $SE = 2.32$). For SS trials, onset time was significantly longer under in-phase than sham stimulation (in-phase - sham: $z = 2.4, p = 0.04$, $SE = 2.87$).

**Table 2.** Effect of stimulation on force measures.

| Trial type | Measures | Summary | | | LME statistic |
|---|---|---|---|---|---|
| | | Inphase | Antiphase | Sham | Main effect |
| GO | Peak force (N) | 30.48±9.09 | 30.59±9.52 | 30.37±9.08 | F(2,10042)=2.5, p=.08 |
| | Peak force rate (N/s) | 355.57±125.33 | 361.6±128.85 | 362±156.87 | **F(2,10034)=4.68, p<.001*** |
| | Onset time (ms) | 703.78±51.26 | 704.72±55.06 | 703.45±53.6 | F(2,9984)=0.59, p=.56 |
| SF | Peak force (N) | 25.10±7.61 | 25.15±8.01 | 25.12±7.52 | F(2,2362)=0.002, p=1 |
| | Peak force rate (N/s) | 329.32±120.86 | 330.11±117.46 | 332.88±118.8 | F(2,2352)=0.89, p=.41 |
| | Onset time (ms) | 703.78±51.26 | 704.72±55.06 | 703.45±53.60 | F(2,9984)=0.59, p=.56 |
| SS | Peak force (N) | 6.80±6.37 | 7.24±7.15 | 6.93±6.44 | F(2,2566)=1.27, p=.28 |
| | Peak force rate (N/s) | 103.85±98.09 | 107.83±101.12 | 115.85±257.15 | F(2,2744)=1.45, p=.23 |
| | Onset time (ms) | 711.5±64.05 | 705.45±66.51 | 704.68±62.16 | **F(2,2296)=3.18, p=0.04*** |

Linear mixed-effects (LME) model results for peak force, peak force rate, and onset time across in-phase, anti-phase and sham stimulation conditions during go (GO), stop-failed (SF), and stop-successful (SS) trials. Asterisks (*) denote statistically significant effects at $p < 0.05$.

These findings suggest that in-phase tACS attenuates motor output and enhances inhibitory control, as reflected in reduced peak force rate during voluntary responses and delayed response onset during inhibition.

#### Correlations between beta power and behavioral measures

***Plausible correlation analysis (SSRT-beta power).*** To assess the relationship between post-stimulation beta power and the SSRT posterior estimates from the BEESTS model, we employed a plausible correlation analysis using Bayesian hierarchical modeling that accounts for individual-level uncertainty and provides a population-level distribution of correlation coefficients. The result revealed a reliable negative association in both sham and in-phase stimulation between SSRT and post-stimulation beta power (Figure 8). Specifically, participants eliciting stronger post-stimulation beta power tended to exhibit shorter SSRTs (sham: $r = -0.221$, 95% CI $[-0.461, -0.040]$, $p = 0.048$; In-phase: $r = -0.265$, 95% CI $[-0.505, 0.003]$, $p = 0.027$). In contrast, no significant correlations were observed in the anti-phase condition.

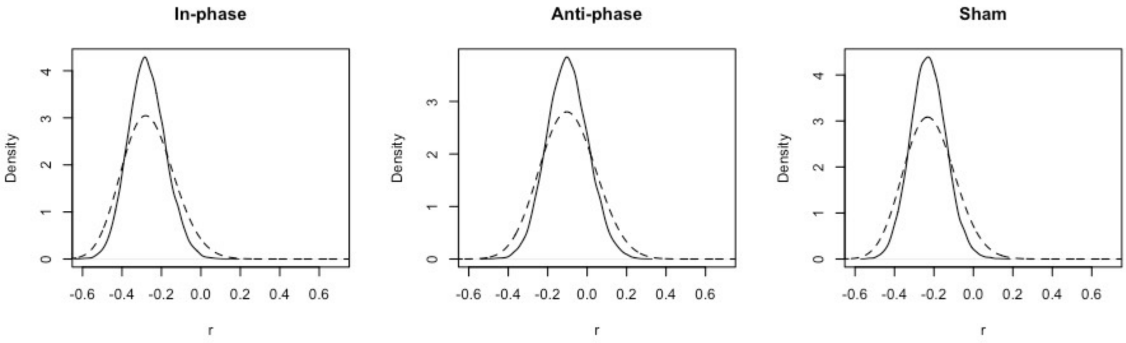


**Figure 8.** Posterior density estimates of plausible correlations (r) between IBF±2 EEG power and hierarchical SSRT. Solid lines represent the individual-level plausible correlation distributions, and dashed lines represent the population-level posterior distributions obtained using a uniform prior ($\kappa = 1$).

These correlations suggest that enhanced beta synchronization during the post-stimulation phase is linked to faster stopping processes. While anti-phase tACS may disrupt the natural relation between inhibitory efficiency and post-movement cortical beta dynamics.

#### Phase-dependency of inhibition performance

To examine whether in-phase versus anti-phase tACS influences the relationship between stop-signal presentation phase and inhibitory control, we conducted paired t-tests comparing peak force on stop trials (SS and SF) between stop signals delivered at the down versus up phases of the online tACS waveform. Building on our previous finding that inhibition success depends on the phase of the endogenous beta cycle at which the stop signal is processed[47], this analysis tested whether externally aligning or misaligning tACS with that rhythm modulates the phase-specific effect.

The analysis revealed a significant phase-dependent modulation of peak force during SS trials exclusively under in-phase tACS: peak force was significantly lower when the stop signal occurred at the down phase compared to the up phase of the stimulation waveform ($t(32) = -3.04$; $p = 0.005$) (Figure 9). By contrast, anti-phase tACS did not produce any phase-dependent effects on peak force in SS trials ($t(32) = -0.16$, $p = 0.88$), nor were significant differences observed in SF trials under either stimulation condition (In-phase: $t(32) = -0.79$, $p = 0.43$; Anti-phase: $t(32) = 0.03$, $p = 0.97$). The in-phase results closely replicate our prior finding[47] that inhibition is strongest when the stop signal coincides with the preferred phase of the beta oscillation, suggesting that in-phase tACS preserves this stable phase relationship. In contrast, anti-phase stimulation appears to disrupt this alignment, attenuating the phase-dependent modulation of inhibitory control.

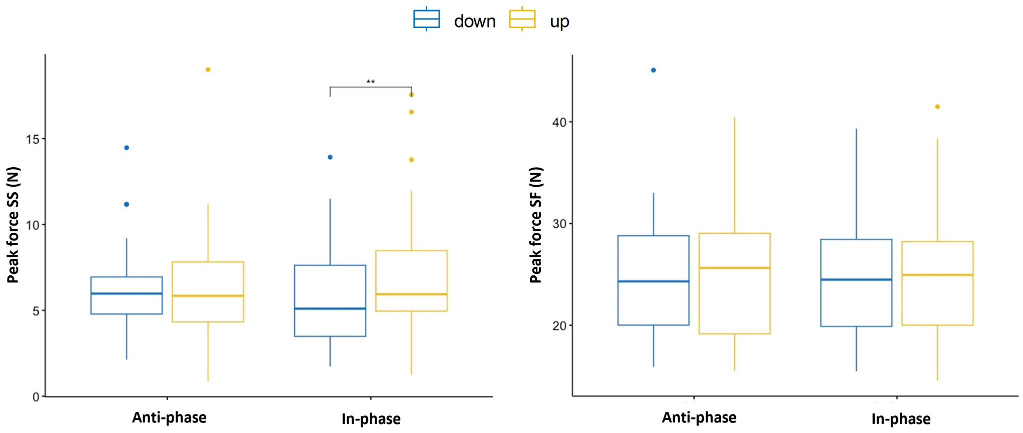


**Figure 9.** Box plots of peak force (N) at the up and down phases on SS and SF trials under in- and anti-phase tACS stimulation. Down and up phases refer to the tACS waveform at the time of the stop signal presentation, respectively. The double asterisk (**) indicates a significant difference between up and down stop signal phases ($p = 0.005$).

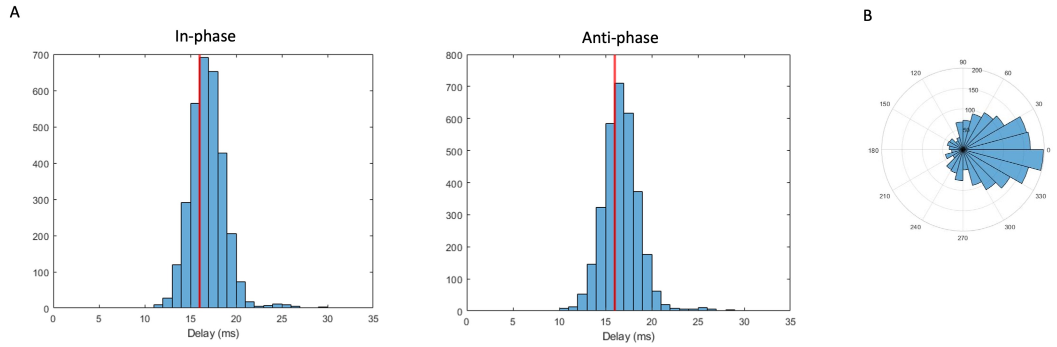


**Figure S1. Closed-loop system evaluation.** (**A**) Distribution of system delays measured over 10,000 trials for in-phase and anti-phase tACS. Median delay (red line) was 16 ms for both (**B**) Polar histogram of phase prediction accuracy, showing the distribution of phase differences between actual and targeted EEG phases (accounting for 16 ms delay), demonstrating precise phase-locking of the closed-loop tACS system.

### Supplementary 2. Self-report discomfort scores

**Table S1.** Mean $\pm$ standard deviation of self-reported discomfort scores (range: 0–100) after each stimulation condition.

| Stimulation condition | Discomfort score (0-100) | Main effect of stimulation |
|---|---|---|
| In-phase | 35.8±22.64 | |
| Anti-phase | 38.74±20.76 | F(2, 68)=16.85, p<0.001* |
| Sham | 20.43±21.66 | |

A repeated measures ANOVA revealed a significant main effect of stimulation condition on perceived discomfort $(F(2, 68) = 16.85, p < 0.001)$.

### Supplementary 3. Context violation and non-parametric integration estimates comparisons

Non-parametric estimated behavioral metrics of stop-signal performance are summarized in Table S3. SSRT was estimated using the integration method with replacement of go omissions[28]. The staircase algorithm successfully

**Table S2.** Tukey's HSD post-hoc comparisons of discomfort scores across stimulation conditions.

| group1 | group2 | Mean diff | p | lower | upper |
|---|---|---|---|---|---|
| Anti-phase | In-phase | 2.9429 | 0.8378 | -9.3944 | 15.2801 |
| Anti-phase | Sham | -15.3714 | 0.0105* | -27.7087 | -3.0342 |
| In-phase | Sham | -18.3143 | 0.0018* | -30.6515 | -5.977 |

Asterisks denote significance ($p < .05$). Sham was rated significantly lower in discomfort than both In-phase and Anti-phase, while no significant difference was found between In-phase and Anti-phase.

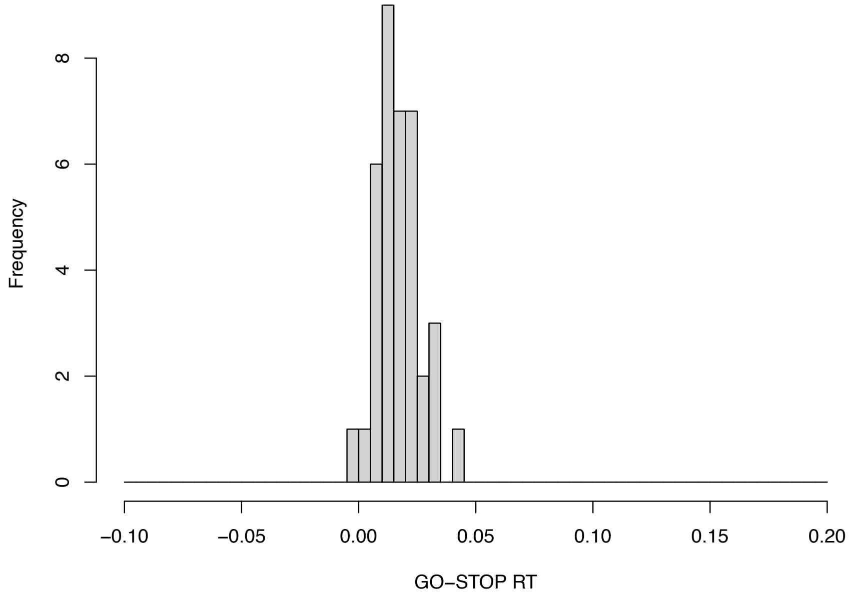


**Figure S2. Assessment of context independence.** GO-STOP RT: Go reaction time – Failed stop reaction time. The dataset did not show a severe violation of independence.

maintained stopping accuracy near the target 50%, with no significant main effect (Friedman test, $\chi^2(2) = 2.9$, $p = 0.23$). The GORT differed significantly across conditions (ANOVA, $F(2,68) = 4.04$, $p = 0.022$), with follow-up comparisons revealing faster responses in the sham vs. anti-phase condition ($p = 0.03$) and slower responses in the in-phase vs. sham ($p = 0.01$). No significant main effects were observed for SSRT or stop-signal delay (SSD), with respective ANOVA results of $F(2,68) = 0.14$ ($p = 0.87$) and $F(2,68) = 1.27$ ($p = 0.29$). Pairwise comparisons for these measures did not reveal any significant differences.

**Table S3.** Summary of descriptive statistics, main effects, and post hoc comparisons across tACS phase conditions for non-parametric estimates.

| | **Mean (SD)** | | | **Main Effect (F/χ², p)** | **In.- Anti.** | | **In. – Sham** | | **Anti. - Sham** | |
|---|---|---|---|---|---|---|---|---|---|---|
| **Parameter** | **In-phase** | **Anti-phase** | **Sham** | | **Main diff.** | **p** | **Main diff.** | **p** | **Main diff.** | **p** |
| P(Inhibition) † | 54.29 (2.12) | 53.94 (1.91) | 53.31 (3.82) | χ²(2)=2.9, p=.23 | 144.50 | .28 | 105.00 | .05 | -129.00 | .54 |
| RTGO (ms) | 818.20 (12.68) | 822.50 (13.11) | 818.11 (13.62) | **F(2,68)=4.04, p=.022** | **-2.71** | **.01** | 0.05 | .96 | **2.30** | **.03** |
| SSRT (ms) | 208.73 (16.94) | 209.46 (13.54) | 209.61 (15.43) | F(2,68)=0.14, p=.87 | -0.39 | .70 | -0.52 | .61 | -0.08 | .94 |
| SFRT (ms) | 801.17 (16.41) | 805.89 (15.44) | 800.87 (17.52) | F(2,68)=2.31, p=.11 | -1.82 | .08 | 0.10 | .92 | **2.19** | **.04** |
| SSD (ms) | 604.87 (17.70) | 608.54 (14.51) | 606.03 (19.01) | F(2,68)=1.27, p=.29 | -1.43 | .16 | -0.54 | .59 | 1.08 | .29 |
| P(Go errors) | 0 | 0 | 0 | NA | NA | NA | NA | NA | NA | NA |
| P(Go omissions) | 2.49 (3.02) | 2.40 (2.53) | 2.34 (3.16) | χ²(2)=0.48, p=.79 | 201.00 | .78 | 149.00 | .99 | -141.50 | .82 |

Main effects and post-hoc results were tested using repeated measures ANOVA and paired t-tests. No corrections were applied to $p$ values.
† Friedman test with Wilcoxon signed-rank tests was conducted for non-normal distributed data.
SSRT=Stop signal reaction time, estimated using the integration method with replacement of go omissions[28]. RTGO=mean reaction time of go trials; SSRT=Stop-signal reaction time; RTSF=mean reaction time of Stop-failed trials; SSD=Stop-signal delay.

### Supplementary 4. BEESTS-CV model prior distributions

The prior distributions:

$$\begin{aligned}
\mu_{go} &\sim Normal(0.5, 0.50)[0, 1],\\
\mu_{stop} &\sim Normal(0.2, 0.50)[0, 1],\\
\sigma_{go}, \sigma_{stop} &\sim Normal(0.03, 0.35)[0, 1],\\
\tau_{go}, \tau_{stop} &\sim Normal(0.03, 0.35)[0, 1],\\
\phi^{-1}(P_{TF}), \phi^{-1}(P_{GF}) &\sim Normal(-1.28, 3),\\
M &\sim Normal(0, 3),\\
S &\sim Normal(1, 3)[0, \infty].
\end{aligned} \tag{S1}$$

The hyper-prior location parameter distributions:

$$\begin{aligned}
\mu_{\mu_{go}} &\sim Normal(0.5, 0.5)[0, 1],\\
\mu_{\mu_{stop}} &\sim Normal(0.2, 0.5)[0, 1],\\
\mu_{\sigma_{go}}, \mu_{\sigma_{stop}} &\sim Normal(0.04, 0.35)[0, \infty],\\
\mu_{\tau_{go}}, \mu_{\tau_{stop}} &\sim Normal(0.04, 0.35)[0, \infty],\\
\mu_{\phi^{-1}(P_{TF})}, \mu_{\phi^{-1}(P_{GF})} &\sim Normal(-1.28, 3)[-\infty, \infty],\\
\mu_M &\sim Normal(0, 3)[-\infty, \infty],\\
\mu_S &\sim Normal(0, 1)[0, \infty].
\end{aligned} \tag{S2}$$

The hyper-prior scale parameters:

$$\sigma_p \sim Exponential(1) \tag{S3}$$

For each parameter $p \in \{\mu_{go}, \mu_{stop}, \sigma_{go}, \sigma_{stop}, \tau_{go}, \tau_{stop}, P_{TF}, P_{GF}, M, S\}$.

### Supplementary 5. BEESTS-CV model fitting and context independence

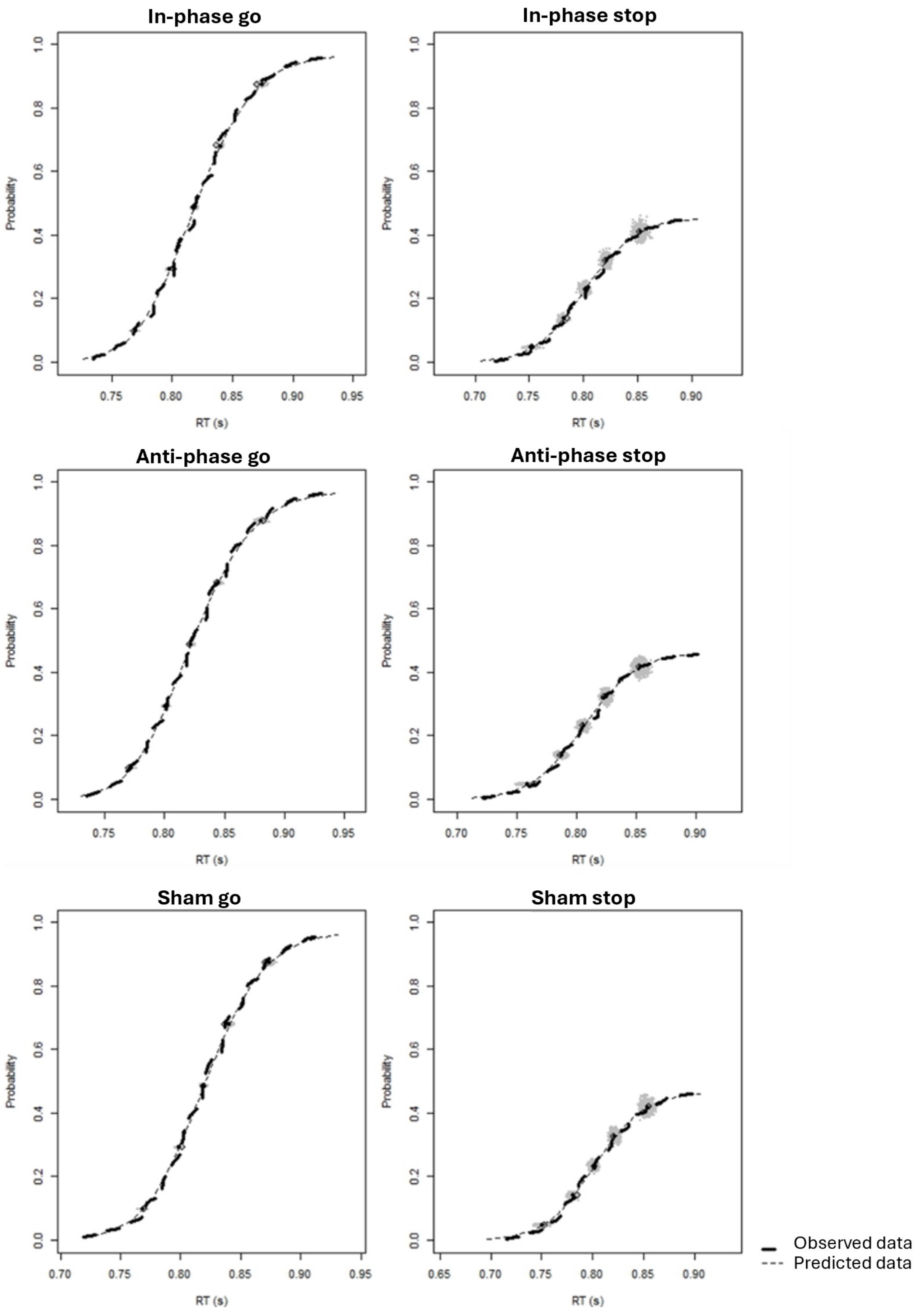


**Figure S3.** Averaged observed and predicted cumulative distribution functions of reaction time (RTs) for go signal (left panels) and stop signals (right panels) collapsed (due to the SSDs determined by a staircase algorithm) across SSDs in three conditions. The thick dashed lines show the observed data. The grey dots show the prediction samples from the joint posterior distribution of the model parameters. The thin dashed lines show the average of the predictions.

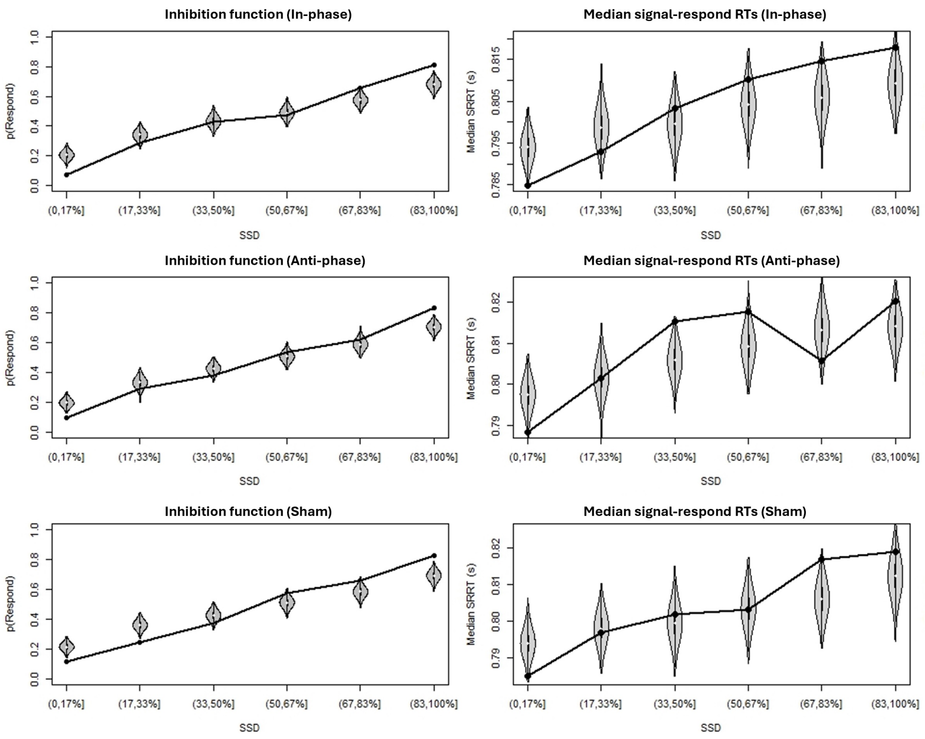


**Figure S4.** Observed vs predicted inhibition functions (left panels) and median signal-respond RT as a function of stop-signal delay (SSD; right panels) for three stimulation conditions. Black bullets show the observed data. Grey violin plots show the distribution of the predictions of the BEESTS-CV model.

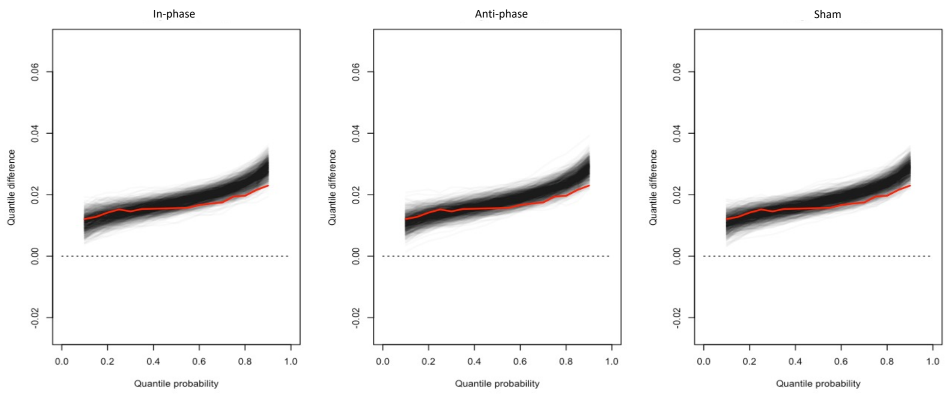


**Figure S5.** Average delta functions in three conditions: in-, anti-phase and sham. The red line shows the observed delta functions averaged across participants. The black lines show 1000 posterior predictive delta functions averaged across participants obtained from fitting the data with the BEESTS-CV model assuming context independence; the darker the area, the denser the predicted delta function.
