## Supplementary material for "Closed-loop phase-locked EEG-tACS enables adaptive control of cortical beta synchrony and motor control": Latex project: 00_Article_Merge.tex

% !TEX root = 00_Article_Merge.tex
%% select between bioRxiv style or submission style by commenting either line
%\documentclass[towcolumn]{bioRxiv}
\documentclass[submit]{bioRxiv}
\renewcommand\familydefault{\sfdefault}
\usepackage{lipsum}
\usepackage{wasysym}
\usepackage{hyperref}
\usepackage{placeins}
\usepackage{lscape}
\usepackage[most]{tcolorbox}
\usepackage{wasysym}
\usepackage{textgreek}
\usepackage{float}

%% setup specific to this preprint -- add or remove as you need.
\hyphenation{empire} % prevents word "empire" being split across lines
\sisetup{range-units = single,
        separate-uncertainty = true,
        multi-part-units = single,
        range-phrase = --} % units setup
\usepackage{multirow} % allows merging of cells in tables
\newcommand{\ismergeddocument}{}

\begin{document}
%% comment out any files that you don't want to include as sections
\include{01_Article_MainText}
\include{02_Article_Supplementary}
\section*{References}
\bibliography{refs.bib}

\end{document}
