## Supplementary material for "Closed-loop phase-locked EEG-tACS enables adaptive control of cortical beta synchrony and motor control": Latex project: 01_Article_MainText.tex

% !TEX root = 00_Article_Merge.tex
\leadauthor{Fang}

\title{Closed-loop phase-locked EEG-tACS enables adaptive control of cortical beta synchrony and motor control}
\shorttitle{Closed-loop anti-phase tACS over preSMA\\desynchronizes beta}

\author[1,2]{Zhou Fang \orcidlink{0009-0002-8871-6419}}
\author[1,2,3]{Alexander Sack \orcidlink {0000-0002-1471-0885}}
\author[1,2,\Letter]{Inge Leunissen \orcidlink {0000-0002-3977-3620}}

\affil[1]{Faculty of Psychology and Neuroscience, Maastricht University, Maastricht, the Netherlands}
\affil[2]{Maastricht Brain Imaging Centre (MBIC), Maastricht University, Maastricht, the Netherlands}
\affil[3]{Faculty of Health, Medicine and Life Sciences, Maastricht University, Maastricht, the Netherlands}
\date{}

\maketitle

\begin{abstract}

Neural oscillations dynamically coordinate communication across distributed brain networks, and their pathological synchronization is a common feature of neurological and psychiatric disease. In Parkinson's disease, excessive beta-band synchronization within cortico-basal ganglia-thalamo-cortical loops is closely linked to motor impairment, highlighting the need for neuromodulation strategies that can selectively and adaptively reshape this brain synchrony online. Transcranial alternating current stimulation (tACS) offers a non-invasive approach to modify endogenous oscillatory activity, yet it is typically delivered in an open-loop manner, limiting control over the timing, direction, and functional network consequences of oscillatory modulation. 

Here, we developed an online adaptive EEG-guided closed-loop system that continuously tracked the instantaneous phase of ongoing cortical oscillations and delivered stimulation either aligned (in-phase) or 180 degrees opposed (anti-phase) to the endogenous beta rhythm. Thirty-eight right-handed healthy participants performed a stop-signal task while receiving adaptive closed-loop $4 \times 1$ HD-tACS over pre-supplementary motor cortex (preSMA) targeting beta-band activity (15-30 Hz) implicated in motor inhibition, under in-phase, anti-phase, or sham stimulation.

Anti-phase tACS produced a robust suppression of beta synchrony relative to both in-phase and sham stimulation, consistent with destructive interference of the endogenous rhythm. Behaviorally, anti-phase tACS reduced inhibition performance by impairing the stopping process, whereas in-phase stimulation reduced peak force rate during go trials and decreased variability of the go response, consistent with increased inhibitory tone and stabilization of ongoing motor states. 

These results provide converging behavioral and neurophysiological evidence for destructive interference of anti-phase tACS and establish a mechanistic framework for phase-specific closed-loop neuromodulation, paving the way to implement a non-invasive therapeutic strategy aimed at desynchronizing pathological beta in neurological conditions such as Parkinson's disease.       

\end{abstract}

\begin{corrauthor}
inge.leunissen\at maastrichtuniversity.nl
\end{corrauthor}

\section*{Introduction}\label{s:introduction}

    Neural oscillations provide a temporal framework for coordinating communication across distributed brain networks. Among these rhythms, beta oscillations (15-30 Hz) are especially central to sensorimotor control. In a healthy brain, beta oscillations naturally desynchronize during movement initiation and execution, and re-synchronize when a rapid halt is required \citep{Wessel2013, Schaum2021,Leunissen2022,Kilavik2013,Engel2010,Barone2021}. These state-dependent shifts enable flexible transitions between executing and withholding actions, and disruptions to this dynamic can have profound consequences for motor control.

    Parkinson's disease (PD) provides a prominent example of such a disruption. In PD, beta synchronization within and between nodes of the cortical-basal ganglia circuit becomes pathologically elevated, and this excessive coupling is strongly associated with the severity of the bradykinesia and rigidity \citep{Kuhn2008,Little2012, Chen2010, Toledo2014}. Consequently, reducing pathological beta synchrony may serve as a pivotal strategy for restoring motor function in PD. Current treatments such as levodopa therapy and deep brain stimulation (DBS) indeed reduce the degree of beta synchronization \citep{Tiwari2012, Ray2008}; however, both approaches have limitations, including side effects, invasiveness, and variable efficacy over time \citep{Brittain2014,Chen2019}. This motivates the development of non-invasive methods that can modulate pathological beta dynamics.

    Transcranial alternating current stimulation (tACS) offers a non-invasive avenue for interacting with endogenous neural rhythms and has therefore gained significant interest as a potential tool for modulating beta activity. By delivering low-intensity sinusoidal currents at targeted frequencies, tACS can induce phase alignment of neuronal spiking activity \citep{Ozen2010,Johnson2020,Krause2019} and modulate brain oscillations through a process known as entrainment \citep{Thut2011,Herrmann2016}. Enhancing beta oscillations with tACS in healthy young adults leads to reduced motor output and stronger motor inhibition \citep{Pogosyan2009,Joundi2012,Leunissen2022, Tan2022}. However, conventional open-loop tACS typically reinforces the oscillatory synchrony when applied for extended durations \citep{Karabanov2016}. This property fundamentally limits its therapeutic potential in PD, where the clinical objective is to reduce pathologically elevated beta. To overcome this constraint, we propose the application of tACS in anti-phase relative to the endogenous beta oscillations, leveraging phase cancellation as a mechanism to actively disrupt pathological beta synchrony.

    Phase cancellation, a fundamental concept in wave physics, involves destructive interference between two waves of the same frequency but opposite phases. In neuroscience, this principle has been translated into the idea that externally applied stimulation can reduce ongoing neural oscillations when delivered in the opposite phase. Preliminary research indicates that this approach holds promise for modulating pathological neural oscillations. For example, \cite{Brittain2014} demonstrated that delivering tACS in the opposing phase to the tremor rhythm can reduce Parkinsonian tremor. Similarly, computational modelling by \cite{Krause2022} suggests that when the phase offset between tACS and ongoing neural oscillations approaches 180 degrees, a transient reduction in oscillation amplitude occurs. These are among the first indications that this idea works in an experimental context at alpha frequencies \citep{Chen2023}.

    Given the potential risk of inadvertently exacerbating beta synchronization in patients with Parkinson's disease (PD) if the approach proves ineffective, we first designed a proof-of-concept study in healthy volunteers. We employed a closed-loop strategy in which beta oscillations in the pre-supplementary motor area (preSMA) were monitored in real-time using electroencephalogram (EEG), and phase-locked tACS sequences were administered in response to the ongoing beta phase. Participants performed a stop-signal task \citep{SlaterHammel1960,Coxon2006,Verbruggen2019}, which probes both movement execution and inhibition, providing a suitable paradigm to assess motor control. This setup enabled us to investigate the impact of phase mismatches between cortical stimulation and brain oscillations on (1) endogenous beta de-/synchronization, and (2) motor performance. 

    As a first step, we evaluated the accuracy and precision of the closed-loop tACS system, which integrates a phase prediction method to phase-lock tACS to brain oscillations while compensating for system delay \citep{Onojima2021,Shirinpour2020}. Our results revealed that the accuracy and precision of the phase locking are sufficiently high to deliver in-/anti-phase stimulation effectively. We then evaluated beta de-/synchronization and motor performance across three stimulation conditions: in-phase, anti-phase and sham. Our findings indicate that compared to the sham condition, in-phase tACS increases beta synchronization and improves motor inhibition, whereas anti-phase tACS reduces beta synchronization and disrupts inhibition. Finally, we observed a positive correlation between the degree of beta de-/entrainment induced by tACS and corresponding behavioral changes. 

    These findings suggest that phase-specific tACS can bidirectionally modulate beta oscillations and motor control, offering a potential non-invasive approach for disrupting pathological beta synchronization in PD. Future work should assess whether anti-phase stimulation can translate into therapeutic benefit in PD.

\section*{Methods}\label{s:methods}

\subsection*{Participants}

38 (16 males) healthy, right-handed (laterality range 20-100, mean 94.61) 18-35-year-old (mean age: 20.47$\pm$2.54) participants were enrolled in this study. Standard screening verified that there were no contraindications for non-invasive brain stimulation \citep{Bikson2009,Woods2016}. The study protocol was approved by the local ethical committee and all participants gave written informed consent prior to participation.

\subsection*{Experimental procedures}

Figure \ref{fig:fig1} illustrates the general experimental procedure. During the experiment, participants underwent a single session with three different stimulation conditions. The session began with a practice block of the stop-signal task \citep{SlaterHammel1960,Coxon2006} including 20 go-only trials and 20 go/stop-mixed trials (detailed in the following section).  Subsequently, we recorded one minute resting EEG while participants kept their eyes open and fixed on a black screen, followed by one minute of task-EEG during performance of the stop signal task. The individual beta peak frequency (IBF) was calculated based on the task-EEG and used as a stimulation frequency for closed-loop tACS (see IBF determination section for more detail). Next, the pre-supplementary motor area (preSMA) was stimulated using tACS either in-phase or anti-phase relative to the ongoing endogenous beta oscillations, based on real-time EEG monitoring. Additionally, a sham condition was included, in which only a brief, one-cycle 0.5 Hz current was applied to mimic the sensation of stimulation without effectively modulating brain activity.

\begin{figure}
    \centering
    \includegraphics[width=0.9\textwidth]{Figures/Fig1_experiment.pdf}
    \caption{\textbf{Experimental Procedure and Stop-Signal Task Design.} Top Panel: The experimental timeline consists of three conditions (in-phase, anti-phase, and sham). Each condition includes three alternating blocks of resting EEG (1 minute) and task EEG (6 minutes) with concurrent tACS stimulation. A 10-minute break separates the conditions.\\
    Middle Panel: Timing of tACS stimulation and stop-signal task within a trial. Stimulation lasts for 2 seconds, with the stop-signal task occurring during the middle 1 second.\\
    Bottom Panel: Stop-signal task. In the majority of trials, participants pinch a force transducer to stop a moving visual indicator as close as possible to the target line. In stop-signal trials, the blue indicator stopped automatically before reaching the target line, serving as a cue for participants to attempt to withhold their response.
    }
    \label{fig:fig1}
\end{figure}

Each stimulation condition commenced with a one-minute resting-state EEG while participants kept their eyes open and fixed on a black screen, followed by a block of the stop-signal task containing 50 go/stop trials with concurrent tACS stimulation applied. Within each trial, the tACS train lasted 2 seconds, with the stop-signal task trial occurring for 1 second in the middle of the stimulation period. The inter-trial interval was set to 5 seconds, during which a white fixation cross was presented on the black screen, and the ongoing EEG signal was used to calculate and predict the optimal phase for timing the onset of the next tACS sequence. This “rest and task” combination was repeated three times within the same condition, and a final one-minute of resting EEG segment was collected before concluding the stimulation block, resulting in 150 trials (of which 50 stop trials) and about 21 minutes in each condition. The order of the go/stop trials within each block and the sequence of stimulation conditions were pseudo-randomized and counterbalanced across participants to control for order effects.

After completion of each stimulation condition, participants were given a 10-minute break and were asked to fill out a Visual Analogue Scale (VAS) questionnaire to assess stimulation-induced discomfort. The VAS consisted of a 10 cm horizontal line with two opposing limits displayed at either end of the line (“no feeling” to “strong pain/discomfort”). The total duration of an experimental session was ~ 83 minutes.

\subsection*{Stop-signal task}

Participants were asked to comfortably sit 1 meter away from the screen (refresh rate 240 Hz) and perform an anticipated version of the stop-signal task \citep{SlaterHammel1960,Coxon2006,Leunissen2017} (Figure \ref{fig:fig1} bottom panel). The task was adapted from the Open-Source Anticipated Response Inhibition Task (OSARI; \citep{He2021}) and implemented in PsychoPy (version 2020.2.10; \citep{Peirce2019}). Modifications were made to the OSARI script to enable precise trial synchronization with the tACS stimulation, allowing for real-time alignment of task events with the stimulation protocol. Each trial began with the display of an empty bar at the center of the screen, with an indicator that progressively filled it from bottom to the top in 1000 ms. Two arrows located adjacent to the bar at 800 ms marked the target line, the color of which changed every time at the end of the trial as feedback on their performance (green, yellow, orange, and red, indicating the distance between the indicator and the target line within 20, 40, 60, and $>$60 ms). Participants were instructed to pinch a force transducer (OMEGA Engineering, Norwalk, CT, USA) with their right index finger and thumb to stop the indicator as close to the target line as possible. The force response threshold was initially set at 25\% of their maximum voluntary force (MVF), but could be reduced in 5\% increments if needed, allowing participants to comfortably reach the threshold throughout the session (mean: 24.1\%; range: 15\%-25\%). MVF was determined as the highest force recorded during three 5-second maximal strength pinches. 

In 33\% of the trials, known as stop trials, the bar would unexpectedly stop filling before reaching the target line. In these cases, participants were instructed to withhold pressing the sensor. A staircasing algorithm was used to adjust the stop signal delay (SSD), enforcing a 50\% success rate in inhibiting responses within each condition (initial SSD 600 ms; step size 25 ms). Stop trials were considered as failed stops (SF) if the participant pressed the sensor hard enough to cross the 25\% MVF threshold or successful stops (SS) if their response remained below this threshold. Feedback on stop trials was provided through color changes in the arrows: green for SS trials and red for SF trials.  

The force signal was acquired with an A/D Instruments Powerlab 26 Series (Dunedin, New Zealand) and stored on a trial-by-trial basis—from the moment the indicator began to fill until 1000 ms—using LabChart, with a sampling rate of 1000 Hz.

\subsection*{tACS}

tACS was applied using a 4 $\times$ 1 HD-tACS (DC STIMULATOR PLUS, NeuroConn, GmbH, Ilmenau, Germany) setup. The electrodes were embedded in a gel-filled cup that was made out of plastic (\diameter 2cm at top, 2.5 cm at bottom, height: 1.3 cm) and mounted in an EEG cap (EASYCAP GmbH, Germany). Electrodes were placed over the preSMA, with the central electrode located at FCz and four others surrounding it at F1, F2, C1, and C2 based on the 10-20 system (Figure \ref{fig:fig2} A). Electrode gel (OneStep Cleargel) was filled in the cup to achieve an impedance level below 10 k$\Omega$. Across all participants, the mean impedance averaged over channels was 7.46 $\pm$ 3.10 k$\Omega$.

The default peak-to-peak stimulation amplitude was set at 2 mA, but it was reduced if the participant experienced discomfort. For one participant, the amplitude was set to 1.6 mA, resulting in an average intensity of 1.99$\pm$0.08 mA.

In in-phase and anti-phase trials, stimulation was delivered at the individual beta frequency (IBF) measured during the stop-signal task. Each train consisted of a 0.5-second ramp-up, one second of continuous stimulation time-locked to the stop signal task trial, and a 0.5-second ramp-down. In the sham condition, a 0.5 Hz one-cycle current starting at the rising phase (0$^\circ$) was employed for two seconds without ramp-up/down.

\begin{figure}
    \centering
    \includegraphics[width=0.9\textwidth]{Figures/Fig2_system.pdf}
    \caption{\textbf{Closed-Loop tACS-EEG System Setup.} (\textbf{A}) 4$\times$1 tACS setup with electrodes placed over the preSMA (centered at FCz) and surrounding regions (F1, F2, C1, C2). (\textbf{B}) Real-time closed-loop tACS-EEG system workflow. EEG data from preSMA is processed to predict the phase of beta oscillations, enabling phase-locked tACS delivery synchronized with the stop-signal task. (\textbf{C}) Simulated electric field distribution. (\textbf{D}) Phase locked tACS for in-phase, anti-phase, and sham conditions.}
    \label{fig:fig2}
\end{figure}

\subsection*{EEG}

EEG was recorded using the actiChamp Plus amplifier (Brain Products GmbH, Gilching, Germany) with passive Ag/AgCl electrodes secured in plastic EEG caps (EASYCAP GmbH, Germany). The EEG signal was streamed in real time via Lab Streaming Layer (LSL) using the ActiChamp LSL Connector (sampling rate: 10,000 Hz)  \citep{Kothe2025}. The electrode placement followed the 10-20 system for localization, with 6 electrodes positioned at Fz, FC1, FC2, Cz, A2 (reference), and AFz (ground). By applying the Onestep Cleargel from Medcat (Klazienaveen, the Netherlands), the impedance was kept below 10 k$\Omega$. Event markers were recorded for the beginning of the trial, the stop signals, and responses.

\subsection*{Online phase-locked closed-loop tACS-EEG}

\subsubsection*{IBF determination}

The individual beta frequency (IBF) was determined as follows. A Laplacian montage was first applied to channel Fz by subtracting the average of surrounding channels (Cz, FC1, and FC2). The spatially filtered signal was then zero-phase FIR filtered (order 128, two-pass) in the beta frequency band (18-22 Hz) using the FieldTrip toolbox \citep{Oostenveld2011}. The FFT of the filtered signal was calculated with zero-padding to 4096 points, and the IBF was detected as the frequency with maximum magnitude in the spectrum \citep{Mansouri2017,Shakeel2020}. Across participants, the IBF was 20.07 $\pm$ 1.28 Hz (mean $\pm$ SD; n = 29).

\subsubsection*{Online EEG analysis and phase estimation}

To deliver the tACS sequence with a predetermined phase with respect to endogenous beta oscillations, an online phase-locked closed-loop tACS-EEG system was established, as illustrated in Figure \ref{fig:fig2} B. End-to-end system delay was empirically measured and compensated in the code (Supplementary 1). The script that was used to process the EEG data and send triggers to the tACS remote was customized in MATLAB R2022a based on the script from Shirinpour et al. \cite{Shirinpour2020}. A 500-ms time window was applied to the streamed data, updating with each new sample chunk (8\textasciitilde12 samples every 0.01 s). The EEG data within this time window underwent the following steps: (1) re-referencing using a surface Laplacian approach, computed by subtracting the average of the surrounding electrodes (Cz, FC1 and FC2) from the central electrode (Fz); (2) filtered by a zero-phase band-pass FIR filter in the range of IBF $\pm$ 2 Hz; (3) the signal edges were trimmed to remove filtering artifacts; and (4) the data was fed into an Autoregressive algorithm (https://github.com/OpitzLab/CL-phase/tree/master/AR), where AR coefficients were generated using the Yule-Walker method to forward-predict the beta wave (as detailed in Zrenner et al. \cite{Zrenner2018}); (5) the instantaneous phase of the predicted wave was determined via the Hilbert transform; (6) once the predicted phase reached \( 90^\circ \pm 5.73^\circ \) (peak), a digital trigger was sent simultaneously to initiate the tACS sequence and Stop-signal trials via an LPT port. With a system delay of 16 ms, the phase 16 ms ahead was predicted, ensuring perfect synchronization of the task onset and phase-locked tACS sequence. To accommodate the 0.5-second ramp-up period for tACS, a corresponding 0.5-second delay was implemented in Psychopy for trial initiation, ensuring that the task period coincides with the 2-mA stabilized tACS current. The tACS onset phase for each trial was set to \( 90^\circ \) for in-phase, \( 0^\circ \) for sham, and \( 270^\circ \) for anti-phase conditions. The performance of the closed-loop system was evaluated using measures of phase prediction accuracy, bias, and variance, as detailed in Supplementary 1.

\subsection*{EEG analysis}

The EEG data were preprocessed using the FieldTrip toolbox \citep{Oostenveld2011} in MATLAB 2022a. Offline analyses focused on channels FC1 and FC2 based on their proximity to fronto-central regions likely associated with pre-supplementary motor area (preSMA) activity during motor inhibition \citep{Schevernels2015,Liebrand2017,Wessel2019}, and adjacent to contralateral sensorimotor cortex that are known generators of post-movement beta rebound (PMBR) \citep{Cardellicchio2020}.

\subsubsection*{Inter-stimulation rest EEG}

EEG data were first segmented into epochs ranging from 0.1 seconds to 3 seconds after the stimulation. Preprocessing involved demeaning, detrending, and filtering. A bandpass filter (1-100 Hz) removed low-frequency drifts and high-frequency noise, while a notch filter (49-51 Hz) eliminated power line interference. Next, EEG data were re-referenced to the common average of all channels. Trials containing artifacts were detected via visual inspection and removed to preserve data integrity. The same preprocessing pipeline was applied to all conditions. EEG data of two participants were removed due to excessive noise at FC1.

We employed a wavelet-based time-frequency decomposition to analyze the spectral content of EEG signals over time. The continuous wavelet transform was applied using a frequency range of 1-45 Hz (0.5 Hz steps) and a time range of the whole epoch (0.02-second steps). The wavelet width increased linearly from 1 to 13 cycles across frequencies to balance temporal and spectral resolution. The analysis generates power spectra averaged across trials within each condition. Time-frequency representations were saved for each participant, condition (in-, anti-phase, and sham), and trial type (go (GO), stop-signal (SS), and stop-failure (SF)), for statistical analyses. 

\subsubsection*{Pre/post rest EEG}

The one-minute pre- and post-EEG for each condition was extracted, epoched into 1-second segments and preprocessed with the same steps as the inter-stimulation rest EEG. A fast Fourier transform (FFT) was performed using the multi-taper method with a Hanning taper. Power spectra were computed across a 1-45 Hz range with a 1 Hz frequency resolution, and data were padded in a 10-s window to enhance spectral resolution. The resulting spectra were then averaged over epochs. The 1/f component was removed by using a log-linear fit to the power spectrum \citep{Haegens2014,Nikulin2006}. Individual spectra were averaged, with the standard deviation represented illustrating variability across participants. 

\subsubsection*{Task EEG}

We previously demonstrated that inhibition success depends on the phase of the beta cycle at which the stop-signal is presented \citep{Fang2024}. To investigate how tACS in- or anti-phase with respect to the endogenous beta rhythm would affect this relationship, we determined the stimulation phase at which stop signals were presented. For each trial, we computed the instantaneous phase of the in-/anti-phase tACS waveform at the moment of stop-signal onset. The analytic phase was then estimated using a Hilbert transform. Then, we categorized the stop trials (SS and SF) into 'up' or 'down' trials based on the stop signals at the 'up phase' (0$^\circ$-180$^\circ$) or 'down phase' (180$^\circ$-360$^\circ$), respectively.

To establish a reference for how beta activity normally evolves during the task in the absence of stimulation, we also analyzed task EEG data from the sham condition. The data were first segmented into epochs from 0.3 to 1.7 seconds relative to sham stimulation onset (the stop signal task duration is from 0.5 to 1.5 seconds to the stimulation). A bandpass filter (5-300 Hz) was applied to remove low-frequency drifts including the sham tACS noise and high-frequency artifacts. The following steps are similar to the aftereffects EEG data: a notch filter (49-51 Hz) was applied; bad epochs were excluded by visual detection; re-referencing based on the average of all channels; wavelet-based time-frequency decomposition was performed to generate power spectra. Time-frequency representations were categorized into: GO, SS and SF for further comparison. 

\subsection*{Behavioral analysis}

\subsubsection*{Force}

Force data, originally recorded in volts, was converted to Newtons. A fifth-order 20 Hz low-pass Butterworth filter was applied to remove high-frequency noise, and baseline correction was performed by subtracting the average force from 400 to 150 ms before the target line. Force measures - peak force (maximum force), peak force rate (force increase rate), and onset of the force - were computed for each trial. Outliers of the force measures were defined as values exceeding 2.5 standard deviations above or below the median and were excluded. We categorized the force metrics into GO, SS, and SF trials across different stimulation conditions.

\subsubsection*{Response times}

Average response times (RT) in GO and SF trials were calculated. Trials with early responses ($>$400 ms before the target) or omissions ($>$1000 ms) were considered errors and removed from both the force and RT data. 

\subsubsection*{Non-parametric analysis}

Stop-signal reaction time (SSRT) estimation depends on the independent race model, which assumes context independence between go and stop processes, we verified this assumption. As expected, response time on go trials (RTGOs) were longer than response times on stop-failed trials (RTSFs, Figure \ref{fig:supp2}, \citep{Verbruggen2019,Bissett2021}). SSRT was estimated non-parametrically using the integration method with replacement of go omissions \citep{Verbruggen2019}. For each condition, we also calculated the probability of early respond go trials (P(Go errors)), go omission (P(Go omission)), successful inhibition (P(Inhibition)) and mean stop signal delay (SSD). 

\subsubsection*{Parametric analysis: Bayesian modelling}

To analyze the stop-signal performance data parametrically under three conditions (in-phase, anti-phase, and sham tACS), we employed the hierarchical BEESTS-CV modeling (Bayesian Ex-Gaussian Estimation of Stop-Signal RT-Context-Independence Violation) \citep{Matzke2021}. This approach is based on the horse-race model assuming that response inhibition depends on the relative finishing times of go and stop processes \citep{Matzke2013,Matzke2016}. On a stop signal trial, SS occurs if the go process takes longer to complete than the SSD and the time required for the stop process to finish. While if the go process completes faster than this combined time (SSD + stop process finishing time), SF occurs.

The go and stop runners' finishing time follows an ex-Gaussian distribution with parameters \( \mu \), \( \sigma \), and \( \tau \). For each condition (in-phase, anti-phase, and sham), we estimated nine parameters per participant: \( \mu_{\text{go}} \), \( \sigma_{\text{go}} \), and \( \tau_{\text{go}} \) for the go process, \( \mu_{\text{stop}} \), \( \sigma_{\text{stop}} \), and \( \tau_{\text{stop}} \) for the stop process, PTF for the probability of trigger failures, PGF for the probability of go failures, \( M \) and \( S \) indicating the finishing time distribution (location and scale) of the go runners that were shaped by the filled-interval illusion on stop-signal trials \citep{Ornstein1969}. PTF and PGF parameters were transformed into a probability scale. Considering the truncated distribution, we transformed the location and scale of the parameters into means and standard deviations. The mean go reaction time (GORT) and Stop signal reaction time (SSRT) on the population level was estimated by computing \( \mu_{\text{go}} + \tau_{\text{go}} \), and \( \mu_{\text{stop}} + \tau_{\text{stop}} \), from the posterior distributions.

The analysis was conducted using the EXG-SS model type of the Dynamic Models of Choice (DMC, \citep{Heathcote2018}) package in the R programming environment (R Core Team, 2015). Non-hierarchical Bayesian estimation was first performed to fit individual-level data. The initial priors of the non-hierarchical fitting with lower and upper bounds were reported in the Supplementary. Hierarchical fitting was then performed using the estimated parameters from the non-hierarchical fitting as the initial values. Differential Evolution Markov Chain Monte Carlo (DE-MCMC, \citep{Braak2006}) sampling with 100 iterations per chain was used to generate posterior distributions for each parameter. We modeled the data for each condition separately. 

To evaluate the goodness of fit of the BEESTS-CV model, we employed two complementary approaches. First, we assessed the descriptive accuracy of the censored BEESTS model by simulating 1000 new data sets from the posterior distribution of participant-level parameters and comparing the predicted data against the observed data with respect to: the average cumulative distribution functions of GORTs and signal-respond RTs (SSRTs), inhibition functions, and median SSRTs across SSDs. Across all measures, the predicted data closely resembled the observed distributions (Figure \ref{fig:supp4}, \ref{fig:supp5}), demonstrating that the model provided a good account of the behavioral performance. Second, we examined whether the context independence assumption of the race model was satisfied by computing a delta function from the difference between the cumulative distribution functions (CDFs) of GORTs and SSRTs across different probabilities \citep{Matzke2021}. A positive and increasing delta function suggests that context independence holds. On the other hand, if the delta function is negative or decreasing, it indicates a violation of context independence. Goodness of fit is assessed visually based on the discrepancies between the observed delta function and delta functions generated by the 1000 posterior predictive simulations. As shown in Figure \ref{fig:supp6}, the observed delta function is positive and has a steeply increasing pattern in all conditions, meaning that the model with context independence describes the data very well.

\subsection*{Statistical analysis}

All statistical analyses were carried out using R 4.5.0 (R Core Team, 2021) and Matlab R2022a (Mathworks, Natick, MA, USA). 

\subsubsection*{VAS}

The VAS scores of discomfort for each stimulation condition were analyzed using repeated measures ANOVA with stimulation condition as the within-subject factor (Table \ref{tab:supp1}, \ref{tab:supp2}). 

\subsubsection*{Force outcomes}

Linear mixed-effects (LME) models were used to analyze force outcomes (peak force, peak force rate, total force, onset time, and time to peak) with stimulation conditions (in-phase, anti-phase, sham) and trial type (GO, SS, and SF) as fixed factors. Random intercepts were modeled at the subject level to account for individual variability in force output, and models were fitted using restricted maximum likelihood (REML). F-statistics, degrees of freedom, and $p$-values of the LME result are reported. Tukey post-hoc contrasts were further examined, with p-values and estimates (mean difference $\pm$ standard error) reported. The analysis was performed in R, utilizing packages nlme 3.1-147 \citep{Pinheiro1999} and multcomp 1.4-13 \citep{Hothorn2008}, for lme model fitting, post-hoc testing, and result extraction.

To test whether stopping performance depended on the relative phase of the tACS waveform, we compared force measures between stop signals presented at the up phase (0$^\circ$-180$^\circ$) and down phase (180$^\circ$-360$^\circ$) of the stimulation cycle. Paired-samples t-tests were conducted within each stimulation condition (in-phase, anti-phase) to assess differences between up- and down-phase for both failed (SF) and successful stop (SS) trials.

\subsubsection*{Non-parametric estimates}

To compare non-parametric estimates of stop-signal performance across in-, anti-phase and sham groups, we applied a Friedman test followed by Wilcoxon signed-rank tests for P(Inhibition), as the data violated the assumption of normality (Shapiro-Wilk test $p < 0.05$). We conducted repeated measures ANOVA with post-hoc tests for the remaining dependent variables (GORT, SSRT, SFRT, SSD, P(Go errors), and P(Go omissions)).

\subsubsection*{BEESTS-CV parameters}

The mean of the posterior distributions of the BEESTS model parameters in three stimulation conditions were reported with a 95\% credible interval (CI; $2.5^{th}-97.5^{th}$ percentiles). To evaluate group differences, Bayesian p-values were indicated as the proportion of samples in one group's distribution exceeding those in another, with values near 0 or 1 indicating significant shifts toward lower or higher parameter estimates, respectively. See \cite{Matzke2019} and \cite{Choo2022} for more detail on this approach.

\subsubsection*{EEG measures}

Statistical comparisons of time-frequency representations were conducted using non-parametric cluster-based permutation tests applied to: (1) inter-stimulation post-task EEG (time range: 0.25-1 s after stimulation), comparing conditions (in-phase, anti-phase, sham) both across all trial types and within each trial type (GO, SS, SF); (2) post-task EEG in the sham condition, comparing trial types (GO, SS, SF); and (3) on-task EEG in the sham condition, comparing trial types (GO, SS, SF). A Monte Carlo approach with 1000 randomizations was applied, using a significance threshold of $p < 0.05$. The design matrix accounted for within-subject dependencies, and significant clusters were outlined in the visualized results.

Paired t-tests were used to identify significant differences in IBF-centered power spectra between pre- and post-rest-EEG for each condition (in-phase, sham, and anti-phase). Significant differences ($p < 0.05$) were highlighted in the plots with shaded regions. 

\subsubsection*{EEG-behavior correlations}

To investigate the relationship between post-stimulation beta power and online inhibitory performance, a plausible correlation analysis was applied between posterior SSRT estimated from the BEESTS model posterior and post-stimulation EEG beta power density. Beta power was extracted within the 0.25-1 s window following stimulation and within the individualized beta frequency band (IBF $\pm$ 2 Hz), and averaged across all trials per condition. A Bayesian hierarchical approach was then used to obtain the population-level distribution of correlation coefficients. Significance was indicated by the probability that the true correlation exceeds zero ($p(r>0)$) is larger than 0.95 or smaller than 0.05.

\section*{Results}\label{s:results}

\subsection*{Inter-stimulation EEG}

All inter-stimulation EEG analyses focused on 0.25-1 s after tACS offset (2.25-3.0 s relative to tACS onset), centered on the individual beta frequency (IBF) at FC1/FC2. Statistical significance was determined using cluster-based permutation testing, with a threshold of $p < 0.05$.

\subsubsection*{Induced beta dynamics in the inter-stimulation interval}

Figure \ref{fig:fig3} illustrates post-stimulation beta activity averaged across all trials for each condition. Figure \ref{fig:fig3} A shows the EEG power spectrum baseline-corrected to the end of the stimulation-free segment (4.5-4.7 s, z-score), and Figure \ref{fig:fig3} B shows paired t-statistic contrasts. These maps indicate that anti-phase stimulation yielded a significant reduction in beta power relative to both sham and in-phase, with clusters centered on IBF and extending up to \( \sim 0.95 \) s after stimulation offset. No significant clusters were observed for the in-phase vs. sham comparison. 

\begin{figure}[h]
    \centering
    \includegraphics[width=0.9\textwidth]{Figures/Fig3_TFR_all.pdf}
    \caption{Time-frequency representations (x: time [s], y: frequency [Hz]) of tACS aftereffects at FC1/FC2 during the inter-stimulation interval (0.25-1 s post stimulation), aligned to each participant’s individual beta frequency (IBF). (\textbf{A}) Baseline-corrected power (z-score; baseline 4.5-4.7 s) shown separately for in-phase, anti-phase, and sham. (\textbf{B}) Cluster-based paired t-contrasts ($p < 0.05$, TFCE corrected) between conditions: in-phase > anti-phase, in-phase > sham, and anti-phase > sham. Significant clusters are outlined in black.}
    \label{fig:fig3}
\end{figure}

\subsubsection*{Phase-specific modulation of trial-type dynamics}

To establish a functional baseline for trial-type comparisons, we first examined beta activity under sham stimulation (Figure \ref{fig:fig4}). Time-frequency maps revealed significantly stronger beta activity following GO and SF trials compared to SS trials, consistent with the established association between PMBR and motor execution \citep{Pfurtscheller1996,Cassim2001,Tan2016} (Figure \ref{fig:fig4} AB). In contrast, SS trials did not involve movement execution and therefore did not evoke a PMBR, but rather showed beta synchronization directly after the stop signal during stimulation (Figure \ref{fig:fig4} CD). This baseline analysis confirmed that movement and inhibition engage different beta dynamics, providing a reference for assessing stimulation effects.

\begin{figure}[h]
    \centering
    \includegraphics[width=0.9\textwidth]{Figures/Fig4_TFR_sham.pdf}
    \caption{Time-frequency representations (x: time [s], y: frequency [Hz]) at FC1/FC2, aligned to each participant’s individual beta frequency (IBF). (\textbf{A}) Post-task baseline-corrected power (z-score; baseline 4.5-4.7 s) shown separately for go, stop-successful (SS) and stop-failed (SF) trials during the inter-stimulation interval (2.1-3.0 s). (\textbf{B}) Cluster-based paired t-contrasts ($p < 0.05$, cluster corrected) of post-task EEG power between trial types (GO $>$ SS, GO $>$ SF, SF $>$ SS). Significant clusters are outlined in black. (\textbf{C}) On-task baseline-corrected power (z-score; baseline 0.7-1.1 s, average stop signal presentation time = 1.1) shown separately for go, stop-successful (SS) and stop-failed (SF) trials. (\textbf{D}) Cluster-based paired t-contrasts ($p < 0.05$, cluster corrected) of on-task EEG power between trial types (GO $>$ SS, GO $>$ SF, SF $>$ SS). Significant clusters are outlined in black.}
    \label{fig:fig4}
\end{figure}

We then examined how stimulation modulated these trial-type specific dynamics (Figure \ref{fig:fig5}). The most robust effect was observed for anti-phase stimulation, which significantly disrupted the beta rebound relative to sham. Following GO trials (Figure \ref{fig:fig5} A), anti-phase stimulation led to a marked suppression of beta activity around the IBF, lasting up to \( \sim 0.95 \) s post-stimulation (Figure \ref{fig:fig5} A). For SF trials, anti-phase again produced significant suppression of beta power, elicited at 0.3-0.5 s around IBF (Figure \ref{fig:fig5} B). For SS trials (Figure \ref{fig:fig5} C), anti-phase reduced stop-signal-related beta synchronization relative to sham and in-phase, with a transient effect at 0.5-0.6 s. Together, these results show that anti-phase stimulation consistently attenuates both motor- and inhibition-related beta dynamics across trial types.

\begin{figure}[h]
    \centering
    \includegraphics[width=1.1\textwidth]{Figures/Fig5_TFR_phase.pdf}
    \caption{Time-frequency representations (x: time [s], y: frequency [Hz]) for cluster-based paired t-statistic ($p < 0.05$, TFCE corrected) between conditions within each trial type. Spectra are at FC1/FC2 during the inter-stimulation interval (2.25-3.0 s), aligned to each participant's individual beta frequency (IBF). Panels depict (\textbf{A}) GO, (\textbf{B}) SF, and (\textbf{C}) SS trials. Rows show condition contrasts: in-phase vs anti-phase (top), in-phase vs sham (middle), and anti-phase vs sham (bottom). Significant clusters are outlined in black ($p < 0.05$).}
    \label{fig:fig5}    
\end{figure}

\subsection*{Pre-post rest EEG results}

Paired t-test between 5-second pre- and post-rest EEG power spectra (FC1/FC2) revealed a significant decrease within the frequency band of IBF$\pm$1 Hz after stimulation (Figure \ref{fig:fig6}). This reduction was not present in the in-phase and sham conditions. Taken together, anti-phase tACS induces frequency-specific suppression of post-stimulation beta oscillations. 

\begin{figure}[h]
    \centering
    \includegraphics[width=0.5\textwidth]{Figures/Fig6_ER.pdf}
    \caption{Pre- and post-stim EEG power spectra (IBF\(\pm\)5 Hz) across three conditions. Pre-spectra are in blue, and post-spectra are in red. Shaded regions represent variability (\(\pm\)SEM), and gray areas indicate frequency ranges with significant pre-post differences from the paired t-test. Anti-phase stimulation resulted in a significant decrease in beta power around IBF\(\pm\)1 Hz.}
    \label{fig:fig6}    
\end{figure}

\subsection*{BEESTS-CV model results}

The inhibitory performance in each stimulation condition was estimated based on the finishing time distributions of the ‘go’ and ‘stop’ runners, as captured by the parameter estimates from the BEESTS-CV-DMC model.
Table \ref{tab:tb1} summarizes the posterior means and 95\% credible intervals (CIs) for each model parameter across conditions, along with the difference between posterior distributions. Statistical significance is represented by Bayesian p-values, defined as the proportion of the posterior distribution of the latter condition that exceeds \( (p > 0.95) \) or falls below \( (p < 0.05) \) the former condition. Figure \ref{fig:fig7} presents the posterior distributions for each population-level parameter, aligning with the statistics in Table \ref{tab:tb1}.

For the ‘stop’ process, we observed an increase in \( \mu_{\text{stop}} \) in the anti-phase (196 ms) compared to the in-phase (191 ms) condition, suggesting that anti-phase stimulation might slow the stop process. Moreover, \( \tau_{\text{stop}} \) was higher in the in-phase (21 ms) condition relative to the anti-phase (12 ms). The overall SSRT remained comparable across conditions, likely due to a trade-off between \( \mu_{\text{stop}} \) and \( \tau_{\text{stop}} \) components.
For the ‘go’ process, the posterior distribution of \( \sigma_{\text{go}} \) is significantly lower in the in-phase condition (33 ms) compared to sham (35 ms). \( \tau_{\text{go}} \) was increased in both the in-phase (18 ms) and anti-phase (18 ms) conditions relative to sham (9 ms), while no significant difference was found in \( \mu_{\text{go}} \), which was translated into a significantly slower mean GORT \( (\mu_{\text{go}} + \tau_{\text{go}}) \) in the anti-phase conditions (823 ms) than in sham (815 ms). This pattern suggests that anti-phase tACS did not change the central tendency of the go process, but selectively increased the tail of the finishing time distribution, reflecting a higher frequency of slow responses. In contrast, in-phase tACS appeared to reduce variability of the go finishing time, yet at the same time induced more frequent slow responses.

Finally, we observed significant conditional impacts on the filled-interval illusion, as reflected in the parameters \( M \) (shift) and \( S \) (spread) of the go finishing distribution on stop-signal trials influenced by the illusion. The \( M \) parameter was lower in the in-phase condition (3.6) relative to sham (5.7), suggesting a reduced elongation of the perceived interval following stop signals. In addition, \( S \) was significantly lower in the anti-phase condition (1.07) relative to sham (1.14), reflecting reduced variability in interval estimation under anti-phase stimulation.

We did not observe significant differences in PTF and PGF across stimulation conditions. This stable likelihood for both go and stop trigger failures indicates that the observed changes in stop and go distributions are unlikely to stem from attentional fluctuations, but rather reflect changes in inhibitory efficiency caused by the stimulation \citep{Matzke2016}.


\begin{landscape}
\begin{table}[]
    \small                     % or \small if you prefer
    \setlength{\tabcolsep}{2.5pt}     % default is ~6pt
    \renewcommand{\arraystretch}{0.9} % default is 1.0
    \caption{Posterior means and 95\% credible intervals (CIs) of the population-level mean parameters across stimulation conditions.}
\begin{tabular}{lllllllllll}
\hline
          & \multicolumn{3}{l}{Mean   (95\% CI)}                                                    & \multicolumn{2}{l}{In.-   Anti.} & \multicolumn{3}{l}{In.   - Sham}                             & \multicolumn{2}{l}{Anti.   - Sham} \\ \hline
Parameter & In-phase                    & Anti-phase                  & Sham                        & Main   diff. (95\% CI)   & p     & \multicolumn{2}{l}{Main   diff.}              & p                 & Main   diff.              & p      \\ \hline
$P_{TF}$      & 0.009 (-0.014-0.085)        & 0.019 (-0.020-0.114)        & 0.019 (-0.019-0.118)        & 0.011 (-0.078-0.106)     & .587  & \multicolumn{2}{l}{0.012 (-0.079-0.111)}      & .591              & 0.001 (-0.100-0.106)      & .505   \\
$P_{GF}$      & 0.028 (0.004-0.069)         & 0.026 (0.008-0.054)         & 0.025 (0.006-0.057)         & -0.003 (-0.049-0.035)    & .462  & \multicolumn{2}{l}{-0.003 (-0.049-0.037)}     & .449              & -0.000 (-0.035-0.036)     & .492   \\
$\mu_{stop}$  & 191.265 (187.335-195.320)   & 196.216 (192.792-199.551)   & 192.475 (188.603-196.375)   & 4.951 (-0.267-10.061)    & \textbf{.968*} & \multicolumn{2}{l}{1.210 (-4.304-6.623)}      & .670              & -3.740 (-8.936-1.489)     & .078   \\
$\sigma_{stop}$ & 10.195 (3.446-17.502)      & 8.843 (3.036-15.118)        & 7.487 (2.510-13.240)        & -1.352 (-11.800-8.669)   & .402  & \multicolumn{2}{l}{-2.708 (-12.659-6.732)}    & .293              & -1.356 (-10.390-7.455)    & .385   \\
$\tau_{stop}$ & 21.437 (13.855-26.133)      & 12.542 (4.552-19.677)       & 18.898 (7.780-25.881)       & -8.894 (-19.079-1.312)   & \textbf{.043*} & \multicolumn{2}{l}{-2.539 (-12.666-9.273)}    & .292              & 6.355 (-5.295-19.312)     & .853   \\
SSRT          & 212.701 (204.361-219.072)   & 208.500 (199.843-216.505)   & 211.349 (199.842-219.484)   & -4.202 (-15.843-7.185)   & .233  & \multicolumn{2}{l}{-1.353 (-12.938-11.521)}   & .389              & 2.849 (-9.953-16.662)     & .664   \\
$\mu_{go}$    & 801.298 (797.328-805.281)   & 804.326 (800.563-808.074)   & 805.808 (801.384-810.306)   & 3.028 (-2.516-8.465)     & .861  & \multicolumn{2}{l}{4.510 (-1.460-10.669)}     & .933              & 1.482 (-4.211-7.341)      & .696   \\
$\sigma_{go}$ & 32.929 (31.347-34.512)      & 34.470 (32.974-35.967)      & 35.397 (33.885-36.881)      & 1.541 (-0.663-3.702)     & .922  & \multicolumn{2}{l}{2.468 (0.254-4.681)}       & \textbf{.984*}    & 0.927 (-1.197-3.059)      & .812   \\
$\tau_{go}$   & 18.039 (15.846-20.121)      & 18.737 (15.675-21.460)      & 9.128 (3.424-13.828)        & 0.698 (-2.730-4.534)     & .648  & \multicolumn{2}{l}{-8.911 (-14.085--2.457)}   & \textbf{.004**}   & -9.609 (-15.406--2.900)   & \textbf{.002**} \\
GORT          & 819.337 (814.760-823.786)   & 823.064 (818.282-827.737)   & 814.809 (807.497-821.470)   & 3.727 (-2.845-10.374)    & .871  & \multicolumn{2}{l}{-4.527 (-12.603-4.325)}    & .145              & -8.254 (-16.579-0.609)    & \textbf{.034*}  \\
M             & 3.565 (1.697-5.446)         & 5.735 (3.682-7.800)         & 5.868 (3.747-7.869)         & 2.170 (-0.647-4.980)     & .936  & \multicolumn{2}{l}{2.303 (-0.429-5.144)}      & \textbf{.952*}    & 0.133 (-2.831-3.083)      & .538   \\
S             & 1.119 (1.065-1.174)         & 1.066 (1.019-1.115)         & 1.139 (1.094-1.184)         & -0.053 (-0.127-0.019)    & .076  & \multicolumn{2}{l}{0.020 (-0.052-0.089)}      & .715              & 0.073 (0.006-0.139)       & \textbf{.984*}  \\ \hline
\end{tabular}
\vspace{1.6ex}
\caption*{\footnotesize Values are reported for in-phase, anti-phase, and sham conditions. Pairwise contrasts are reported alongside the corresponding ‘Main diff’ (i.e., the value computed by subtracting the posterior samples of the latter condition from the former condition, such that positive values indicate higher estimates in the first condition) and Bayesian p-values (i.e., posterior probabilities that the parameter value in the second condition exceeds that in the first) indicated with \( p < .05 \) (*) and \( p < .01 \) (**).}
\label{tab:tb1}
\end{table}
\end{landscape}

\begin{figure}[h]
    \centering
    \includegraphics[width=1.1\textwidth]{Figures/Fig7_beests.pdf}
    \caption{Posterior distributions of population-level parameters estimated by the BEESTS-CV-DMC model across in-phase (red), anti-phase (blue), and sham (dot) conditions, aligning with the statistics in Table \ref{tab:tb1}. Asterisks indicate that the Bayesian \( p \)-values \( > 0.95 \) or \( < 0.05 \).}
    \label{fig:fig7}
\end{figure}

\subsection*{Force results}

A linear mixed-effects model was conducted to examine the effects of stimulation condition (in-phase, anti-phase, and sham) on force outcomes. The analysis revealed a statistically significant main effect of stimulation on peak force rate during GO trials (\( F(2, 10034)=4.68, p < 0.001 \)) and on response onset time during stop-signal (SS) trials (\( F(2, 2296)=3.18, p=0.04 \); Table \ref{tab:tb2}). Post-hoc pairwise comparisons (Tukey-HSD) revealed that in-phase elicited significantly lower peak force rate during GO trials compared with both anti-phase (\( z=-2.45, p=0.04 \)) and sham (\( z=-2.8, p=0.01 \); \( SE=2.32 \)). For SS trials, onset time was significantly longer under in-phase than sham stimulation (in-phase - sham: \( z=2.4, p=0.04 \), \( SE=2.87 \)).

% Please add the following required packages to your document preamble:
\begin{table}[h]
\centering
\caption{Effect of stimulation on force measures.}
\begin{tabular}{llllll}
\hline
\multirow{2}{*}{Trial type} & \multirow{2}{*}{Measures}                                                         & \multicolumn{3}{l}{Summary}                   & LME statistic                               \\ \cline{3-6} 
                            &                                                                                   & Inphase       & Antiphase     & Sham          & Main effect                                 \\ \hline
\multirow{3}{*}{GO}         & Peak force (N)                                                                    & 30.48$\pm$9.09    & 30.59$\pm$9.52    & 30.37$\pm$9.08    & F(2,10042)=2.5, p=.08                       \\
                            & \begin{tabular}[c]{@{}l@{}}Peak force rate (N/s)\\  \end{tabular} & 355.57$\pm$125.33 & 361.6$\pm$128.85  & 362$\pm$156.87    & \textbf{F(2,10034)=4.68, p\textless{}.001*} \\
                            & Onset time (ms)                                                                   & 703.78$\pm$51.26  & 704.72$\pm$55.06  & 703.45$\pm$53.6   & F(2,9984)=0.59, p=.56                       \\ \hline
\multirow{3}{*}{SF}         & Peak force (N)                                                                    & 25.10$\pm$7.61    & 25.15$\pm$8.01    & 25.12$\pm$7.52    & F(2,2362)=0.002, p=1                        \\
                            & Peak force rate (N/s)                                                             & 329.32$\pm$120.86 & 330.11$\pm$117.46 & 332.88$\pm$118.8  & F(2,2352)=0.89, p=.41                       \\
                            & Onset time (ms)                                                                   & 703.78$\pm$51.26  & 704.72$\pm$55.06  & 703.45$\pm$53.60  & F(2,9984)=0.59, p=.56                       \\ \hline
\multirow{3}{*}{SS}         & Peak force (N)                                                                    & 6.80$\pm$6.37     & 7.24$\pm$7.15     & 6.93$\pm$6.44     & F(2,2566)=1.27, p=.28                       \\
                            & Peak force rate (N/s)                                                             & 103.85$\pm$98.09  & 107.83$\pm$101.12 & 115.85$\pm$257.15 & F(2,2744)=1.45, p=.23                       \\
                            & Onset time (ms)                                                                   & 711.5$\pm$64.05   & 705.45$\pm$66.51  & 704.68$\pm$62.16  & \textbf{F(2,2296)=3.18, p=0.04*}            \\ \hline
\end{tabular}
\vspace{1.6ex}
\caption*{\footnotesize Linear mixed-effects (LME) model results for peak force, peak force rate, and onset time across in-phase, anti-phase and sham stimulation conditions during go (GO), stop-failed (SF), and stop-successful (SS) trials. Asterisks (*) denote statistically significant effects at $p < 0 .05$.}
\label{tab:tb2}
\end{table}

These findings suggest that in-phase tACS attenuates motor output and enhances inhibitory control, as reflected in reduced peak force rate during voluntary responses and delayed response onset during inhibition.

\subsection*{Correlations between beta power and behavioral measures}

\subsubsection*{Plausible correlation analysis (SSRT-beta power)}

To assess the relationship between post-stimulation beta power and the SSRT posterior estimates from the BEESTS model, we employed a plausible correlation analysis using Bayesian hierarchical modeling that accounts for individual-level uncertainty and provides a population-level distribution of correlation coefficients. The result revealed a reliable negative association in both sham and in-phase stimulation between SSRT and post-stimulation beta power (Figure \ref{fig:fig8}). Specifically, participants eliciting stronger post-stimulation beta power tended to exhibit shorter SSRTs (sham: \( r = -0.221 \), 95\% CI [\( -0.461, -0.040 \)], \( p = 0.048 \); In-phase: \( r = -0.265 \), 95\% CI [\( -0.505, 0.003 \)], \( p = 0.027 \)). In contrast, no significant correlations were observed in the anti-phase condition.

\begin{figure}[h]
    \centering
    \includegraphics[width=0.9\textwidth]{Figures/Fig8_SSRT.pdf}
    \caption{Posterior density estimates of plausible correlations (r) between IBF\(\pm\)2 EEG power and hierarchical SSRT. Solid lines represent the individual-level plausible correlation distributions, and dashed lines represent the population-level posterior distributions obtained using a uniform prior (\(\kappa = 1\)).}
    \label{fig:fig8}
\end{figure}

These correlations suggest that enhanced beta synchronization during the post-stimulation phase is linked to faster stopping processes. While anti-phase tACS may disrupt the natural relation between inhibitory efficiency and post-movement cortical beta dynamics.

\subsection*{Phase-dependency of inhibition performance}

To examine whether in-phase versus anti-phase tACS influences the relationship between stop-signal presentation phase and inhibitory control, we conducted paired t-tests comparing peak force on stop trials (SS and SF) between stop signals delivered at the down versus up phases of the online tACS waveform. Building on our previous finding that inhibition success depends on the phase of the endogenous beta cycle at which the stop signal is processed \citep{Fang2024}, this analysis tested whether externally aligning or misaligning tACS with that rhythm modulates the phase-specific effect.

The analysis revealed a significant phase-dependent modulation of peak force during SS trials exclusively under in-phase tACS: peak force was significantly lower when the stop signal occurred at the down phase compared to the up phase of the stimulation waveform (\( t(32) = -3.04 \); \( p = 0.005 \)) (Figure \ref{fig:fig9}). By contrast, anti-phase tACS did not produce any phase-dependent effects on peak force in SS trials (\( t(32) = -0.16 \), \( p = 0.88 \)), nor were significant differences observed in SF trials under either stimulation condition (In-phase: \( t(32) = -0.79 \), \( p = 0.43 \); Anti-phase: \( t(32) = 0.03 \), \( p = 0.97 \)). The in-phase results closely replicate our prior finding \citep{Fang2024} that inhibition is strongest when the stop signal coincides with the preferred phase of the beta oscillation, suggesting that in-phase tACS preserves this stable phase relationship. In contrast, anti-phase stimulation appears to disrupt this alignment, attenuating the phase-dependent modulation of inhibitory control.

\begin{figure}
    \centering
    \includegraphics[width=0.8\textwidth]{Figures/Fig9_phase.pdf}
    \caption{Box plots of peak force (N) at the up and down phases on SS and SF trials under in- and anti-phase tACS stimulation. Down and up phases refer to the tACS waveform at the time of the stop signal presentation, respectively. The double asterisk (**) indicates a significant difference between up and down stop signal phases ($p = 0.005$).}
    \label{fig:fig9}
\end{figure}

\section*{Discussion}\label{s:disucssion}

Pathologically elevated beta synchronization is a defining neurophysiological feature of Parkinson's disease (PD) and a key therapeutic target for neuromodulation \citep{Brown2007}. Motivated by this need to suppress aberrant oscillatory coupling, we developed a closed-loop tACS-EEG system delivering stimulation precisely aligned, or opposed in phase, to ongoing beta activity in the pre-supplementary motor area (preSMA). Using this approach, we investigated how phase-specific stimulation influences neural dynamics and motor inhibition. Our findings demonstrate that anti-phase tACS disrupts beta synchronization and modulates motor inhibition performance, mirroring the therapeutic goal in PD treatment.

\subsection*{Phase-specific effects of closed-loop tACS on beta synchrony}

tACS can entrain neural activity and modulate oscillatory power in a frequency- and phase-specific manner \citep{Ozen2010,Johnson2020,Krause2019,Thut2011,Herrmann2016,Helfrich2014,Tseng2016}. Recent work highlights that the phase relationship between externally applied currents and intrinsic rhythms determines whether stimulation amplifies or disrupts oscillatory activity \citep{Krause2022,Chen2023,Krause2023,Haslacher2024}. In vitro, in-phase fields promote synchrony, whereas out-of-phase fields reduce it \citep{Frhlich2010,Reato2010}. Clinically, phase-locked tACS applied at the tremor frequency has been shown to suppress tremor \citep{Brittain2014}.

In line with these predictions, our inter-stimulation EEG analyses revealed robust suppression of beta power following anti-phase tACS compared with both sham and in-phase stimulation, with effects centered around the individual beta frequency. This attenuation likely reflects destructive interference, whereby stimulation delivered in the opposite phase counteracts ongoing oscillatory activity. This suppression may represent the destructive counterpart of an outlasting 'echo' of tACS entrainment \citep{Geffen2025}.

By contrast, in-phase tACS did not significantly increase beta power, likely due to the short stimulation trains (2 s including 0.5 s ramp-up/down), which may have limited entrainment accumulation. Nevertheless, subtle behavioral modulations emerged under in-phase tACS, including reduced peak force and lower response variability during movement.

\subsection*{Ongoing brain state shapes the phase effects}

In line with numerous previous studies, our sham-condition EEG analyses replicated well-established task-related beta patterns: (1) a stop-signal-locked beta increase associated with successful inhibition \citep{Swann2009,Wagner2018,Leunissen2022}, and (2) a post-movement beta enhancement following GO and SF trials, reflecting a re-establishment of the motor state after voluntary movement \citep{Pfurtscheller1996,Cassim2001,SolisEscalante2012,Tan2016}. Going and stopping rely on partly distinct networks: go responses engage motor and premotor regions, whereas stopping recruits the rIFG-preSMA-STN inhibitory circuit \citep{Aron2007,Rae2015,Jahfari2011}. Failed stops instead involve enhanced rAI-preSMA coupling linked to salience monitoring \citep{Cai2014,Limongi2017}.

Across all trial types, anti-phase tACS reduced post-stimulation beta power, consistent with destructive interference. This effect was strongest in GO and SF trials, where stimulation ended during the natural beta rebound, producing a desynchronization “echo”. In contrast, in successful stops (SS), beta re-synchronizes earlier, time-locked to the stop signal occurring during stimulation, leaving fewer active dynamics to be disrupted. The modest desynchronization in SS trials may also reflect compensatory overshoot or recruitment of alternative networks to sustain inhibition \citep{Nowak2017,To2018,Pariz2023}. These observations align with prior work showing that tACS effects are state-dependent, varying with ongoing neural activity and task engagement \citep{Violante2017,Vosskuhl2018,AbellanedaPrez2020}. Anti-phase stimulation thus most effectively disrupts beta synchrony when delivered during strong endogenous coupling.

\subsection*{Behavioral consequences of phase-specific tACS}

Beyond the neural effects, stimulation also modulated behavior. Consistent with the observed beta desynchronization, anti-phase tACS impaired inhibitory control, as reflected in a higher \( \mu_{\text{stop}} \) estimated with the BEESTS model. Because \( \mu_{\text{stop}} \) represents the mean of the Gaussian component of the stop-signal reaction time (SSRT) distribution, it captures the typical latency of the stopping process. An increase in \( \mu_{\text{stop}} \) therefore indicates slower and less efficient inhibition, supporting the view that beta synchrony facilitates rapid stopping. However, neither the overall SSRT derived from the integration method nor the BEESTS-estimated mean SSRT differed significantly across conditions, implying that compensatory mechanisms (e.g., proactive slowing or adaptive plasticity in fronto-basal ganglia circuits) may have helped to maintain performance despite local beta disruption. Importantly, the correlation between post-stimulation beta power and SSRT that was evident in the sham and in-phase conditions was absent during anti-phase stimulation, suggesting that anti-phase tACS disrupted the normal coupling between cortical beta synchronization and inhibitory efficiency.

In a similar way, anti-phase stimulation also abolished the phase-dependent modulation of stopping that was observed when stimulation was aligned with ongoing beta activity. Peak force on stop trials was lower when the stop signal coincided with the down phase of the in-phase waveform, but this modulation was not present when the phase relationship was reversed. This mirrors our prior finding that inhibition efficiency depends on the phase of the stimulation waveform at which the stop signal occurs \citep{Fang2024}. Together, these observations suggest that aligning stimulation with endogenous beta preserves functional phase relationships, whereas misalignment disrupts the propagation of stop signals through the network.

In-phase tACS reduced the rate of force in go trials and showed delayed force onset in successful stops. The reduction in peak force rate resembles earlier findings that beta-frequency stimulation can act as an active “brake” on the motor system \citep{Pogosyan2009,Joundi2012,Leunissen2022}, potentially reflecting a transient increase in inhibitory tone or reduced corticospinal drive. At the same time, in-phase stimulation decreased \( \sigma_{\text{go}} \), indicating lower variability in go-process finishing times and the increased \( \tau_{\text{go}} \). This pattern could suggest a stabilization of internal response timing, i.e., less trial-to-trial variability, but occasional longer responses when flexibility was required. Such an effect would be consistent with proposals that beta activity contributes to the maintenance of ongoing sensorimotor states and temporal regularity \citep{Engel2010,Kilavik2013,Spitzer2017,Barone2021}. However, the precise functional contribution of beta remains debated, and these findings could also arise from enhanced timing precision, altered temporal weighting, or other network-level adjustments induced by stimulation. Thus, while in-phase stimulation appears to have modulated both the strength and stability of motor responses, the relative contribution of inhibitory versus stabilizing mechanisms cannot yet be fully dissociated.

\subsection*{Implications and limitations}

These findings provide causal evidence that phase alignment governs the behavioral impact of brain stimulation. By demonstrating that identical stimulation can either impair or facilitate inhibition depending on its phase relation to ongoing beta activity, the results highlight the importance of timing stimulation to the brain's intrinsic dynamics.

Several limitations should be acknowledged. First, our primary neurophysiological analyses focused on the after-effects of stimulation. While these after-effects suggest sustained modulation, our results were largely blind to the immediate, online impact of tACS on neural oscillations. Second, we employed only two phase conditions, in-phase and anti-phase, which limits our ability to precisely map the full spectrum of phase-dependent effects. Finally, the efficacy of tACS can vary significantly across individuals due to distinct coupling strengths to external stimulation. Future research should account for these individual differences, exploring optimized stimulation durations and intensities to enhance precision and efficacy. 

Despite these constraints, the results provide a mechanistic framework for state- and phase-dependent brain stimulation. In clinical contexts such as Parkinson's disease, where excessive beta synchrony impairs movement, anti-phase stimulation could be leveraged to transiently desynchronize pathological rhythms, whereas in-phase or phase-adaptive approaches may help restore deficient coupling in other contexts. Ultimately, closed-loop tACS offers a promising route toward individualized, temporally precise neuromodulation that aligns intervention with the brain's natural oscillatory architecture.

\section*{Conclusion}\label{s:conclusion}

This study shows that the behavioral impact of beta-frequency tACS depends critically on its phase alignment with ongoing neural activity. Anti-phase tACS disrupted beta synchrony and impaired inhibitory control. In contrast, phase-aligned stimulation modulated go performance through reduced force output and lower response variability, consistent with increased inhibitory tone and greater temporal stability. These findings offer mechanistic insight into how the precise timing of stimulation relative to intrinsic neural dynamics governs its functional outcome. Moreover, they establish a framework for individualized, temporally adaptive neuromodulation that could serve as a non-invasive therapeutic approach to desynchronize pathological beta activity in Parkinson's disease.

%\nocite{*}
\ifdefined\ismergeddocument
\else
\section*{References}
\bibliography{refs.bib}
\fi

\section*{Author information}
Zhou Fang: Writing - original draft, validation, resources, methodology, investigation, formal analysis, data curation, conceptualization. Alexander T. Sack: Writing - review \& editing, supervision, methodology, funding acquisition, conceptualization. Inge Leunissen: Writing - review \& editing, validation, supervision, methodology, funding acquisition, conceptualization.
