## Supplementary material for "Closed-loop phase-locked EEG-tACS enables adaptive control of cortical beta synchrony and motor control": Latex project: 02_Article_Supplementary.tex

% !TEX root = 00_Article_Merge.tex
%  CONFIGURE NEW SINGLE-PAGE FORMAT 

\onecolumn % go back to one column
\fancyhead{} % make sure we get no headers
\renewcommand{\floatpagefraction}{0.1}
\lfoot[\bSupInf]{\dAuthor}
\rfoot[\dAuthor]{\cSupInf}
\newpage
\captionsetup[table]{position=above}
\captionsetup*{format=largeformat} % make figure legend slightly larger than in the paper
\setcounter{figure}{0} % reset figure counter for Supp. Figures
\setcounter{table}{0} % reset figure counter for Supp. Tables
\setcounter{equation}{0} % reset equation counter for Supp. Equations
%\setcounter{page}{1} % reset page count
\makeatletter 
\renewcommand{\thefigure}{S\@arabic\c@figure} % make Figure legend start with Figure S
\renewcommand{\thetable}{S\@arabic\c@table} % make Table legend start with Table S
\makeatother
\def\theequation{S\arabic{equation}}

%  MAIN TEXT 

\newpage
\section*{Supplementary Material}
\phantomsection
\label{supplementary}

\subsection*{Supplementary 1. Closed-loop evaluation}\label{supp1}

To evaluate the performance of the closed-loop system, we recorded \(\sim\)10-minute tACS-free EEG data where one participant was performing the stop-signal task to simulate 10,000 trials of real-time beta peak detection (the algorithm iteratively continued until 10,000 peaks were detected). The performance of phase prediction was quantified offline based on accuracy, bias, and variance \citep{Shirinpour2020}. The accuracy was evaluated by the equation:
$$
1-\frac{1}{N\times180}\sum_{i=1}^{N}\left|\theta_i-\theta_0\right|
$$
where \( N \) is the number of trials used to estimate the phase; \( \theta_i \) is the estimated phase in the trials \( i \); \( \theta_0 \) is the target phase, 0°. An accuracy of 1 means that the phase has been estimated precisely every time, whereas an accuracy of 0 means that the phase has been detected opposite to the target phase, e.g., peaks instead of troughs. With an accuracy of 0.5, phase estimation is uniformly randomly distributed. Bias was determined by computing the average difference between the estimated and target phases, revealing any systematic deviations from the intended outcome. Variance, quantified using standard deviation, indicates the spread of the outcome distribution, reflecting the consistency of phase estimation across trials.
Figure \ref{fig:supp1} A shows the distribution of system delays over 10,000 simulated trials for in-phase and anti-phase stimulation, with the median delay (red line) at 16 ms for both conditions. Statistical comparison revealed no significant difference in delay between in- and anti-phase tACS (paired t-test: \( t(28)=1.648 \), \( p=0.110 \)), indicating consistent timing performance across stimulation modes.

Figure \ref{fig:supp1} B presents the accuracy of the phase prediction algorithm, depicted as the distribution of phase differences (in degrees) between the actual EEG phase at stimulation onset and the targeted phase, accounting for the measured 16 ms system delay. The clustering of phase differences near the target indicates the system’s reliable phase-locked stimulation capability, essential for accurate closed-loop tACS delivery.

\begin{figure}[h]
    \centering
    \includegraphics[width=0.85\textwidth]{Figures/Figs1_system.pdf}
    \caption{\textbf{Closed-loop system evaluation.} (\textbf{A}) Distribution of system delays measured over 10,000 trials for in-phase and anti-phase tACS. Median delay (red line) was 16 ms for both (\textbf{B}) Polar histogram of phase prediction accuracy, showing the distribution of phase differences between actual and targeted EEG phases (accounting for 16 ms delay), demonstrating precise phase-locking of the closed-loop tACS system.}
    \label{fig:supp1}
\end{figure}

\subsection*{Supplementary 2. Self-report discomfort scores}\label{supp2}

% \usepackage{multirow}
\begin{table}[h]
    \caption{Mean \(\pm\) standard deviation of self-reported discomfort scores (range: 0\textendash 100) after each stimulation condition. }
\begin{tabular}{lll}
\hline
Stimulation condition & Discomfort score (0-100) & Main effect of stimulation                          \\ \hline
In-phase              & 35.8±22.64               & \multirow{3}{*}{F(2, 68)=16.85, p\textless{}0.001*} \\ \cline{1-2}
Anti-phase            & 38.74±20.76              &                                                     \\ \cline{1-2}
Sham                  & 20.43±21.66              &                                                     \\ \hline
\end{tabular}
\vspace{1.6ex}
\caption*{\footnotesize A repeated measures ANOVA revealed a significant main effect of stimulation condition on perceived discomfort \( (F(2, 68) = 16.85, p < 0.001) \).}
\label{tab:supp1}
\end{table}

\begin{table}[h]
    \caption{Tukey’s HSD post-hoc comparisons of discomfort scores across stimulation conditions.}
\begin{tabular}{llllll}
\hline
group1     & group2   & Mean diff & p       & lower    & upper   \\ \hline
Anti-phase & In-phase & 2.9429    & 0.8378  & -9.3944  & 15.2801 \\ \hline
Anti-phase & Sham     & -15.3714  & 0.0105* & -27.7087 & -3.0342 \\ \hline
In-phase   & Sham     & -18.3143  & 0.0018* & -30.6515 & -5.977  \\ \hline
\end{tabular}
\vspace{1.6ex}
\caption*{\footnotesize Asterisks denote significance ($p < .05$). Sham was rated significantly lower in discomfort than both In-phase and Anti-phase, while no significant difference was found between In-phase and Anti-phase.}
\label{tab:supp2}
\end{table}

\subsection*{Supplementary 3.	Context violation and non-parametric integration estimates comparisons}\label{supp3}

\begin{figure}[h]
    \centering
    \includegraphics[width=0.7\textwidth]{Figures/Figs2_race.pdf}
    \caption{\textbf{Assessment of context independence.} GO-STOP RT: Go reaction time – Failed stop reaction time. The dataset did not show a severe violation of independence.} % \citep{Bissett2021}
    \label{fig:supp2}
\end{figure} 

Non-parametric estimated behavioral metrics of stop-signal performance are summarized in Table \ref{tab:supp3}. SSRT was estimated using the integration method with replacement of go omissions \citep{Verbruggen2019}. The staircase algorithm successfully maintained stopping accuracy near the target 50\%, with no significant main effect (Friedman test, \( \chi^2(2) = 2.9 \), \( p = 0.23 \)). The GORT differed significantly across conditions (ANOVA, \( F(2,68) = 4.04 \), \( p = 0.022 \)), with follow-up comparisons revealing faster responses in the sham vs. anti-phase condition (\( p = 0.03 \)) and slower responses in the in-phase vs. sham (\( p = 0.01 \)). No significant main effects were observed for SSRT or stop-signal delay (SSD), with respective ANOVA results of \( F(2,68) = 0.14 \) (\( p = 0.87 \)) and \( F(2,68) = 1.27 \) (\( p = 0.29 \)). Pairwise comparisons for these measures did not reveal any significant differences.

% Please add the following required packages to your document preamble:
\begin{table}[]
    \small                     % or \small if you prefer
    \setlength{\tabcolsep}{2.5pt}     % default is ~6pt
    \renewcommand{\arraystretch}{0.9} % default is 1.0
    \caption{Summary of descriptive statistics, main effects, and post hoc comparisons across tACS phase conditions for non-parametric estimates.}
\begin{tabular}{lllllllllll}
\hline
                   & \multicolumn{3}{c}{\textbf{Mean (SD)}}                                       & \multirow{2}{*}{\textbf{Main Effect (F/χ², p)}}             & \multicolumn{2}{c}{\textbf{In.- Anti.}} & \multicolumn{1}{c}{\textbf{In. – Sham}} &            & \multicolumn{2}{c}{\textbf{Anti. - Sham}} \\ \cline{1-4} \cline{6-11} 
\textbf{Parameter} & \textbf{In-phase} & \textbf{Anti-phase} & \textbf{Sham}                      &                                                             & \textbf{Main diff.}    & \textbf{p}     & \textbf{Main diff.}                     & \textbf{p} & \textbf{Main diff.}     & \textbf{p}      \\ \hline
P(Inhibition) †    & 54.29 (2.12)      & 53.94 (1.91)        & 53.31 (3.82)                       & \begin{tabular}[c]{@{}l@{}}χ²(2)=2.9, p=.23\end{tabular}  & 144.50                 & .28            & 105.00                                  & .05        & -129.00                 & .54             \\
RTGO (ms)          & 818.20 (12.68)    & 822.50 (13.11)      & 818.11 (13.62)                     & \textbf{F(2,68)=4.04, p=.022}                               & \textbf{-2.71}         & \textbf{.01}   & 0.05                                    & .96        & \textbf{2.30}           & \textbf{.03}    \\
SSRT (ms)          & 208.73 (16.94)    & 209.46 (13.54)      & 209.61 (15.43)                     & F(2,68)=0.14, p=.87                                         & -0.39                  & .70            & -0.52                                   & .61        & -0.08                   & .94             \\
SFRT (ms)          & 801.17 (16.41)    & 805.89 (15.44)      & \multicolumn{1}{c}{800.87 (17.52)} & F(2,68)=2.31, p=.11                                         & -1.82                  & .08           & 0.10                                    & .92       & \textbf{2.19}                    & \textbf{.04}            \\
SSD (ms)           & 604.87 (17.70)    & 608.54 (14.51)      & 606.03 (19.01)                     & F(2,68)=1.27, p=.29                                         & -1.43                  & .16            & -0.54                                   & .59        & 1.08                    & .29             \\
P(Go errors)       & 0                 & 0                   & 0                                  & NA                                                          & NA                     & NA             & NA                                      & NA         & NA                      & NA              \\
P(Go omissions)    & 2.49 (3.02)       & 2.40 (2.53)         & 2.34 (3.16)                        & \begin{tabular}[c]{@{}l@{}}χ²(2)=0.48, p=.79\end{tabular} & 201.00                 & .78           & 149.00                                  & .99       & -141.50                 & .82            \\ \hline
\end{tabular}
\vspace{1.6ex}
\caption*{\footnotesize Main effects and post-hoc results were tested using repeated measures ANOVA and paired t-tests. No corrections were applied to $p$ values.\\
† Friedman test with Wilcoxon signed-rank tests was conducted for non-normal distributed data.\\
SSRT=Stop signal reaction time, estimated using the integration method with replacement of go omissions \citep{Verbruggen2019}. RTGO=mean reaction time of go trials; SSRT=Stop-signal reaction time; RTSF=mean reaction time of Stop-failed trials; SSD=Stop-signal delay.}
\label{tab:supp3}
\end{table}

\subsection*{Supplementary 4. BEESTS-CV model prior distributions}

The prior distributions:
\begin{equation}
\begin{aligned}
\mu_{go} &\sim Normal(0.5, 0.50)[0,1], \\
\mu_{stop} &\sim Normal(0.2, 0.50)[0,1], \\
\sigma_{go}, \sigma_{stop} &\sim Normal(0.03, 0.35)[0, 1], \\
\tau_{go}, \tau_{stop} &\sim Normal(0.03, 0.35)[0, 1], \\
\phi^{-1}(P_{TF}), \phi^{-1}(P_{GF}) &\sim Normal(-1.28, 3), \\
M &\sim Normal(0, 3), \\
S &\sim Normal(1, 3)[0, \infty].
\end{aligned}
\end{equation}

The hyper-prior location parameter distributions:
\begin{equation}
\begin{aligned}
\mu_{\mu_{go}} &\sim Normal(0.5, 0.5)[0, 1], \\
\mu_{\mu_{stop}} &\sim Normal(0.2, 0.5)[0, 1], \\
\mu_{\sigma_{go}}, \mu_{\sigma_{stop}} &\sim Normal(0.04, 0.35)[0, \infty], \\
\mu_{\tau_{go}}, \mu_{\tau_{stop}} &\sim Normal(0.04, 0.35)[0, \infty], \\
\mu_{\phi^{-1}(P_{TF})}, \mu_{\phi^{-1}(P_{GF})} &\sim Normal(-1.28, 3)[-\infty, \infty], \\
\mu_{M} &\sim Normal(0, 3)[-\infty, \infty], \\
\mu_{S} &\sim Normal(0, 1)[0, \infty].
\end{aligned}
\end{equation}

The hyper-prior scale parameters:
\begin{equation}
\sigma_{p} \sim Exponential(1)
\end{equation}

For each parameter $p \in \{\mu_{go}, \mu_{stop}, \sigma_{go}, \sigma_{stop}, \tau_{go}, \tau_{stop}, P_{TF}, P_{GF}, M, S\}$.

\subsection*{Supplementary 5.	BEESTS-CV model fitting and context independence}

\begin{figure}[h]
    \centering
    \includegraphics[width=0.75\textwidth]{Figures/Figs4_cdf.pdf}
    \caption{Averaged observed and predicted cumulative distribution functions of reaction time (RTs) for go signal (left panels) and stop signals (right panels) collapsed (due to the SSDs determined by a staircase algorithm) across SSDs in three conditions. The thick dashed lines show the observed data. The grey dots show the prediction samples from the joint posterior distribution of the model parameters. The thin dashed lines show the average of the predictions.}
    \label{fig:supp4}
\end{figure}


\begin{figure}[h]
    \centering
    \includegraphics[width=0.75\textwidth]{Figures/Figs5_inhibition_function.pdf}
    \caption{Observed vs predicted inhibition functions (left panels) and median signal-respond RT as a function of stop-signal delay (SSD; right panels) for three stimulation conditions. Black bullets show the observed data. Grey violin plots show the distribution of the predictions of the BEESTS-CV model.}
    \label{fig:supp5}
\end{figure}
\newpage
\begin{figure}[h]
    \centering
    \includegraphics[width=0.75\textwidth]{Figures/Figs6_delta.pdf}
    \caption{Average delta functions in three conditions: in-, anti-phase and sham. The red line shows the observed delta functions averaged across participants. The black lines show 1000 posterior predictive delta functions averaged across participants obtained from fitting the data with the BEESTS-CV model assuming context independence; the darker the area, the denser the predicted delta function.}
    \label{fig:supp6}
\end{figure}

\FloatBarrier
\ifdefined\ismergeddocument
\else
\subsection*{References}
\bibliography{refs.bib}
\fi
