## Supplementary material for "Closed-loop phase-locked EEG-tACS enables adaptive control of cortical beta synchrony and motor control": Latex project: refs.bib

@article{Swann2009,
  title = {Intracranial EEG Reveals a Time- and Frequency-Specific Role for the Right Inferior Frontal Gyrus and Primary Motor Cortex in Stopping Initiated Responses},
  volume = {29},
  ISSN = {1529-2401},
  url = {http://dx.doi.org/10.1523/JNEUROSCI.3359-09.2009},
  DOI = {10.1523/jneurosci.3359-09.2009},
  number = {40},
  journal = {The Journal of Neuroscience},
  publisher = {Society for Neuroscience},
  author = {Swann,  Nicole and Tandon,  Nitin and Canolty,  Ryan and Ellmore,  Timothy M. and McEvoy,  Linda K. and Dreyer,  Stephen and DiSano,  Michael and Aron,  Adam R.},
  year = {2009},
  month = oct,
  pages = {12675-12685}
}

@article{Swann2012,
  title = {Roles for the pre-supplementary motor area and the right inferior frontal gyrus in stopping action: Electrophysiological responses and functional and structural connectivity},
  volume = {59},
  ISSN = {1053-8119},
  url = {http://dx.doi.org/10.1016/j.neuroimage.2011.09.049},
  DOI = {10.1016/j.neuroimage.2011.09.049},
  number = {3},
  journal = {NeuroImage},
  publisher = {Elsevier BV},
  author = {Swann,  Nicole C. and Cai,  Weidong and Conner,  Christopher R. and Pieters,  Thomas A. and Claffey,  Michael P. and George,  Jobi S. and Aron,  Adam R. and Tandon,  Nitin},
  year = {2012},
  month = feb,
  pages = {2860-2870}
}

@article{Wagner2018,
  title = {Establishing a Right Frontal Beta Signature for Stopping Action in Scalp EEG: Implications for Testing Inhibitory Control in Other Task Contexts},
  volume = {30},
  ISSN = {1530-8898},
  url = {http://dx.doi.org/10.1162/jocn_a_01183},
  DOI = {10.1162/jocn_a_01183},
  number = {1},
  journal = {Journal of Cognitive Neuroscience},
  publisher = {MIT Press - Journals},
  author = {Wagner,  Johanna and Wessel,  Jan R. and Ghahremani,  Ayda and Aron,  Adam R.},
  year = {2018},
  month = jan,
  pages = {107-118}
}

@article{Pfurtscheller1996,
  title = {Post-movement beta synchronization. A correlate of an idling motor area?},
  volume = {98},
  ISSN = {0013-4694},
  url = {http://dx.doi.org/10.1016/0013-4694(95)00258-8},
  DOI = {10.1016/0013-4694(95)00258-8},
  number = {4},
  journal = {Electroencephalography and Clinical Neurophysiology},
  publisher = {Elsevier BV},
  author = {Pfurtscheller,  G. and Stancák,  A. and Neuper,  C.},
  year = {1996},
  month = apr,
  pages = {281-293}
}

@article{Wessel2013,
  title = {Unexpected Events Induce Motor Slowing via a Brain Mechanism for Action-Stopping with Global Suppressive Effects},
  volume = {33},
  ISSN = {1529-2401},
  url = {http://dx.doi.org/10.1523/JNEUROSCI.3456-13.2013},
  DOI = {10.1523/jneurosci.3456-13.2013},
  number = {47},
  journal = {The Journal of Neuroscience},
  publisher = {Society for Neuroscience},
  author = {Wessel,  Jan R. and Aron,  Adam R.},
  year = {2013},
  month = nov,
  pages = {18481-18491}
}

@article{Alegre2013,
  title = {The subthalamic nucleus is involved in successful inhibition in the stop-signal task: A local field potential study in Parkinson's disease},
  volume = {239},
  ISSN = {0014-4886},
  url = {http://dx.doi.org/10.1016/j.expneurol.2012.08.027},
  DOI = {10.1016/j.expneurol.2012.08.027},
  journal = {Experimental Neurology},
  publisher = {Elsevier BV},
  author = {Alegre,  Manuel and Lopez-Azcarate,  Jon and Obeso,  Ignacio and Wilkinson,  Leonora and Rodriguez-Oroz,  Maria C. and Valencia,  Miguel and Garcia-Garcia,  David and Guridi,  Jorge and Artieda,  Julio and Jahanshahi,  Marjan and Obeso,  Jose A.},
  year = {2013},
  month = jan,
  pages = {1-12}
}

@article{Ray2012,
  title = {The role of the subthalamic nucleus in response inhibition: Evidence from local field potential recordings in the human subthalamic nucleus},
  volume = {60},
  ISSN = {1053-8119},
  url = {http://dx.doi.org/10.1016/j.neuroimage.2011.12.035},
  DOI = {10.1016/j.neuroimage.2011.12.035},
  number = {1},
  journal = {NeuroImage},
  publisher = {Elsevier BV},
  author = {Ray,  Nicola J. and Brittain,  John-Stuart and Holland,  Peter and Joundi,  Raed A. and Stein,  John F. and Aziz,  Tipu Z. and Jenkinson,  Ned},
  year = {2012},
  month = mar,
  pages = {271-278}
}

@article{Ray2008,
  title = {Local field potential beta activity in the subthalamic nucleus of patients with Parkinson's disease is associated with improvements in bradykinesia after dopamine and deep brain stimulation},
  volume = {213},
  ISSN = {0014-4886},
  url = {http://dx.doi.org/10.1016/j.expneurol.2008.05.008},
  DOI = {10.1016/j.expneurol.2008.05.008},
  number = {1},
  journal = {Experimental Neurology},
  publisher = {Elsevier BV},
  author = {Ray,  N.J. and Jenkinson,  N. and Wang,  S. and Holland,  P. and Brittain,  J.S. and Joint,  C. and Stein,  J.F. and Aziz,  T.},
  year = {2008},
  month = sep,
  pages = {108-113}
}

@article{Brittain2013,
  title = {Tremor Suppression by Rhythmic Transcranial Current Stimulation},
  volume = {23},
  ISSN = {0960-9822},
  url = {http://dx.doi.org/10.1016/j.cub.2013.01.068},
  DOI = {10.1016/j.cub.2013.01.068},
  number = {5},
  journal = {Current Biology},
  publisher = {Elsevier BV},
  author = {Brittain,  John-Stuart and Probert-Smith,  Penny and Aziz,  Tipu Z. and Brown,  Peter},
  year = {2013},
  month = mar,
  pages = {436-440}
}

@article{Brittain2014,
  title = {The highs and lows of beta activity in cortico‐basal ganglia loops},
  volume = {39},
  ISSN = {1460-9568},
  url = {http://dx.doi.org/10.1111/ejn.12574},
  DOI = {10.1111/ejn.12574},
  number = {11},
  journal = {European Journal of Neuroscience},
  publisher = {Wiley},
  author = {Brittain,  John‐Stuart and Sharott,  Andrew and Brown,  Peter},
  year = {2014},
  month = apr,
  pages = {1951-1959}
}

@article{Schaum2021,
  title = {Right inferior frontal gyrus implements motor inhibitory control via beta-band oscillations in humans},
  volume = {10},
  ISSN = {2050-084X},
  url = {http://dx.doi.org/10.7554/eLife.61679},
  DOI = {10.7554/elife.61679},
  journal = {eLife},
  publisher = {eLife Sciences Publications,  Ltd},
  author = {Schaum,  Michael and Pinzuti,  Edoardo and Sebastian,  Alexandra and Lieb,  Klaus and Fries,  Pascal and Mobascher,  Arian and Jung,  Patrick and Wibral,  Michael and T\"{u}scher,  Oliver},
  year = {2021},
  month = mar 
}

@article{Diesburg2021,
  title = {The Pause-then-Cancel model of human action-stopping: Theoretical considerations and empirical evidence},
  volume = {129},
  ISSN = {0149-7634},
  url = {http://dx.doi.org/10.1016/j.neubiorev.2021.07.019},
  DOI = {10.1016/j.neubiorev.2021.07.019},
  journal = {Neuroscience \& Biobehavioral Reviews},
  publisher = {Elsevier BV},
  author = {Diesburg,  Darcy A. and Wessel,  Jan R.},
  year = {2021},
  month = oct,
  pages = {17-34}
}

@article{Fonken2016,
  title = {Frontal and motor cortex contributions to response inhibition: evidence from electrocorticography},
  volume = {115},
  ISSN = {1522-1598},
  url = {http://dx.doi.org/10.1152/jn.00708.2015},
  DOI = {10.1152/jn.00708.2015},
  number = {4},
  journal = {Journal of Neurophysiology},
  publisher = {American Physiological Society},
  author = {Fonken,  Yvonne M. and Rieger,  Jochem W. and Tzvi,  Elinor and Crone,  Nathan E. and Chang,  Edward and Parvizi,  Josef and Knight,  Robert T. and Kr\"{a}mer,  Ulrike M.},
  year = {2016},
  month = apr,
  pages = {2224-2236}
}

@article{Leunissen2022,
  title = {Effects of beta-band and gamma-band rhythmic stimulation on motor inhibition},
  volume = {25},
  ISSN = {2589-0042},
  url = {http://dx.doi.org/10.1016/j.isci.2022.104338},
  DOI = {10.1016/j.isci.2022.104338},
  number = {5},
  journal = {iScience},
  publisher = {Elsevier BV},
  author = {Leunissen,  Inge and Van Steenkiste,  Manon and Heise,  Kirstin-Friederike and Monteiro,  Thiago Santos and Dunovan,  Kyle and Mantini,  Dante and Coxon,  James P. and Swinnen,  Stephan P.},
  year = {2022},
  month = may,
  pages = {104338}
}

@article{Kuhn2008,
  title = {High-Frequency Stimulation of the Subthalamic Nucleus Suppresses Oscillatory   Activity in Patients with Parkinson's Disease in Parallel with Improvement in Motor Performance},
  volume = {28},
  ISSN = {1529-2401},
  url = {http://dx.doi.org/10.1523/JNEUROSCI.0282-08.2008},
  DOI = {10.1523/jneurosci.0282-08.2008},
  number = {24},
  journal = {Journal of Neuroscience},
  publisher = {Society for Neuroscience},
  author = {Kuhn,  A. A. and Kempf,  F. and Brucke,  C. and Gaynor Doyle,  L. and Martinez-Torres,  I. and Pogosyan,  A. and Trottenberg,  T. and Kupsch,  A. and Schneider,  G.-H. and Hariz,  M. I. and Vandenberghe,  W. and Nuttin,  B. and Brown,  P.},
  year = {2008},
  month = jun,
  pages = {6165-6173}
}

@article{Little2012,
  title = {Beta band stability over time correlates with Parkinsonian rigidity and bradykinesia},
  volume = {236},
  ISSN = {0014-4886},
  url = {http://dx.doi.org/10.1016/j.expneurol.2012.04.024},
  DOI = {10.1016/j.expneurol.2012.04.024},
  number = {2},
  journal = {Experimental Neurology},
  publisher = {Elsevier BV},
  author = {Little,  S. and Pogosyan,  A. and Kuhn,  A.A. and Brown,  P.},
  year = {2012},
  month = aug,
  pages = {383-388}
}

@article{Chen2010,
  title = {Complexity of subthalamic 13-35Hz oscillatory activity directly correlates with clinical impairment in patients with Parkinson's disease},
  volume = {224},
  ISSN = {0014-4886},
  url = {http://dx.doi.org/10.1016/j.expneurol.2010.03.015},
  DOI = {10.1016/j.expneurol.2010.03.015},
  number = {1},
  journal = {Experimental Neurology},
  publisher = {Elsevier BV},
  author = {Chen,  Chiung Chu and Hsu,  Yi Ting and Chan,  Hsiao Lung and Chiou,  Shang Ming and Tu,  Po Hsun and Lee,  Shih Tseng and Tsai,  Chon Haw and Lu,  Chin Song and Brown,  Peter},
  year = {2010},
  month = jul,
  pages = {234-240}
}

@article{Toledo2014,
  title = {High beta activity in the subthalamic nucleus and freezing of gait in Parkinson's disease},
  volume = {64},
  ISSN = {0969-9961},
  url = {http://dx.doi.org/10.1016/j.nbd.2013.12.005},
  DOI = {10.1016/j.nbd.2013.12.005},
  journal = {Neurobiology of Disease},
  publisher = {Elsevier BV},
  author = {Toledo,  Jon B. and López-Azcárate,  Jon and Garcia-Garcia,  David and Guridi,  Jorge and Valencia,  Miguel and Artieda,  Julio and Obeso,  Jose and Alegre,  Manuel and Rodriguez-Oroz,  Maria},
  year = {2014},
  month = apr,
  pages = {60-65}
}

@article{Tiwari2012,
  title = {Evidences of possible side effects of neuroleptic drugs: A systematic review},
  volume = {1},
  ISSN = {2305-0500},
  url = {http://dx.doi.org/10.1016/S2305-0500(13)60105-0},
  DOI = {10.1016/s2305-0500(13)60105-0},
  number = {4},
  journal = {Asian Pacific Journal of Reproduction},
  publisher = {Asian Pacific Journal of Tropical Medicine Press},
  author = {Tiwari,  Prashant and Panik,  Rajnikant and Bhattacharya,  Arin and Ahirwar,  Dheeraj and Chandy,  Anish},
  year = {2012},
  month = dec,
  pages = {330-336}
}

@article{Chen2019,
  title = {Invasive and Noninvasive Brain Stimulation in Parkinson's Disease: Clinical Effects and Future Perspectives},
  volume = {106},
  ISSN = {1532-6535},
  url = {http://dx.doi.org/10.1002/cpt.1542},
  DOI = {10.1002/cpt.1542},
  number = {4},
  journal = {Clinical Pharmacology \& Therapeutics},
  publisher = {Wiley},
  author = {Chen,  Kai-Hsiang Stanley and Chen,  Robert},
  year = {2019},
  month = jul,
  pages = {763-775}
}

@article{Mogul2014,
  title = {Electrical Control of Epilepsy},
  volume = {16},
  ISSN = {1545-4274},
  url = {http://dx.doi.org/10.1146/annurev-bioeng-071813-104720},
  DOI = {10.1146/annurev-bioeng-071813-104720},
  number = {1},
  journal = {Annual Review of Biomedical Engineering},
  publisher = {Annual Reviews},
  author = {Mogul,  David J. and van Drongelen,  Wim},
  year = {2014},
  month = jul,
  pages = {483-504}
}

@article{Grover2021,
  title = {Synchronizing Brain Rhythms to Improve Cognition},
  volume = {72},
  ISSN = {1545-326X},
  url = {http://dx.doi.org/10.1146/annurev-med-060619-022857},
  DOI = {10.1146/annurev-med-060619-022857},
  number = {1},
  journal = {Annual Review of Medicine},
  publisher = {Annual Reviews},
  author = {Grover,  Shrey and Nguyen,  John A. and Reinhart,  Robert M.G.},
  year = {2021},
  month = jan,
  pages = {29-43}
}

@article{Ozen2010,
  title = {Transcranial Electric Stimulation Entrains Cortical Neuronal Populations in Rats},
  volume = {30},
  ISSN = {1529-2401},
  url = {http://dx.doi.org/10.1523/jneurosci.5252-09.2010},
  DOI = {10.1523/jneurosci.5252-09.2010},
  number = {34},
  journal = {The Journal of Neuroscience},
  publisher = {Society for Neuroscience},
  author = {Ozen,  Simal and Sirota,  Anton and Belluscio,  Mariano A. and Anastassiou,  Costas A. and Stark,  Eran and Koch,  Christof and Buzsáki,  Gy\"{o}rgy},
  year = {2010},
  month = aug,
  pages = {11476-11485}
}

@article{Johnson2020,
  title = {Dose-dependent effects of transcranial alternating current stimulation on spike timing in awake nonhuman primates},
  volume = {6},
  ISSN = {2375-2548},
  url = {http://dx.doi.org/10.1126/sciadv.aaz2747},
  DOI = {10.1126/sciadv.aaz2747},
  number = {36},
  journal = {Science Advances},
  publisher = {American Association for the Advancement of Science (AAAS)},
  author = {Johnson,  Luke and Alekseichuk,  Ivan and Krieg,  Jordan and Doyle,  Alex and Yu,  Ying and Vitek,  Jerrold and Johnson,  Matthew and Opitz,  Alexander},
  year = {2020},
  month = sep 
}

@article{Krause2019,
  title = {Transcranial alternating current stimulation entrains single-neuron activity in the primate brain},
  volume = {116},
  ISSN = {1091-6490},
  url = {http://dx.doi.org/10.1073/pnas.1815958116},
  DOI = {10.1073/pnas.1815958116},
  number = {12},
  journal = {Proceedings of the National Academy of Sciences},
  publisher = {Proceedings of the National Academy of Sciences},
  author = {Krause,  Matthew R. and Vieira,  Pedro G. and Csorba,  Bennett A. and Pilly,  Praveen K. and Pack,  Christopher C.},
  year = {2019},
  month = mar,
  pages = {5747-5755}
}

@article{Krause2023,
  title = {Transcranial electrical stimulation: How can a simple conductor orchestrate complex brain activity?},
  volume = {21},
  ISSN = {1545-7885},
  url = {http://dx.doi.org/10.1371/journal.pbio.3001973},
  DOI = {10.1371/journal.pbio.3001973},
  number = {1},
  journal = {PLOS Biology},
  publisher = {Public Library of Science (PLoS)},
  author = {Krause,  Matthew R. and Vieira,  Pedro G. and Pack,  Christopher C.},
  year = {2023},
  month = jan,
  pages = {e3001973}
}

@article{Thut2011,
  title = {Entrainment of Perceptually Relevant Brain Oscillations by Non-Invasive Rhythmic Stimulation of the Human Brain},
  volume = {2},
  ISSN = {1664-1078},
  url = {http://dx.doi.org/10.3389/fpsyg.2011.00170},
  DOI = {10.3389/fpsyg.2011.00170},
  journal = {Frontiers in Psychology},
  publisher = {Frontiers Media SA},
  author = {Thut,  Gregor and Schyns,  Philippe G. and Gross,  Joachim},
  year = {2011}
}

@article{Herrmann2016,
  title = {Shaping Intrinsic Neural Oscillations with Periodic Stimulation},
  volume = {36},
  ISSN = {1529-2401},
  url = {http://dx.doi.org/10.1523/JNEUROSCI.0236-16.2016},
  DOI = {10.1523/jneurosci.0236-16.2016},
  number = {19},
  journal = {The Journal of Neuroscience},
  publisher = {Society for Neuroscience},
  author = {Herrmann,  Christoph S. and Murray,  Micah M. and Ionta,  Silvio and Hutt,  Axel and Lefebvre,  Jérémie},
  year = {2016},
  month = may,
  pages = {5328-5337}
}

@article{Pogosyan2009,
  title = {Boosting Cortical Activity at Beta-Band Frequencies Slows Movement in Humans},
  volume = {19},
  ISSN = {0960-9822},
  url = {http://dx.doi.org/10.1016/j.cub.2009.07.074},
  DOI = {10.1016/j.cub.2009.07.074},
  number = {19},
  journal = {Current Biology},
  publisher = {Elsevier BV},
  author = {Pogosyan,  Alek and Gaynor,  Louise Doyle and Eusebio,  Alexandre and Brown,  Peter},
  year = {2009},
  month = oct,
  pages = {1637-1641}
}

@article{Joundi2012,
  title = {Driving Oscillatory Activity in the Human Cortex Enhances Motor Performance},
  volume = {22},
  ISSN = {0960-9822},
  url = {http://dx.doi.org/10.1016/j.cub.2012.01.024},
  DOI = {10.1016/j.cub.2012.01.024},
  number = {5},
  journal = {Current Biology},
  publisher = {Elsevier BV},
  author = {Joundi,  Raed A. and Jenkinson,  Ned and Brittain,  John-Stuart and Aziz,  Tipu Z. and Brown,  Peter},
  year = {2012},
  month = mar,
  pages = {403-407}
}

@article{Engel2010,
  title = {Beta-band oscillations — signalling the status quo?},
  volume = {20},
  ISSN = {0959-4388},
  url = {http://dx.doi.org/10.1016/j.conb.2010.02.015},
  DOI = {10.1016/j.conb.2010.02.015},
  number = {2},
  journal = {Current Opinion in Neurobiology},
  publisher = {Elsevier BV},
  author = {Engel,  Andreas K and Fries,  Pascal},
  year = {2010},
  month = apr,
  pages = {156-165}
}

@article{Kilavik2013,
  title = {The ups and downs of beta oscillations in sensorimotor cortex},
  volume = {245},
  ISSN = {0014-4886},
  url = {http://dx.doi.org/10.1016/j.expneurol.2012.09.014},
  DOI = {10.1016/j.expneurol.2012.09.014},
  journal = {Experimental Neurology},
  publisher = {Elsevier BV},
  author = {Kilavik,  Bjørg Elisabeth and Zaepffel,  Manuel and Brovelli,  Andrea and MacKay,  William A. and Riehle,  Alexa},
  year = {2013},
  month = jul,
  pages = {15-26}
}

@article{Spitzer2017,
  title = {Beyond the Status Quo: A Role for Beta Oscillations in Endogenous Content (Re)Activation},
  volume = {4},
  ISSN = {2373-2822},
  url = {http://dx.doi.org/10.1523/ENEURO.0170-17.2017},
  DOI = {10.1523/eneuro.0170-17.2017},
  number = {4},
  journal = {eneuro},
  publisher = {Society for Neuroscience},
  author = {Spitzer,  Bernhard and Haegens,  Saskia},
  year = {2017},
  month = jul,
  pages = {ENEURO.0170--17.2017}
}

@article{Barone2021,
  title = {Understanding the Role of Sensorimotor Beta Oscillations},
  volume = {15},
  ISSN = {1662-5137},
  url = {http://dx.doi.org/10.3389/fnsys.2021.655886},
  DOI = {10.3389/fnsys.2021.655886},
  journal = {Frontiers in Systems Neuroscience},
  publisher = {Frontiers Media SA},
  author = {Barone,  Jacopo and Rossiter,  Holly E.},
  year = {2021},
  month = may 
}

@article{Tan2022,
  title = {The Effects of Dual-Site Beta tACS over the rIFG and preSMA on Response Inhibition in Young and Older Adults},
  url = {http://dx.doi.org/10.1101/2022.11.30.518460},
  DOI = {10.1101/2022.11.30.518460},
  journal = {bioRxiv},
  publisher = {Cold Spring Harbor Laboratory},
  author = {Tan,  Jane and Iyer,  Kartik K. and Nitsche,  Michael A. and Puri,  Rohan and Hinder,  Mark R. and Fujiyama,  Hakuei},
  year = {2022},
  month = dec 
}

@article{Weinberger2006,
  title = {Beta Oscillatory Activity in the Subthalamic Nucleus and Its Relation to Dopaminergic Response in Parkinson's Disease},
  volume = {96},
  ISSN = {1522-1598},
  url = {http://dx.doi.org/10.1152/jn.00697.2006},
  DOI = {10.1152/jn.00697.2006},
  number = {6},
  journal = {Journal of Neurophysiology},
  publisher = {American Physiological Society},
  author = {Weinberger,  Moran and Mahant,  Neil and Hutchison,  William D. and Lozano,  Andres M. and Moro,  Elena and Hodaie,  Mojgan and Lang,  Anthony E. and Dostrovsky,  Jonathan O.},
  year = {2006},
  month = dec,
  pages = {3248-3256}
}

@article{Lofredi2023,
  title = {Subthalamic beta bursts correlate with dopamine-dependent motor symptoms in 106 Parkinson's patients},
  volume = {9},
  ISSN = {2373-8057},
  url = {http://dx.doi.org/10.1038/s41531-022-00443-3},
  DOI = {10.1038/s41531-022-00443-3},
  number = {1},
  journal = {npj Parkinson's Disease},
  publisher = {Springer Science and Business Media LLC},
  author = {Lofredi,  Roxanne and Okudzhava,  Liana and Irmen,  Friederike and Br\"{u}cke,  Christof and Huebl,  Julius and Krauss,  Joachim K. and Schneider,  Gerd-Helge and Faust,  Katharina and Neumann,  Wolf-Julian and K\"{u}hn,  Andrea A.},
  year = {2023},
  month = jan 
}

@article{Krause2022,
  title = {Brain stimulation competes with ongoing oscillations for control of spike timing in the primate brain},
  volume = {20},
  ISSN = {1545-7885},
  url = {http://dx.doi.org/10.1371/journal.pbio.3001650},
  DOI = {10.1371/journal.pbio.3001650},
  number = {5},
  journal = {PLOS Biology},
  publisher = {Public Library of Science (PLoS)},
  author = {Krause,  Matthew R. and Vieira,  Pedro G. and Thivierge,  Jean-Philippe and Pack,  Christopher C.},
  editor = {Luo,  Huan},
  year = {2022},
  month = may,
  pages = {e3001650}
}

@article{Karabanov2016,
  title = {Transcranial brain stimulation: closing the loop between brain and stimulation},
  volume = {29},
  ISSN = {1473-6551},
  url = {http://dx.doi.org/10.1097/WCO.0000000000000342},
  DOI = {10.1097/wco.0000000000000342},
  number = {4},
  journal = {Current Opinion in Neurology},
  publisher = {Ovid Technologies (Wolters Kluwer Health)},
  author = {Karabanov,  Anke and Thielscher,  Axel and Siebner,  Hartwig Roman},
  year = {2016},
  month = aug,
  pages = {397-404}
}

@article{Brown2007,
  title = {Abnormal oscillatory synchronisation in the motor system leads to impaired movement},
  volume = {17},
  ISSN = {0959-4388},
  url = {http://dx.doi.org/10.1016/j.conb.2007.12.001},
  DOI = {10.1016/j.conb.2007.12.001},
  number = {6},
  journal = {Current Opinion in Neurobiology},
  publisher = {Elsevier BV},
  author = {Brown,  P},
  year = {2007},
  month = dec,
  pages = {656-664}
}

@article{Kothe2025,
  title     = {The Lab Streaming Layer for Synchronized Multimodal Recording},
  author    = {Kothe, Christian and Shirazi, Seyed Yahya and Stenner, Tristan and Medine, David and Boulay, Chadwick and Grivich, Matthew I. and Artoni, Fiorenzo and Mullen, Tim and Delorme, Arnaud and Makeig, Scott},
  journal   = {Imaging Neuroscience},
  volume    = {3},
  pages     = {IMAG.a.136},
  year      = {2025},
  doi       = {10.1162/IMAG.a.136},
  url       = {https://doi.org/10.1162/IMAG.a.136},
  publisher = {MIT Press}
}


@article{Chen2023,
  title = {Alpha oscillatory activity is causally linked to working memory retention},
  volume = {21},
  ISSN = {1545-7885},
  url = {http://dx.doi.org/10.1371/journal.pbio.3001999},
  DOI = {10.1371/journal.pbio.3001999},
  number = {2},
  journal = {PLOS Biology},
  publisher = {Public Library of Science (PLoS)},
  author = {Chen,  Xueli and Ma,  Ru and Zhang,  Wei and Zeng,  Ginger Qinghong and Wu,  Qianying and Yimiti,  Ajiguli and Xia,  Xinzhao and Cui,  Jiangtian and Liu,  Qiongwei and Meng,  Xueer and Bu,  Junjie and Chen,  Qi and Pan,  Yu and Yu,  Nancy Xiaonan and Wang,  Shouyan and Deng,  Zhi-De and Sack,  Alexander T. and Laughlin,  Myles Mc and Zhang,  Xiaochu},
  editor = {Hanslmayr,  Simon},
  year = {2023},
  month = feb,
  pages = {e3001999}
}

@article{SlaterHammel1960,
  title = {Reliability,  Accuracy,  and Refractoriness of a Transit Reaction},
  volume = {31},
  ISSN = {1067-1188},
  url = {http://dx.doi.org/10.1080/10671188.1960.10613098},
  DOI = {10.1080/10671188.1960.10613098},
  number = {2},
  journal = {Research Quarterly. American Association for Health,  Physical Education and Recreation},
  publisher = {Informa UK Limited},
  author = {Slater-Hammel,  A. T.},
  year = {1960},
  month = may,
  pages = {217-228}
}

@article{Coxon2006,
  title = {Intracortical Inhibition During Volitional Inhibition of Prepared Action},
  volume = {95},
  ISSN = {1522-1598},
  url = {http://dx.doi.org/10.1152/jn.01334.2005},
  DOI = {10.1152/jn.01334.2005},
  number = {6},
  journal = {Journal of Neurophysiology},
  publisher = {American Physiological Society},
  author = {Coxon,  James P. and Stinear,  Cathy M. and Byblow,  Winston D.},
  year = {2006},
  month = jun,
  pages = {3371-3383}
}

@article{Verbruggen2019,
  title = {A consensus guide to capturing the ability to inhibit actions and impulsive behaviors in the stop-signal task},
  volume = {8},
  ISSN = {2050-084X},
  url = {http://dx.doi.org/10.7554/eLife.46323},
  DOI = {10.7554/elife.46323},
  journal = {eLife},
  publisher = {eLife Sciences Publications,  Ltd},
  author = {Verbruggen,  Frederick and Aron,  Adam R and Band,  Guido PH and Beste,  Christian and Bissett,  Patrick G and Brockett,  Adam T and Brown,  Joshua W and Chamberlain,  Samuel R and Chambers,  Christopher D and Colonius,  Hans and Colzato,  Lorenza S and Corneil,  Brian D and Coxon,  James P and Dupuis,  Annie and Eagle,  Dawn M and Garavan,  Hugh and Greenhouse,  Ian and Heathcote,  Andrew and Huster,  René J and Jahfari,  Sara and Kenemans,  J Leon and Leunissen,  Inge and Li,  Chiang-Shan R and Logan,  Gordon D and Matzke,  Dora and Morein-Zamir,  Sharon and Murthy,  Aditya and Paré,  Martin and Poldrack,  Russell A and Ridderinkhof,  K Richard and Robbins,  Trevor W and Roesch,  Matthew and Rubia,  Katya and Schachar,  Russell J and Schall,  Jeffrey D and Stock,  Ann-Kathrin and Swann,  Nicole C and Thakkar,  Katharine N and van der Molen,  Maurits W and Vermeylen,  Luc and Vink,  Matthijs and Wessel,  Jan R and Whelan,  Robert and Zandbelt,  Bram B and Boehler,  C Nico},
  year = {2019},
  month = apr 
}

@article{Onojima2021,
  title = {A closed-loop stimulation approach with real-time estimation of the instantaneous phase of neural oscillations by a Kalman filter},
  url = {http://dx.doi.org/10.1101/2021.04.25.441309},
  DOI = {10.1101/2021.04.25.441309},
  journal = {bioRxiv},
  publisher = {Cold Spring Harbor Laboratory},
  author = {Onojima,  Takayuki and Kitajo,  Keiichi},
  year = {2021},
  month = apr 
}

@article{Shirinpour2020,
  title = {Experimental evaluation of methods for real-time EEG phase-specific transcranial magnetic stimulation},
  volume = {17},
  ISSN = {1741-2552},
  url = {http://dx.doi.org/10.1088/1741-2552/ab9dba},
  DOI = {10.1088/1741-2552/ab9dba},
  number = {4},
  journal = {Journal of Neural Engineering},
  publisher = {IOP Publishing},
  author = {Shirinpour,  Sina and Alekseichuk,  Ivan and Mantell,  Kathleen and Opitz,  Alexander},
  year = {2020},
  month = jul,
  pages = {046002}
}

@article{Bikson2009,
  title = {Establishing safety limits for transcranial direct current stimulation},
  volume = {120},
  ISSN = {1388-2457},
  url = {http://dx.doi.org/10.1016/j.clinph.2009.03.018},
  DOI = {10.1016/j.clinph.2009.03.018},
  number = {6},
  journal = {Clinical Neurophysiology},
  publisher = {Elsevier BV},
  author = {Bikson,  Marom and Datta,  Abhishek and Elwassif,  Maged},
  year = {2009},
  month = jun,
  pages = {1033-1034}
}

@article{Woods2016,
  title = {A technical guide to tDCS,  and related non-invasive brain stimulation tools},
  volume = {127},
  ISSN = {1388-2457},
  url = {http://dx.doi.org/10.1016/j.clinph.2015.11.012},
  DOI = {10.1016/j.clinph.2015.11.012},
  number = {2},
  journal = {Clinical Neurophysiology},
  publisher = {Elsevier BV},
  author = {Woods,  A.J. and Antal,  A. and Bikson,  M. and Boggio,  P.S. and Brunoni,  A.R. and Celnik,  P. and Cohen,  L.G. and Fregni,  F. and Herrmann,  C.S. and Kappenman,  E.S. and Knotkova,  H. and Liebetanz,  D. and Miniussi,  C. and Miranda,  P.C. and Paulus,  W. and Priori,  A. and Reato,  D. and Stagg,  C. and Wenderoth,  N. and Nitsche,  M.A.},
  year = {2016},
  month = feb,
  pages = {1031-1048}
}

@article{Leunissen2017,
  title = {Reliable estimation of inhibitory efficiency: to anticipate,  choose or simply react?},
  volume = {45},
  ISSN = {1460-9568},
  url = {http://dx.doi.org/10.1111/ejn.13590},
  DOI = {10.1111/ejn.13590},
  number = {12},
  journal = {European Journal of Neuroscience},
  publisher = {Wiley},
  author = {Leunissen,  Inge and Zandbelt,  Bram B. and Potocanac,  Zrinka and Swinnen,  Stephan P. and Coxon,  James P.},
  editor = {Rossell,  Susan},
  year = {2017},
  month = jun,
  pages = {1512-1523}
}

@article{He2021,
  title = {OSARI,  an Open-Source Anticipated Response Inhibition Task},
  volume = {54},
  ISSN = {1554-3528},
  url = {http://dx.doi.org/10.3758/s13428-021-01680-9},
  DOI = {10.3758/s13428-021-01680-9},
  number = {3},
  journal = {Behavior Research Methods},
  publisher = {Springer Science and Business Media LLC},
  author = {He,  Jason L. and Hirst,  Rebecca J. and Puri,  Rohan and Coxon,  James and Byblow,  Winston and Hinder,  Mark and Skippen,  Patrick and Matzke,  Dora and Heathcote,  Andrew and Wadsley,  Corey G. and Silk,  Tim and Hyde,  Christian and Parmar,  Dinisha and Pedapati,  Ernest and Gilbert,  Donald L. and Huddleston,  David A. and Mostofsky,  Stewart and Leunissen,  Inge and MacDonald,  Hayley J. and Chowdhury,  Nahian S. and Gretton,  Matthew and Nikitenko,  Tess and Zandbelt,  Bram and Strickland,  Luke and Puts,  Nicolaas A. J.},
  year = {2021},
  month = nov,
  pages = {1530-1540}
}


@article{Peirce2019,
  title = {PsychoPy2: Experiments in behavior made easy},
  volume = {51},
  ISSN = {1554-3528},
  url = {http://dx.doi.org/10.3758/s13428-018-01193-y},
  DOI = {10.3758/s13428-018-01193-y},
  number = {1},
  journal = {Behavior Research Methods},
  publisher = {Springer Science and Business Media LLC},
  author = {Peirce,  Jonathan and Gray,  Jeremy R. and Simpson,  Sol and MacAskill,  Michael and H\"{o}chenberger,  Richard and Sogo,  Hiroyuki and Kastman,  Erik and Lindeløv,  Jonas Kristoffer},
  year = {2019},
  month = feb,
  pages = {195-203}
}

@article{Oostenveld2011,
  title = {FieldTrip: Open Source Software for Advanced Analysis of MEG,  EEG,  and Invasive Electrophysiological Data},
  volume = {2011},
  ISSN = {1687-5273},
  url = {http://dx.doi.org/10.1155/2011/156869},
  DOI = {10.1155/2011/156869},
  journal = {Computational Intelligence and Neuroscience},
  publisher = {Wiley},
  author = {Oostenveld,  Robert and Fries,  Pascal and Maris,  Eric and Schoffelen,  Jan-Mathijs},
  year = {2011},
  pages = {1-9}
}

@article{Mansouri2017,
  title = {A Fast EEG Forecasting Algorithm for Phase-Locked Transcranial Electrical Stimulation of the Human Brain},
  volume = {11},
  ISSN = {1662-453X},
  url = {http://dx.doi.org/10.3389/fnins.2017.00401},
  DOI = {10.3389/fnins.2017.00401},
  journal = {Frontiers in Neuroscience},
  publisher = {Frontiers Media SA},
  author = {Mansouri,  Farrokh and Dunlop,  Katharine and Giacobbe,  Peter and Downar,  Jonathan and Zariffa,  José},
  year = {2017},
  month = jul 
}

@article{Shakeel2020,
  title = {Time-Series Prediction of the Oscillatory Phase of EEG Signals Using the Least Mean Square Algorithm-Based AR Model},
  volume = {10},
  ISSN = {2076-3417},
  url = {http://dx.doi.org/10.3390/app10103616},
  DOI = {10.3390/app10103616},
  number = {10},
  journal = {Applied Sciences},
  publisher = {MDPI AG},
  author = {Shakeel,  Aqsa and Tanaka,  Toshihisa and Kitajo,  Keiichi},
  year = {2020},
  month = may,
  pages = {3616}
}

@article{Zrenner2018,
  title = {Real-time EEG-defined excitability states determine efficacy of TMS-induced plasticity in human motor cortex},
  volume = {11},
  ISSN = {1935-861X},
  url = {http://dx.doi.org/10.1016/j.brs.2017.11.016},
  DOI = {10.1016/j.brs.2017.11.016},
  number = {2},
  journal = {Brain Stimulation},
  publisher = {Elsevier BV},
  author = {Zrenner,  Christoph and Desideri,  Debora and Belardinelli,  Paolo and Ziemann,  Ulf},
  year = {2018},
  month = mar,
  pages = {374-389}
}

@article{Schevernels2015,
  title = {Electrophysiological evidence for the involvement of proactive and reactive control in a rewarded stop-signal task},
  volume = {121},
  ISSN = {1053-8119},
  url = {http://dx.doi.org/10.1016/j.neuroimage.2015.07.023},
  DOI = {10.1016/j.neuroimage.2015.07.023},
  journal = {NeuroImage},
  publisher = {Elsevier BV},
  author = {Schevernels,  Hanne and Bombeke,  Klaas and Van der Borght,  Liesbet and Hopf,  Jens-Max and Krebs,  Ruth M. and Boehler,  C. Nicolas},
  year = {2015},
  month = nov,
  pages = {115-125}
}

@article{Liebrand2017,
  title = {Temporal Dynamics of Proactive and Reactive Motor Inhibition},
  volume = {11},
  ISSN = {1662-5161},
  url = {http://dx.doi.org/10.3389/fnhum.2017.00204},
  DOI = {10.3389/fnhum.2017.00204},
  journal = {Frontiers in Human Neuroscience},
  publisher = {Frontiers Media SA},
  author = {Liebrand,  Matthias and Pein,  Inga and Tzvi,  Elinor and Kr\"{a}mer,  Ulrike M.},
  year = {2017},
  month = apr 
}

@article{Wessel2019,
  title = {beta-Bursts Reveal the Trial-to-Trial Dynamics of Movement Initiation and Cancellation},
  volume = {40},
  ISSN = {1529-2401},
  url = {http://dx.doi.org/10.1523/JNEUROSCI.1887-19.2019},
  DOI = {10.1523/jneurosci.1887-19.2019},
  number = {2},
  journal = {The Journal of Neuroscience},
  publisher = {Society for Neuroscience},
  author = {Wessel,  Jan R.},
  year = {2019},
  month = nov,
  pages = {411-423}
}

@article{Cardellicchio2020,
  title = {Beta Rebound as an Index of Temporal Integration of Somatosensory and Motor Signals},
  volume = {14},
  ISSN = {1662-5137},
  url = {http://dx.doi.org/10.3389/fnsys.2020.00063},
  DOI = {10.3389/fnsys.2020.00063},
  journal = {Frontiers in Systems Neuroscience},
  publisher = {Frontiers Media SA},
  author = {Cardellicchio,  Pasquale and Hilt,  Pauline M. and Dolfini,  Elisa and Fadiga,  Luciano and D'Ausilio,  Alessandro},
  year = {2020},
  month = sep 
}

@article{Haegens2014,
  title = {Inter- and intra-individual variability in alpha peak frequency},
  volume = {92},
  ISSN = {1053-8119},
  url = {http://dx.doi.org/10.1016/j.neuroimage.2014.01.049},
  DOI = {10.1016/j.neuroimage.2014.01.049},
  journal = {NeuroImage},
  publisher = {Elsevier BV},
  author = {Haegens,  Saskia and Cousijn,  Helena and Wallis,  George and Harrison,  Paul J. and Nobre,  Anna C.},
  year = {2014},
  month = may,
  pages = {46-55}
}

@article{Nikulin2006,
  title = {Phase synchronization between alpha and beta oscillations in the human electroencephalogram},
  volume = {137},
  ISSN = {0306-4522},
  url = {http://dx.doi.org/10.1016/j.neuroscience.2005.10.031},
  DOI = {10.1016/j.neuroscience.2005.10.031},
  number = {2},
  journal = {Neuroscience},
  publisher = {Elsevier BV},
  author = {Nikulin,  V.V. and Brismar,  T.},
  year = {2006},
  month = jan,
  pages = {647-657}
}

@article{Fang2024,
  title = {The phase of tACS-entrained pre-SMA beta oscillations modulates motor inhibition},
  volume = {290},
  ISSN = {1053-8119},
  url = {http://dx.doi.org/10.1016/j.neuroimage.2024.120572},
  DOI = {10.1016/j.neuroimage.2024.120572},
  journal = {NeuroImage},
  publisher = {Elsevier BV},
  author = {Fang,  Zhou and Sack,  Alexander T. and Leunissen,  Inge},
  year = {2024},
  month = apr,
  pages = {120572}
}

@article{Bissett2021,
  title = {Design issues and solutions for stop-signal data from the Adolescent Brain Cognitive Development (ABCD) study},
  volume = {10},
  ISSN = {2050-084X},
  url = {http://dx.doi.org/10.7554/eLife.60185},
  DOI = {10.7554/elife.60185},
  journal = {eLife},
  publisher = {eLife Sciences Publications,  Ltd},
  author = {Bissett,  Patrick G and Hagen,  McKenzie P and Jones,  Henry M and Poldrack,  Russell A},
  year = {2021},
  month = mar 
}

@article{Matzke2013,
  title = {Bayesian parametric estimation of stop-signal reaction time distributions.},
  volume = {142},
  ISSN = {0096-3445},
  url = {http://dx.doi.org/10.1037/a0030543},
  DOI = {10.1037/a0030543},
  number = {4},
  journal = {Journal of Experimental Psychology: General},
  publisher = {American Psychological Association (APA)},
  author = {Matzke,  Dora and Dolan,  Conor V. and Logan,  Gordon D. and Brown,  Scott D. and Wagenmakers,  Eric-Jan},
  year = {2013},
  month = nov,
  pages = {1047-1073}
}

@article{Matzke2016,
  title = {A Bayesian approach for estimating the probability of trigger failures in the stop-signal paradigm},
  volume = {49},
  ISSN = {1554-3528},
  url = {http://dx.doi.org/10.3758/s13428-015-0695-8},
  DOI = {10.3758/s13428-015-0695-8},
  number = {1},
  journal = {Behavior Research Methods},
  publisher = {Springer Science and Business Media LLC},
  author = {Matzke,  Dora and Love,  Jonathon and Heathcote,  Andrew},
  year = {2016},
  month = jan,
  pages = {267-281}
}

@article{Matzke2019,
  title = {Inhibiting responses to difficult choices.},
  volume = {148},
  ISSN = {0096-3445},
  url = {http://dx.doi.org/10.1037/xge0000525},
  DOI = {10.1037/xge0000525},
  number = {1},
  journal = {Journal of Experimental Psychology: General},
  publisher = {American Psychological Association (APA)},
  author = {Matzke,  Dora and Curley,  Samuel and Gong,  Charlene Q. and Heathcote,  Andrew},
  year = {2019},
  month = jan,
  pages = {124-142}
}

@article{Matzke2021,
  title = {Stopping timed actions.},
  url = {http://dx.doi.org/10.31234/osf.io/9h3v7},
  DOI = {10.31234/osf.io/9h3v7},
  journal = {OSF Preprints},
  publisher = {Center for Open Science},
  author = {Matzke,  Dora and Strickland,  Luke Joseph Gough and Sripada,  Chandra and Weigard,  Alexander Samuel and Puri,  Rohan and He,  Jason and Hirst,  Rebecca and Heathcote,  Andrew},
  year = {2021},
  month = mar 
}

@book{Ornstein1969,
  title     = {On the Experience of Time},
  author    = {Ornstein, Robert Evan},
  year      = {1969},
  publisher = {Penguin Books},
  address   = {Harmondsworth, England}
}

@article{Heathcote2018,
  title = {Dynamic models of choice},
  volume = {51},
  ISSN = {1554-3528},
  url = {http://dx.doi.org/10.3758/s13428-018-1067-y},
  DOI = {10.3758/s13428-018-1067-y},
  number = {2},
  journal = {Behavior Research Methods},
  publisher = {Springer Science and Business Media LLC},
  author = {Heathcote,  Andrew and Lin,  Yi-Shin and Reynolds,  Angus and Strickland,  Luke and Gretton,  Matthew and Matzke,  Dora},
  year = {2018},
  month = jun,
  pages = {961-985}
}

@article{Braak2006,
  title = {A Markov Chain Monte Carlo version of the genetic algorithm Differential Evolution: easy Bayesian computing for real parameter spaces},
  volume = {16},
  ISSN = {1573-1375},
  url = {http://dx.doi.org/10.1007/s11222-006-8769-1},
  DOI = {10.1007/s11222-006-8769-1},
  number = {3},
  journal = {Statistics and Computing},
  publisher = {Springer Science and Business Media LLC},
  author = {Braak,  Cajo J. F. Ter},
  year = {2006},
  month = sep,
  pages = {239-249}
}

@manual{Pinheiro1999,
  title = {nlme: Linear and Nonlinear Mixed Effects Models},
  url = {http://dx.doi.org/10.32614/CRAN.package.nlme},
  DOI = {10.32614/cran.package.nlme},
  journal = {CRAN: Contributed Packages},
  publisher = {The R Foundation},
  author = {Pinheiro,  Jos{\'e} and Bates,  Douglas},
  year = {1999},
  month = nov 
}

@article{Hothorn2008,
  title = {Simultaneous Inference in General Parametric Models},
  volume = {50},
  ISSN = {1521-4036},
  url = {http://dx.doi.org/10.1002/bimj.200810425},
  DOI = {10.1002/bimj.200810425},
  number = {3},
  journal = {Biometrical Journal},
  publisher = {Wiley},
  author = {Hothorn,  Torsten and Bretz,  Frank and Westfall,  Peter},
  year = {2008},
  month = jun,
  pages = {346-363}
}

@article{Choo2022,
  title = {Right inferior frontal gyrus damage is associated with impaired initiation of inhibitory control,  but not its implementation},
  volume = {11},
  ISSN = {2050-084X},
  url = {http://dx.doi.org/10.7554/eLife.79667},
  DOI = {10.7554/elife.79667},
  journal = {eLife},
  publisher = {eLife Sciences Publications,  Ltd},
  author = {Choo,  Yoojeong and Matzke,  Dora and Bowren,  Mark D and Tranel,  Daniel and Wessel,  Jan R},
  year = {2022},
  month = dec 
}

@article{SolisEscalante2012,
  title = {Cue-induced beta rebound during withholding of overt and covert foot movement},
  volume = {123},
  ISSN = {1388-2457},
  url = {http://dx.doi.org/10.1016/j.clinph.2012.01.013},
  DOI = {10.1016/j.clinph.2012.01.013},
  number = {6},
  journal = {Clinical Neurophysiology},
  publisher = {Elsevier BV},
  author = {Solis-Escalante,  Teodoro and M\"{u}ller-Putz,  Gernot R. and Pfurtscheller,  Gert and Neuper,  Christa},
  year = {2012},
  month = jun,
  pages = {1182-1190}
}

@article{Cassim2001,
  title = {Does post-movement beta synchronization reflect an idling motor cortex?},
  url = {https://doi.org/10.1097/00001756-200112040-00051},
  DOI = {10.1097/00001756-200112040-00051},
  journal = {NeuroReport},
  publisher = {Lippincott Williams \& Wilkins},
  author = {Cassim, François and Monaca, Christelle and Szurhaj, William and Bourriez, Jean-Louis and Defebvre, Luc and Derambure, Philippe and Guieu, Jean-Daniel},
  year = {2001},
  month = dec
}

@article{Aron2007,
  title = {Triangulating a Cognitive Control Network Using Diffusion-Weighted Magnetic Resonance Imaging (MRI) and Functional MRI},
  volume = {27},
  ISSN = {1529-2401},
  url = {http://dx.doi.org/10.1523/JNEUROSCI.0519-07.2007},
  DOI = {10.1523/jneurosci.0519-07.2007},
  number = {14},
  journal = {The Journal of Neuroscience},
  publisher = {Society for Neuroscience},
  author = {Aron,  Adam R. and Behrens,  Tim E. and Smith,  Steve and Frank,  Michael J. and Poldrack,  Russell A.},
  year = {2007},
  month = apr,
  pages = {3743-3752}
}
@article{Rae2015,
  title = {The Prefrontal Cortex Achieves Inhibitory Control by Facilitating Subcortical Motor Pathway Connectivity},
  volume = {35},
  ISSN = {1529-2401},
  url = {http://dx.doi.org/10.1523/JNEUROSCI.3093-13.2015},
  DOI = {10.1523/jneurosci.3093-13.2015},
  number = {2},
  journal = {The Journal of Neuroscience},
  publisher = {Society for Neuroscience},
  author = {Rae,  Charlotte L. and Hughes,  Laura E. and Anderson,  Michael C. and Rowe,  James B.},
  year = {2015},
  month = jan,
  pages = {786-794}
}

@article{Jahfari2011,
  title = {Effective Connectivity Reveals Important Roles for Both the Hyperdirect (Fronto-Subthalamic) and the Indirect (Fronto-Striatal-Pallidal) Fronto-Basal Ganglia Pathways during Response Inhibition},
  volume = {31},
  ISSN = {1529-2401},
  url = {http://dx.doi.org/10.1523/JNEUROSCI.5253-10.2011},
  DOI = {10.1523/jneurosci.5253-10.2011},
  number = {18},
  journal = {The Journal of Neuroscience},
  publisher = {Society for Neuroscience},
  author = {Jahfari,  Sara and Waldorp,  Lourens and van den Wildenberg,  Wery P. M. and Scholte,  H. Steven and Ridderinkhof,  K. Richard and Forstmann,  Birte U.},
  year = {2011},
  month = may,
  pages = {6891-6899}
}

@article{Cai2014,
  title = {Dissociable Roles of Right Inferior Frontal Cortex and Anterior Insula in Inhibitory Control: Evidence from Intrinsic and Task-Related Functional Parcellation,  Connectivity,  and Response Profile Analyses across Multiple Datasets},
  volume = {34},
  ISSN = {1529-2401},
  url = {http://dx.doi.org/10.1523/jneurosci.3048-14.2014},
  DOI = {10.1523/jneurosci.3048-14.2014},
  number = {44},
  journal = {The Journal of Neuroscience},
  publisher = {Society for Neuroscience},
  author = {Cai,  Weidong and Ryali,  Srikanth and Chen,  Tianwen and Li,  Chiang-Shan R. and Menon,  Vinod},
  year = {2014},
  month = oct,
  pages = {14652-14667}
}

@article{Limongi2017,
  title = {Successful and unsuccessful response inhibitions differentially affect the effective connectivity between insular,  presupplementary-motor,  and striatal areas.},
  volume = {131},
  ISSN = {0735-7044},
  url = {http://dx.doi.org/10.1037/bne0000175},
  DOI = {10.1037/bne0000175},
  number = {1},
  journal = {Behavioral Neuroscience},
  publisher = {American Psychological Association (APA)},
  author = {Limongi,  Roberto and Pérez,  Francisco J.},
  year = {2017},
  pages = {20-32}
}

@article{Nowak2017,
  title = {Driving Human Motor Cortical Oscillations Leads to Behaviorally Relevant Changes in Local GABAAInhibition: A tACS-TMS Study},
  volume = {37},
  ISSN = {1529-2401},
  url = {http://dx.doi.org/10.1523/JNEUROSCI.0098-17.2017},
  DOI = {10.1523/jneurosci.0098-17.2017},
  number = {17},
  journal = {The Journal of Neuroscience},
  publisher = {Society for Neuroscience},
  author = {Nowak,  Magdalena and Hinson,  Emily and van Ede,  Freek and Pogosyan,  Alek and Guerra,  Andrea and Quinn,  Andrew and Brown,  Peter and Stagg,  Charlotte J.},
  year = {2017},
  month = mar,
  pages = {4481-4492}
}

@article{To2018,
  title = {Changing Brain Networks Through Non-invasive Neuromodulation},
  volume = {12},
  ISSN = {1662-5161},
  url = {http://dx.doi.org/10.3389/fnhum.2018.00128},
  DOI = {10.3389/fnhum.2018.00128},
  journal = {Frontiers in Human Neuroscience},
  publisher = {Frontiers Media SA},
  author = {To,  Wing Ting and De Ridder,  Dirk and Hart Jr.,  John and Vanneste,  Sven},
  year = {2018},
  month = apr 
}
@article{Pariz2023,
  title = {Selective control of synaptic plasticity in heterogeneous networks through transcranial alternating current stimulation (tACS)},
  volume = {19},
  ISSN = {1553-7358},
  url = {http://dx.doi.org/10.1371/journal.pcbi.1010736},
  DOI = {10.1371/journal.pcbi.1010736},
  number = {4},
  journal = {PLOS Computational Biology},
  publisher = {Public Library of Science (PLoS)},
  author = {Pariz,  Aref and Trotter,  Daniel and Hutt,  Axel and Lefebvre,  Jeremie},
  editor = {Bush,  Daniel},
  year = {2023},
  month = apr,
  pages = {e1010736}
}

@article{Violante2017,
  title = {Externally induced frontoparietal synchronization modulates network dynamics and enhances working memory performance},
  volume = {6},
  ISSN = {2050-084X},
  url = {http://dx.doi.org/10.7554/eLife.22001},
  DOI = {10.7554/elife.22001},
  journal = {eLife},
  publisher = {eLife Sciences Publications,  Ltd},
  author = {Violante,  Ines R and Li,  Lucia M and Carmichael,  David W and Lorenz,  Romy and Leech,  Robert and Hampshire,  Adam and Rothwell,  John C and Sharp,  David J},
  year = {2017},
  month = mar 
}

@article{Vosskuhl2018,
  title = {Non-invasive Brain Stimulation: A Paradigm Shift in Understanding Brain Oscillations},
  volume = {12},
  ISSN = {1662-5161},
  url = {http://dx.doi.org/10.3389/fnhum.2018.00211},
  DOI = {10.3389/fnhum.2018.00211},
  journal = {Frontiers in Human Neuroscience},
  publisher = {Frontiers Media SA},
  author = {Vosskuhl,  Johannes and Str\"{u}ber,  Daniel and Herrmann,  Christoph S.},
  year = {2018},
  month = may 
}

@article{AbellanedaPrez2020,
  title = {Differential tDCS and tACS Effects on Working Memory-Related Neural Activity and Resting-State Connectivity},
  volume = {13},
  ISSN = {1662-453X},
  url = {http://dx.doi.org/10.3389/fnins.2019.01440},
  DOI = {10.3389/fnins.2019.01440},
  journal = {Frontiers in Neuroscience},
  publisher = {Frontiers Media SA},
  author = {Abellaneda-Pérez,  Kilian and Vaqué-Alcázar,  Lídia and Perellón-Alfonso,  Ruben and Bargalló,  Núria and Kuo,  Min-Fang and Pascual-Leone,  Alvaro and Nitsche,  Michael A. and Bartrés-Faz,  David},
  year = {2020},
  month = jan 
}

@article{Tan2016,
  title = {Post-Movement Beta Activity in Sensorimotor Cortex Indexes Confidence in the Estimations from Internal Models},
  volume = {36},
  ISSN = {1529-2401},
  url = {http://dx.doi.org/10.1523/JNEUROSCI.3204-15.2016},
  DOI = {10.1523/jneurosci.3204-15.2016},
  number = {5},
  journal = {The Journal of Neuroscience},
  publisher = {Society for Neuroscience},
  author = {Tan,  Huiling and Wade,  Cian and Brown,  Peter},
  year = {2016},
  month = feb,
  pages = {1516-1528}
}

@article{Helfrich2014,
  title = {Entrainment of Brain Oscillations by Transcranial Alternating Current Stimulation},
  volume = {24},
  ISSN = {0960-9822},
  url = {http://dx.doi.org/10.1016/j.cub.2013.12.041},
  DOI = {10.1016/j.cub.2013.12.041},
  number = {3},
  journal = {Current Biology},
  publisher = {Elsevier BV},
  author = {Helfrich,  Randolph F. and Schneider,  Till R. and Rach,  Stefan and Trautmann-Lengsfeld,  Sina A. and Engel,  Andreas K. and Herrmann,  Christoph S.},
  year = {2014},
  month = feb,
  pages = {333-339}
}

@article{Tseng2016,
  title = {The critical role of phase difference in gamma oscillation within the temporoparietal network for binding visual working memory},
  volume = {6},
  ISSN = {2045-2322},
  url = {http://dx.doi.org/10.1038/srep32138},
  DOI = {10.1038/srep32138},
  number = {1},
  journal = {Scientific Reports},
  publisher = {Springer Science and Business Media LLC},
  author = {Tseng,  Philip and Chang,  Yu-Ting and Chang,  Chi-Fu and Liang,  Wei-Kuang and Juan,  Chi-Hung},
  year = {2016},
  month = aug 
}

@article{Haslacher2024,
  title = {Working memory enhancement using real-time phase-tuned transcranial alternating current stimulation},
  volume = {17},
  ISSN = {1935-861X},
  url = {http://dx.doi.org/10.1016/j.brs.2024.07.007},
  DOI = {10.1016/j.brs.2024.07.007},
  number = {4},
  journal = {Brain Stimulation},
  publisher = {Elsevier BV},
  author = {Haslacher,  David and Cavallo,  Alessia and Reber,  Philipp and Kattein,  Anna and Thiele,  Moritz and Nasr,  Khaled and Hashemi,  Kimia and Sokoliuk,  Rodika and Thut,  Gregor and Soekadar,  Surjo R.},
  year = {2024},
  month = jul,
  pages = {850-859}
}

@article{Frhlich2010,
  title = {Endogenous Electric Fields May Guide Neocortical Network Activity},
  volume = {67},
  ISSN = {0896-6273},
  url = {http://dx.doi.org/10.1016/j.neuron.2010.06.005},
  DOI = {10.1016/j.neuron.2010.06.005},
  number = {1},
  journal = {Neuron},
  publisher = {Elsevier BV},
  author = {Fr\"{o}hlich,  Flavio and McCormick,  David A.},
  year = {2010},
  month = jul,
  pages = {129-143}
}

@article{Reato2010,
  title = {Low-Intensity Electrical Stimulation Affects Network Dynamics by Modulating Population Rate and Spike Timing},
  volume = {30},
  ISSN = {1529-2401},
  url = {http://dx.doi.org/10.1523/JNEUROSCI.2059-10.2010},
  DOI = {10.1523/jneurosci.2059-10.2010},
  number = {45},
  journal = {The Journal of Neuroscience},
  publisher = {Society for Neuroscience},
  author = {Reato,  Davide and Rahman,  Asif and Bikson,  Marom and Parra,  Lucas C.},
  year = {2010},
  month = nov,
  pages = {15067-15079}
}

@article{Geffen2025,
  title = {mu-Transcranial Alternating Current Stimulation Induces Phasic Entrainment and Plastic Facilitation of Corticospinal Excitability},
  volume = {61},
  ISSN = {1460-9568},
  url = {http://dx.doi.org/10.1111/ejn.70042},
  DOI = {10.1111/ejn.70042},
  number = {5},
  journal = {European Journal of Neuroscience},
  publisher = {Wiley},
  author = {Geffen,  Asher and Bland,  Nicholas and Sale,  Martin V.},
  year = {2025},
  month = mar 
}
