## Supplementary figures and images for "Closed-loop phase-locked EEG-tACS enables adaptive control of cortical beta synchrony and motor control"

### Fig1_experiment.pdf

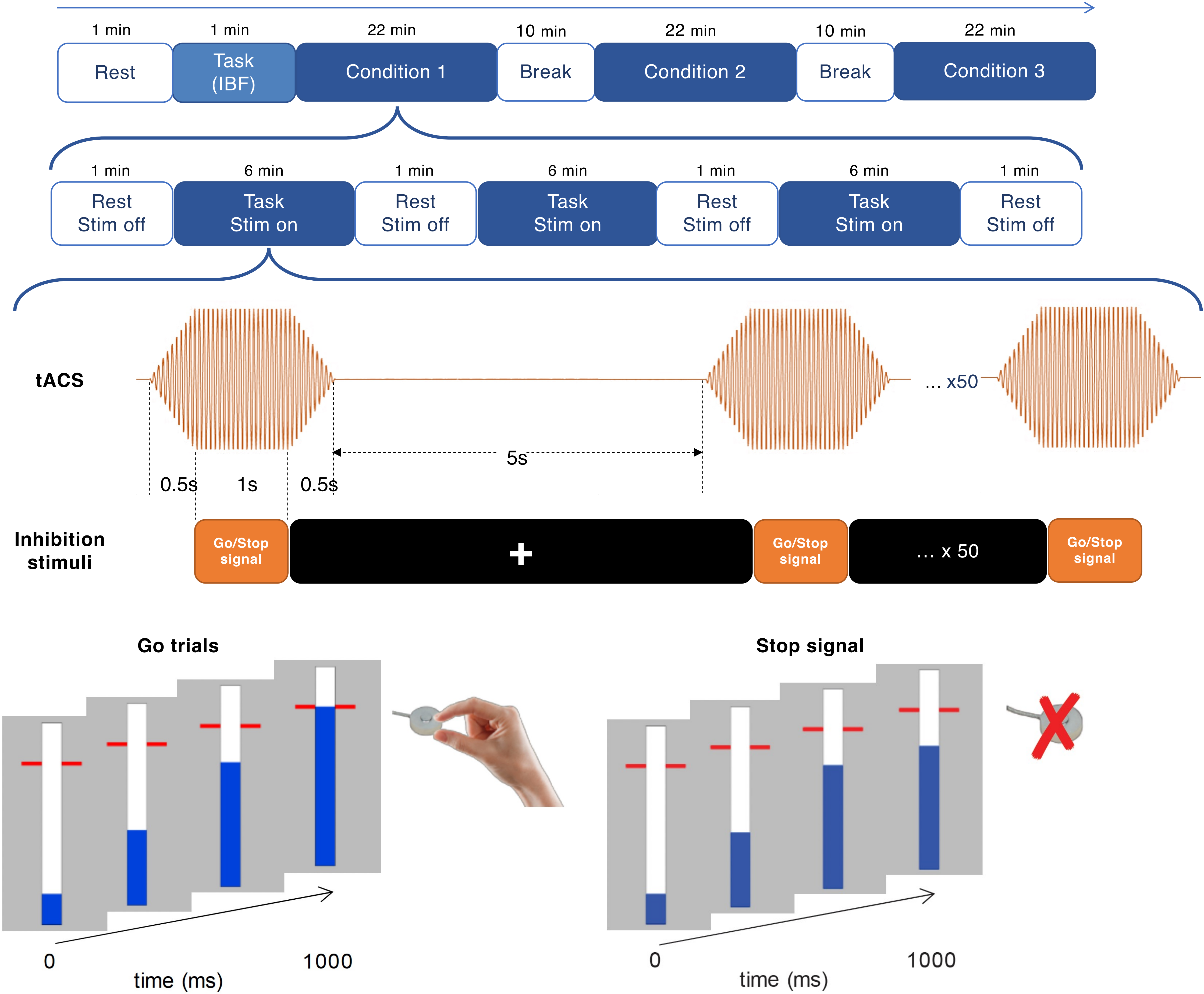

### Fig2_system.pdf

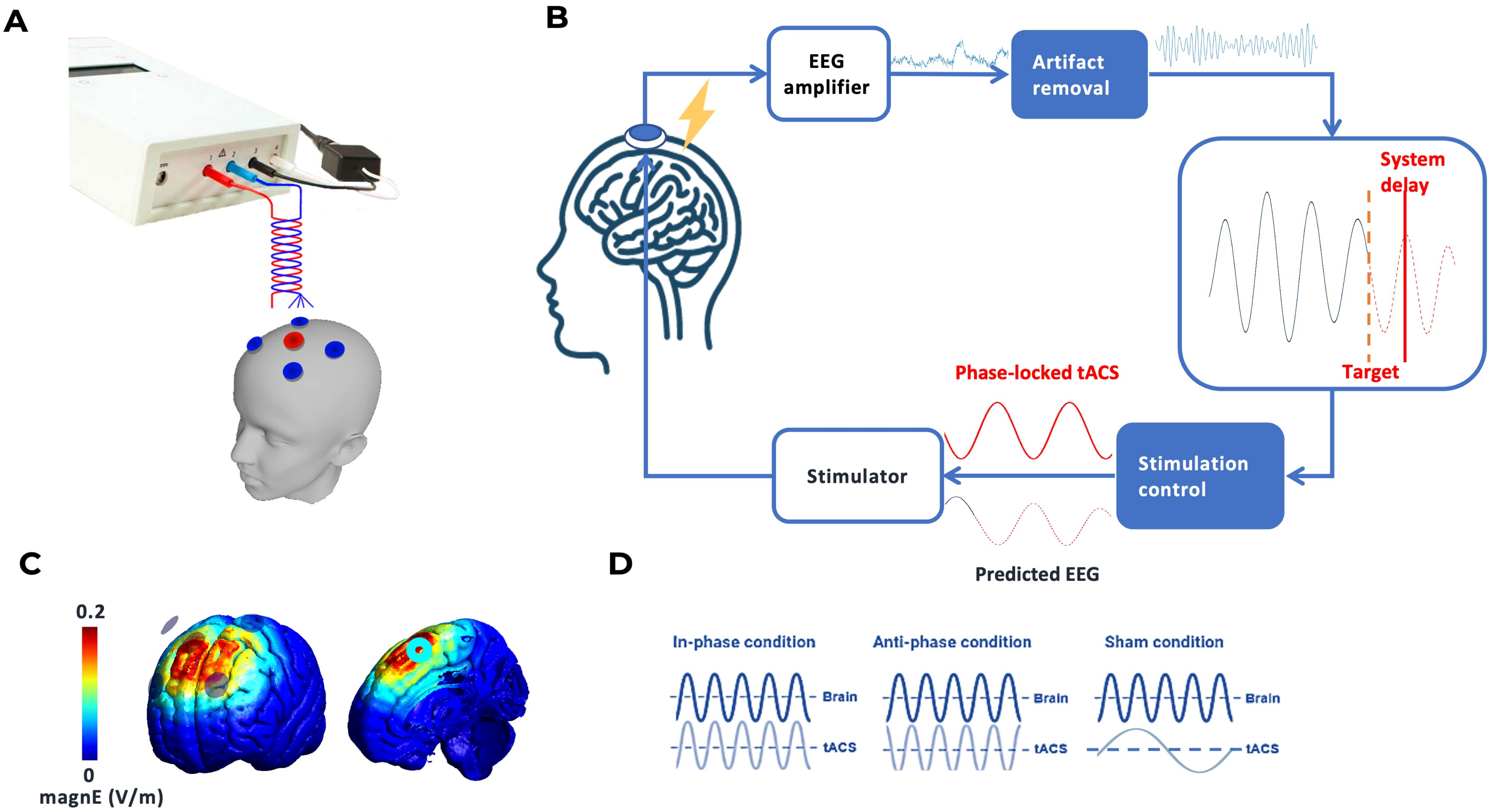

### Fig3_TFR_all.pdf

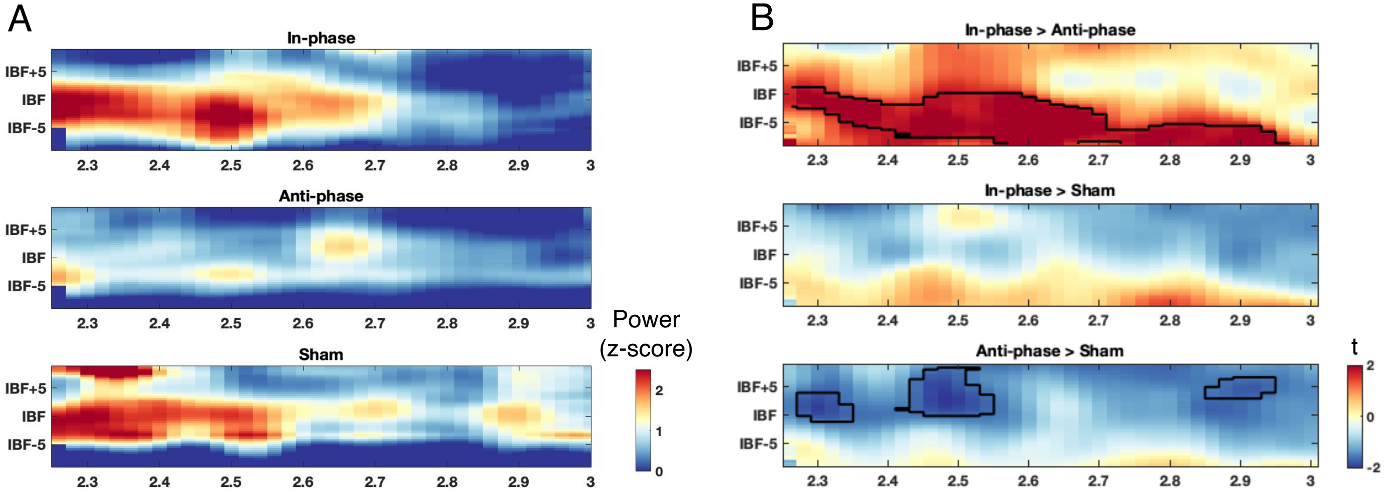

### Fig4_TFR_sham.pdf

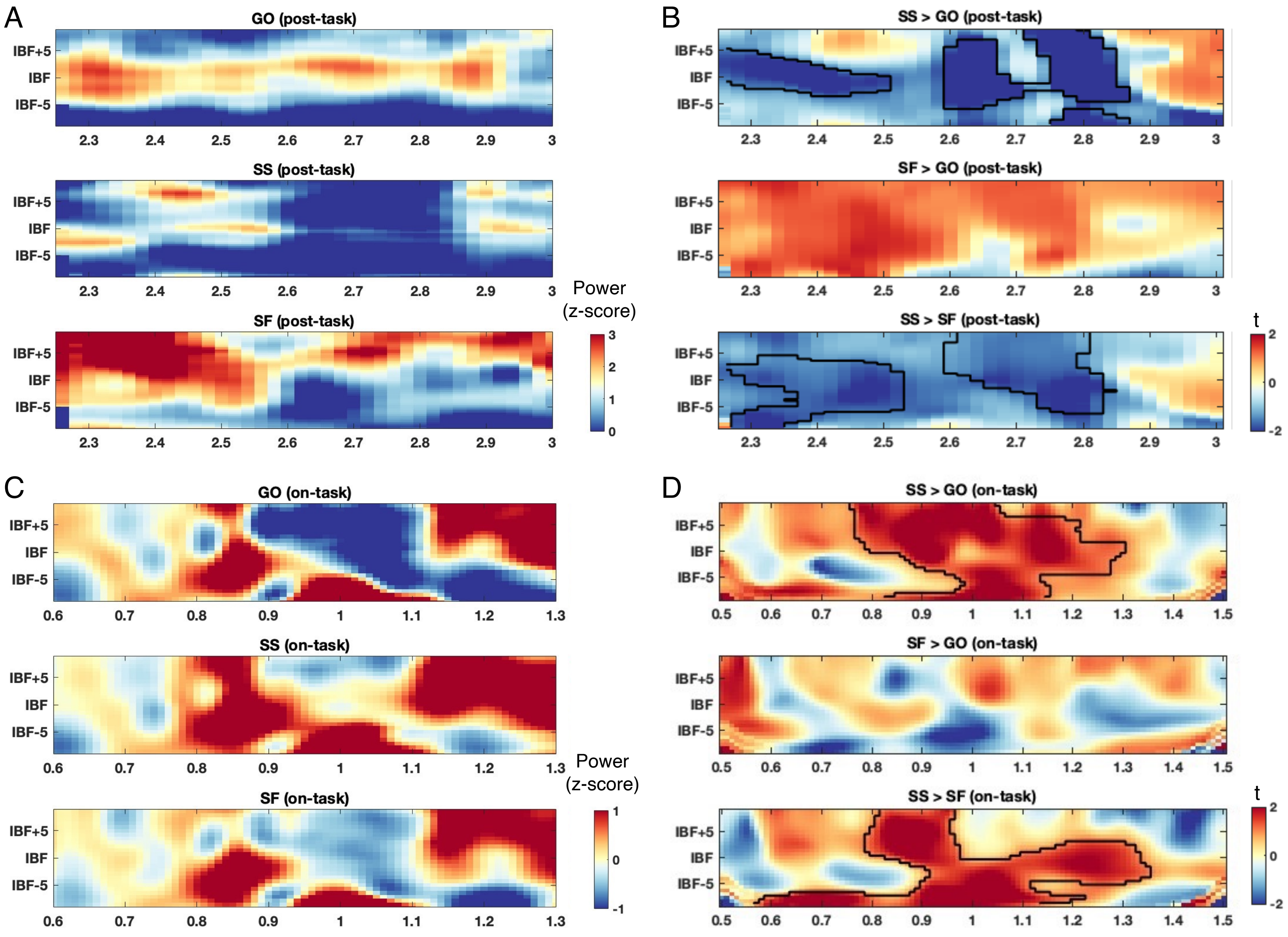

### Fig5_TFR_phase.pdf

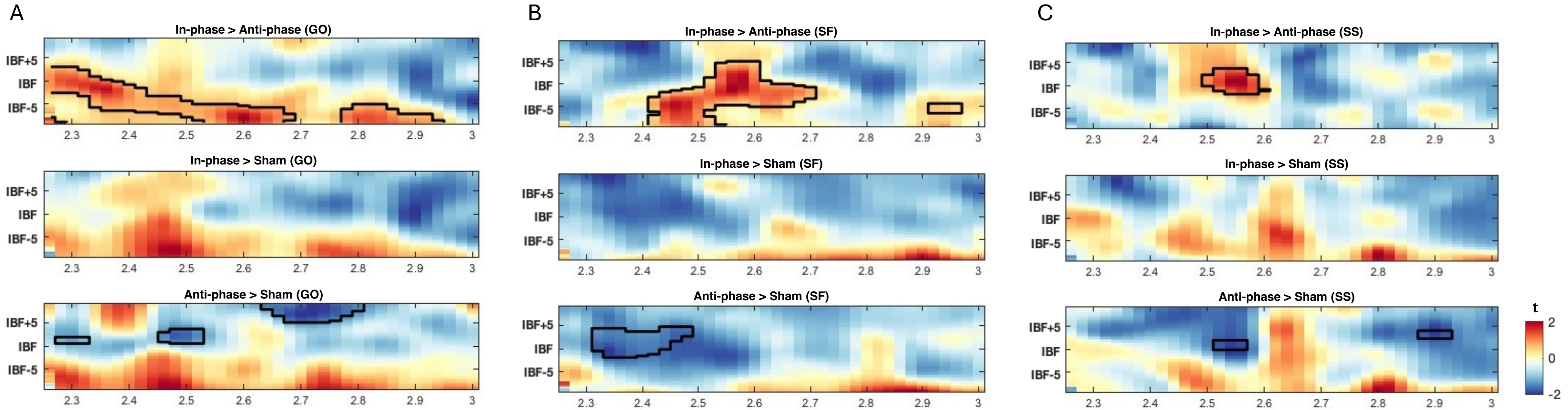

### Fig6_ER.pdf

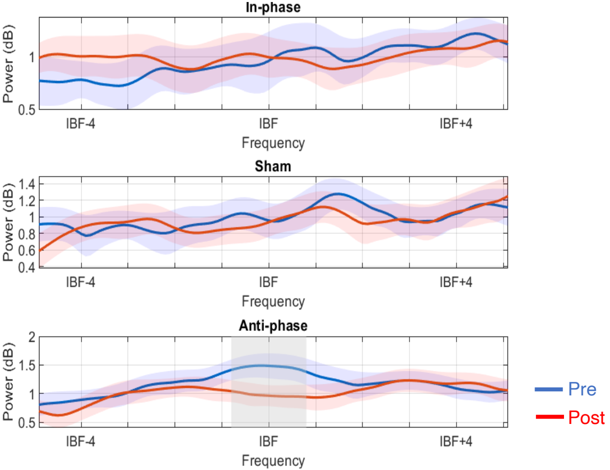

### Fig7_beests.pdf

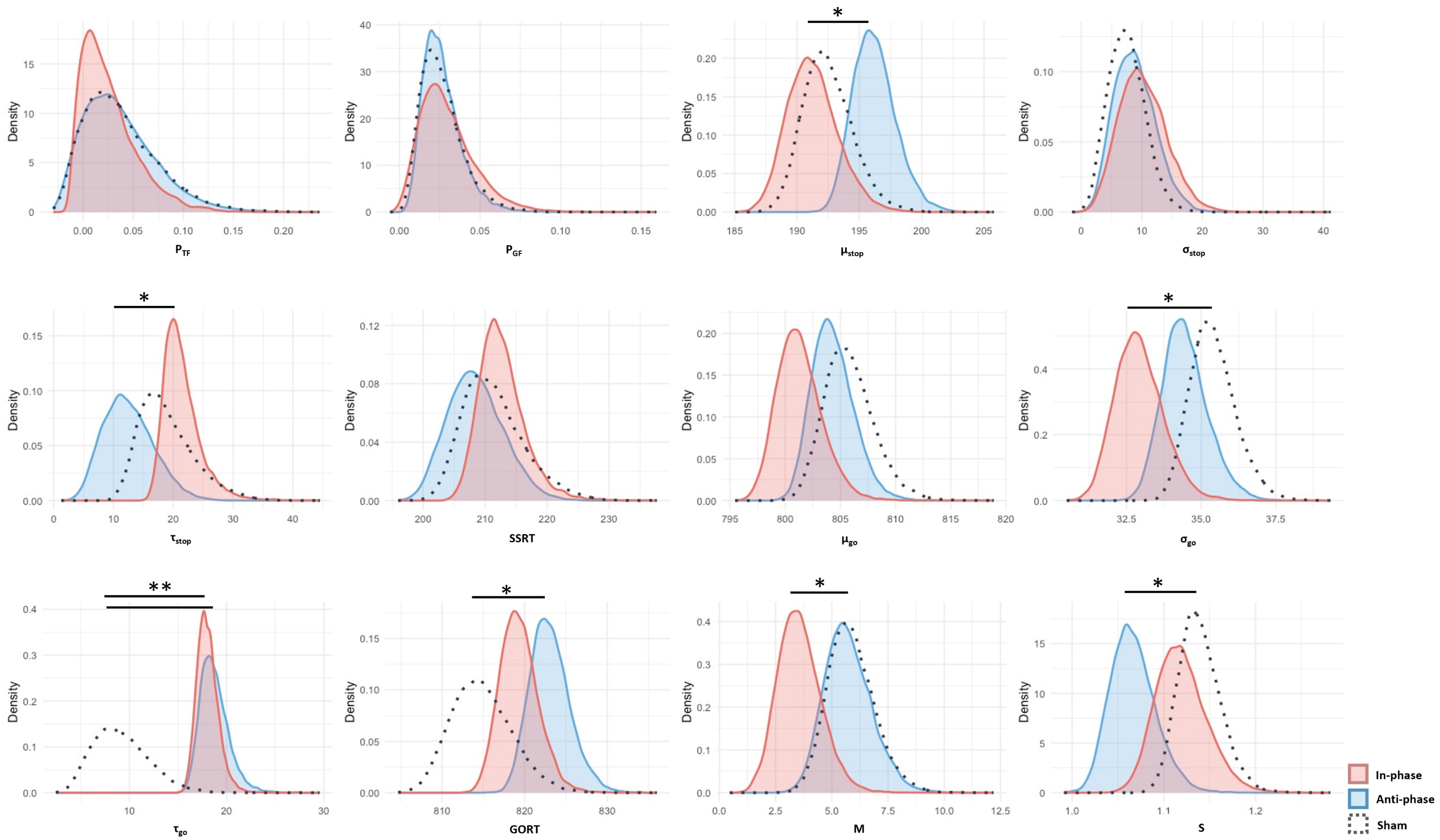

### Fig8_SSRT.pdf

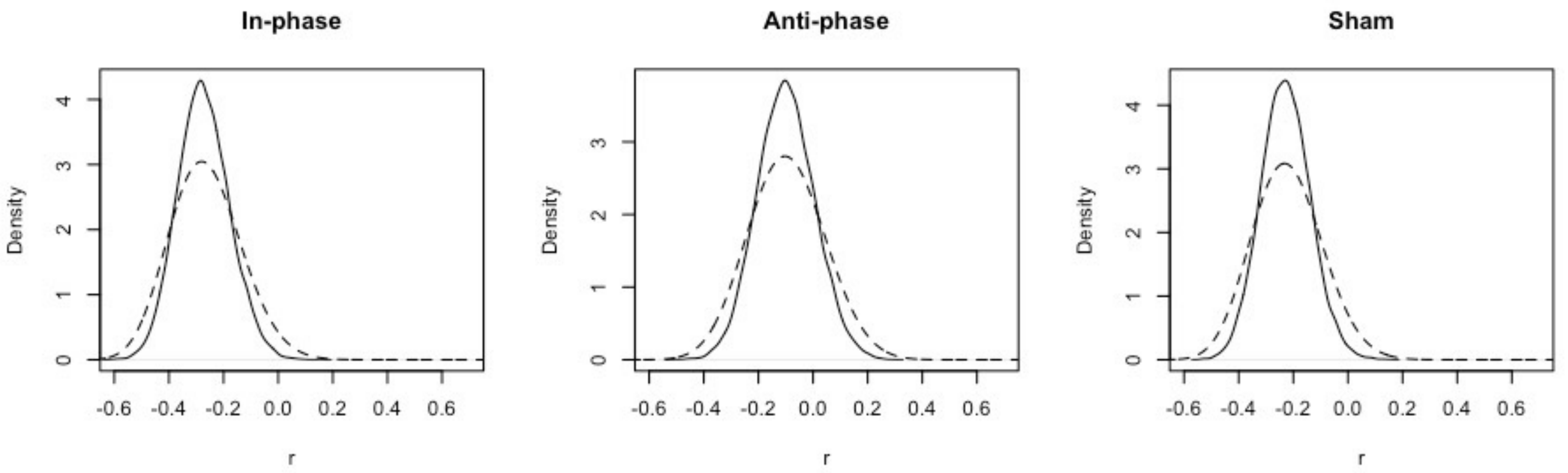

### Fig9_phase.pdf

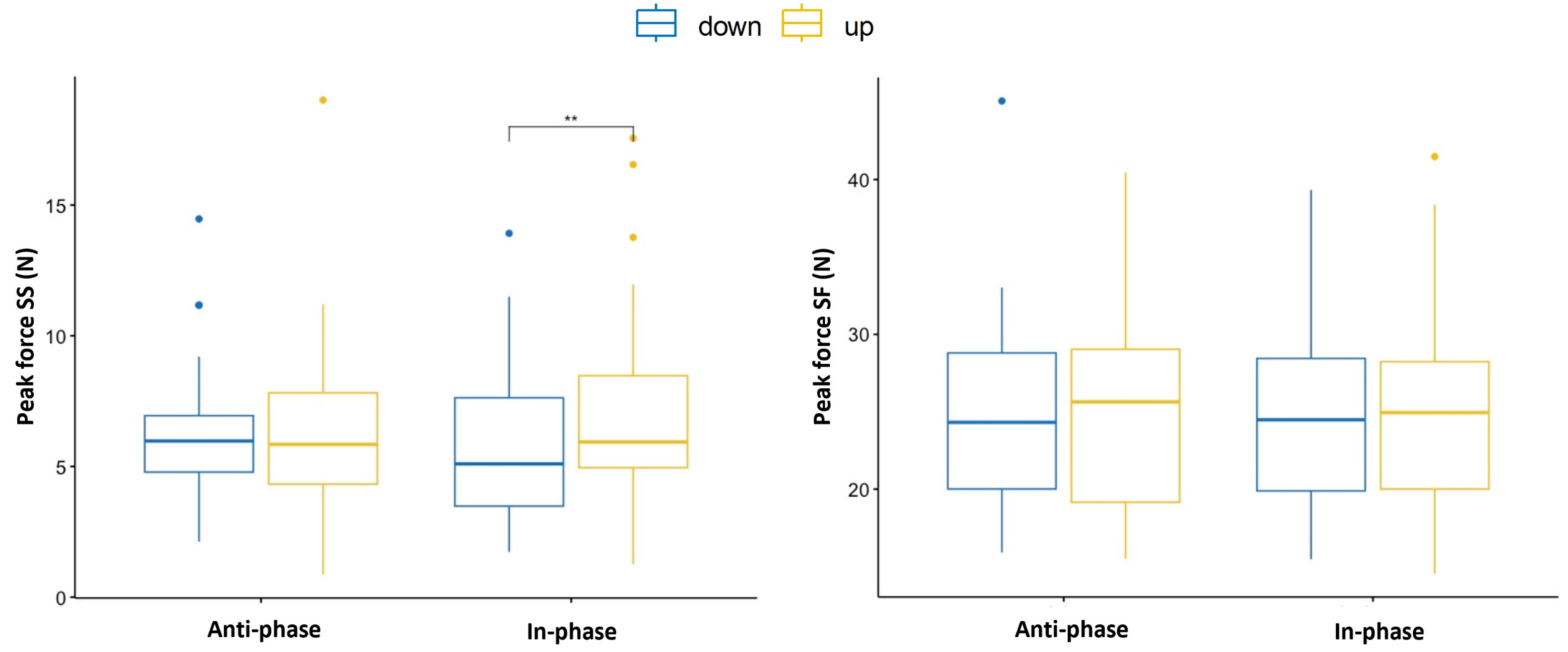

### Figs1_system.pdf

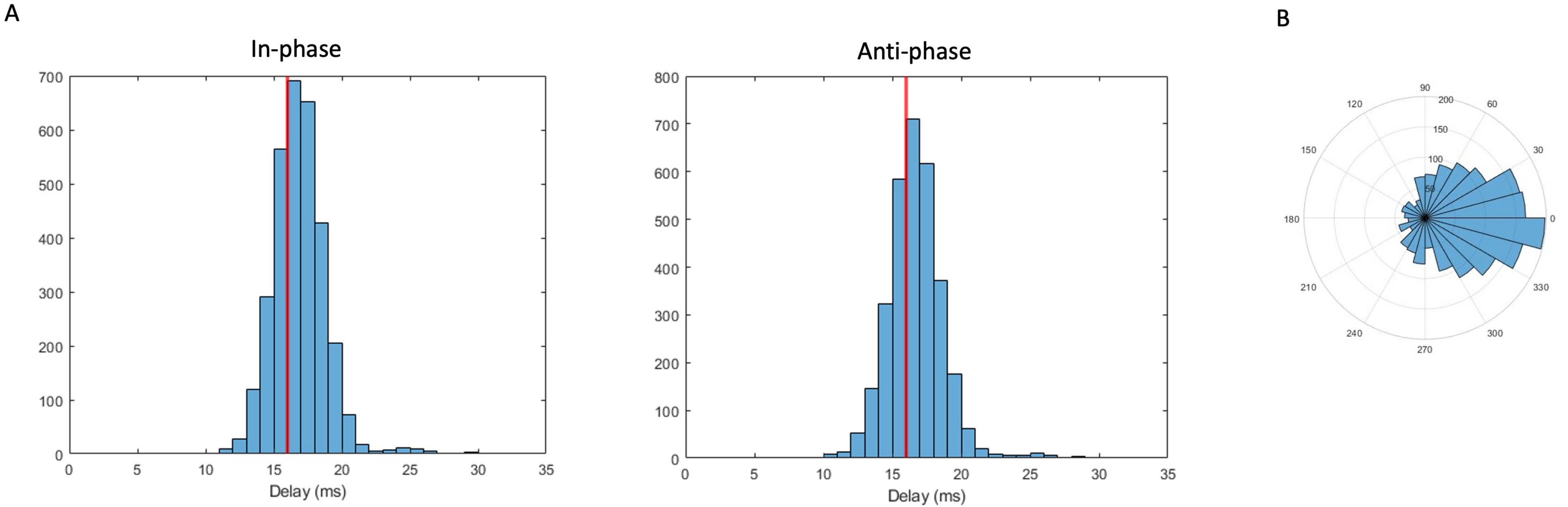

### Figs2_race.pdf

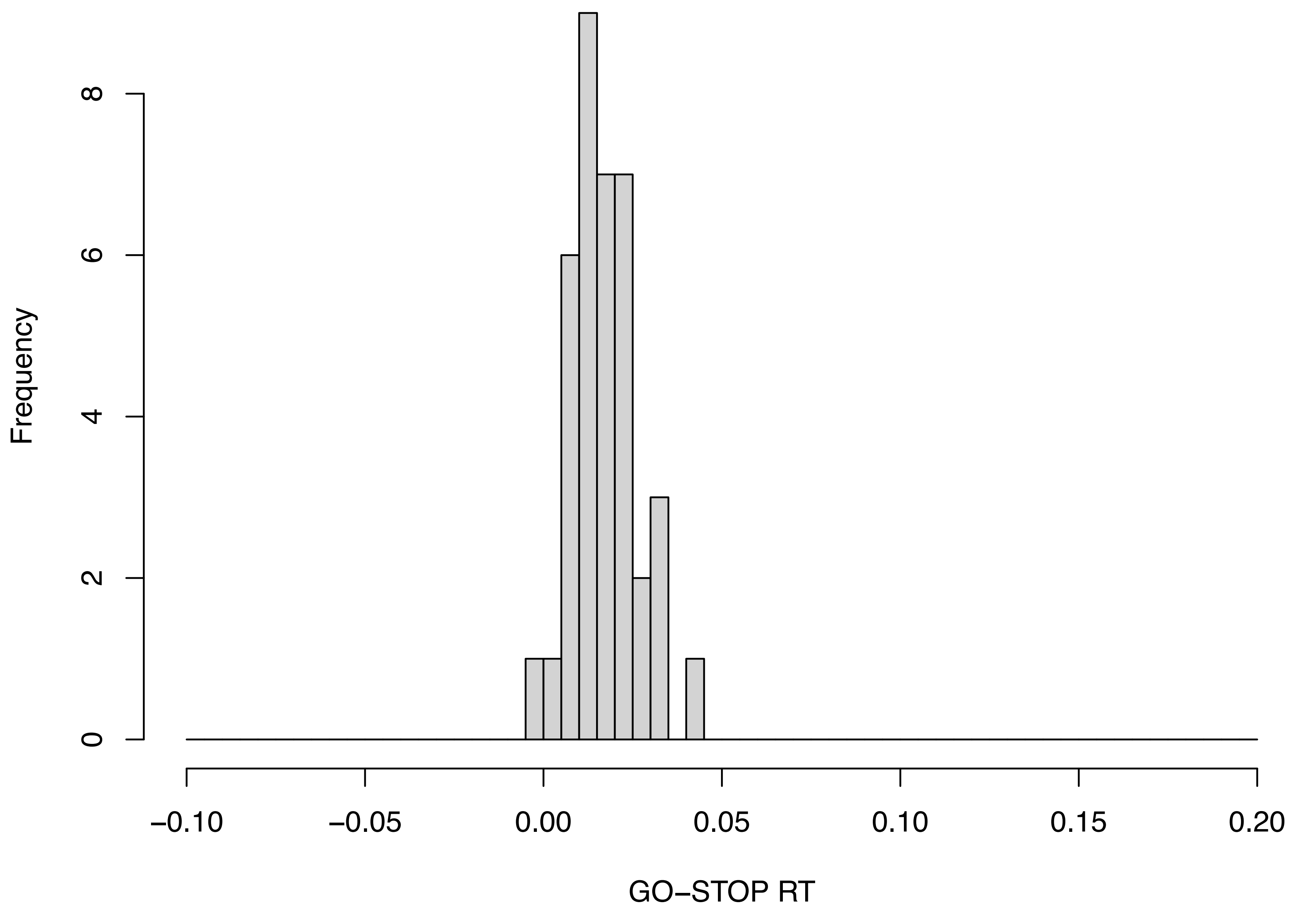

### Figs4_cdf.pdf

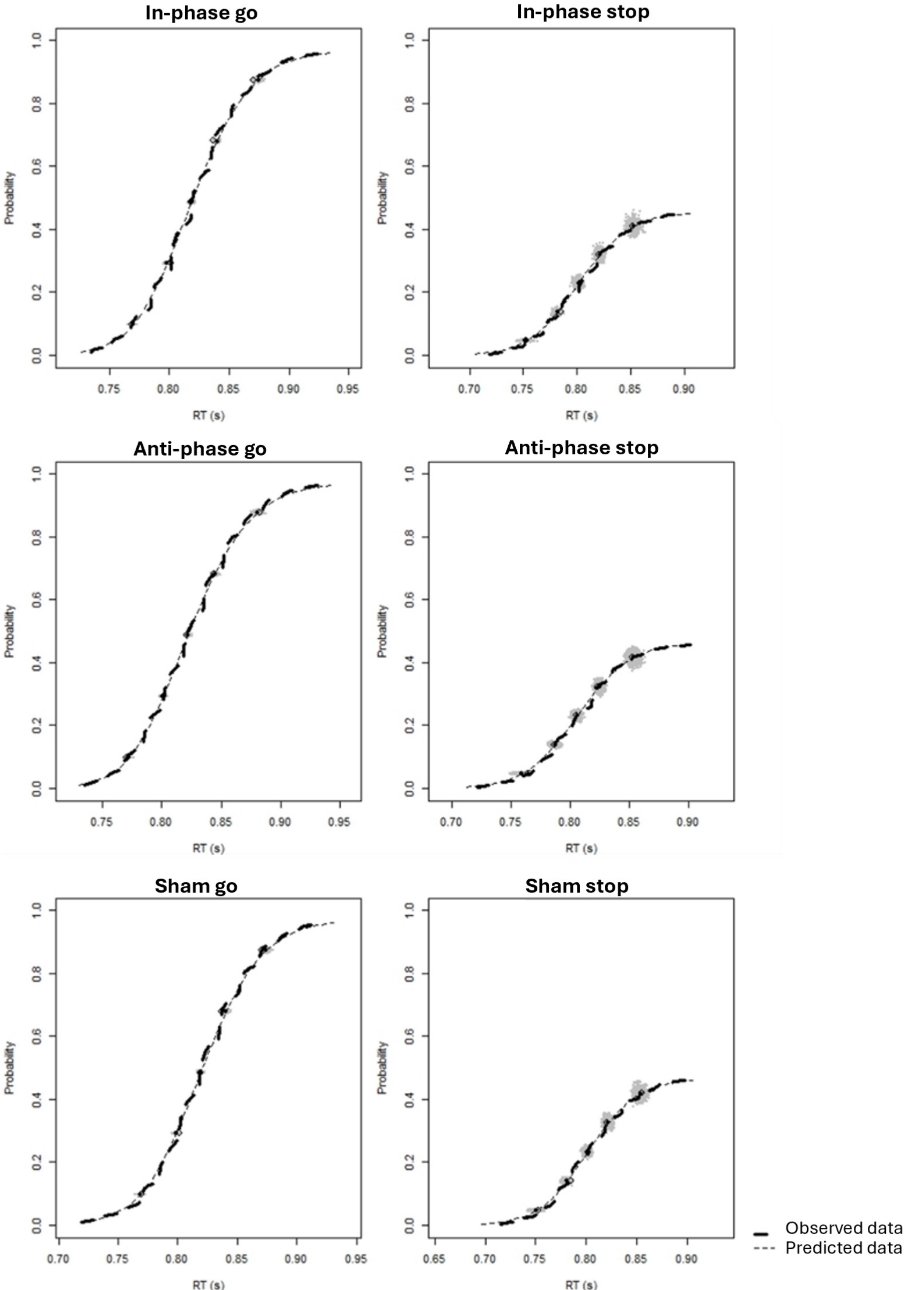

### Figs5_inhibition_function.pdf

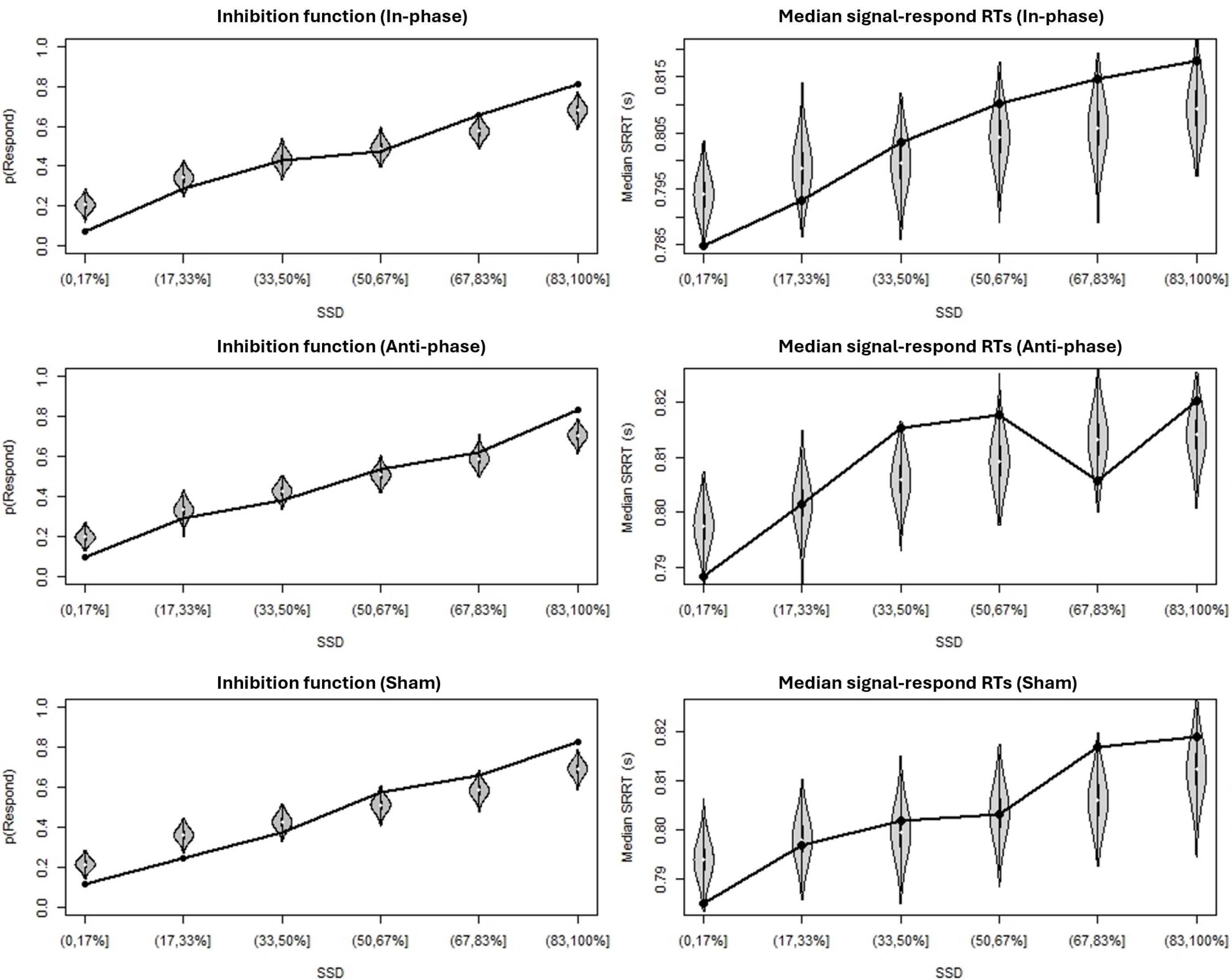

### Figs6_delta.pdf

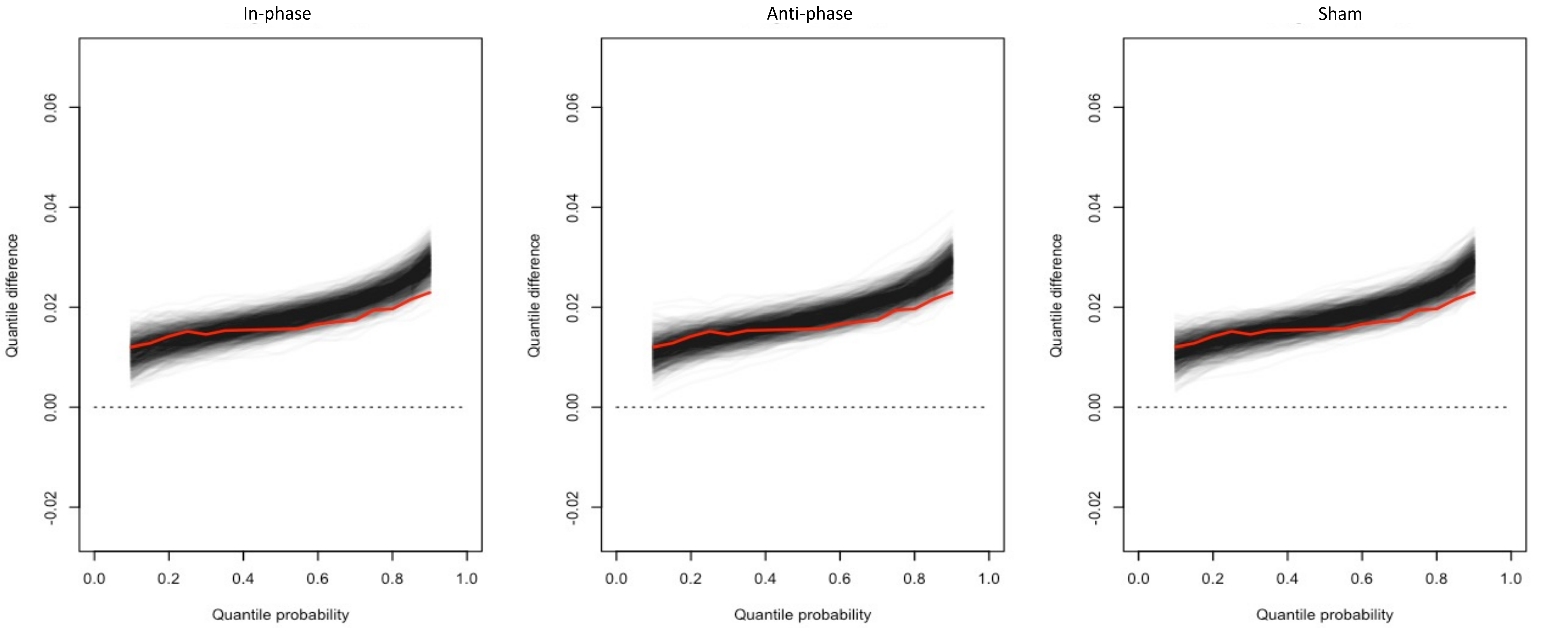
